# Geometric machine learning informed by ground truth: Recovery of conformational continuum from single-particle cryo-EM data of biomolecules

**DOI:** 10.1101/2021.06.18.449029

**Authors:** Evan Seitz, Francisco Acosta-Reyes, Suvrajit Maji, Peter Schwander, Joachim Frank

## Abstract

This work is based on the manifold-embedding approach to study biological molecules exhibiting continuous conformational changes. Previous work established a method capable of reconstructing 3D movies and accompanying energetics of atomic-level structures from single-particle cryo-EM images of macromolecules displaying multiple conformational degrees of freedom. Here, we introduce an unsupervised geometric machine learning approach that is informed by detailed heuristic analysis of manifolds formed by simulated heterogeneous cryo-EM datasets generated from an atomic structure. These simulated data were generated with increasing complexity to account for multiple conformational motions, state occupancies and typical microscope parameters in a wide range of signal-to-noise ratios. Using these datasets as ground-truth, we provide detailed exposition of our findings using several conformational motions while exploring the available parameter space. Guided by these insights, we build a framework to leverage the high-dimensional geometric information obtained towards reconstituting a quasi-continuum of conformational states in the form of a free-energy landscape and respective 3D density maps for all states therein. As shown by a direct comparison of results, this framework offers substantial improvements relative to the previous work.

## INTRODUCTION

Molecular machines, consisting of assemblies of proteins or nucleoproteins, take on a range of unique configurations or *conformational states* as they go through their functional cycles^1^. These states are typically characterized by different spatial constellations of relatively rigid domains, and can be organized in a *state space* according to the continuous motions of each domain along a unique coordinate. Specific sequences of the states in this space form pathways along which the molecular machine may transform. When energetics of states are known, in terms of the machine’s free-energy landscape, a path is singled out along which the machine performs its metabolic function^2^.

A number of recent studies^1,3,4^ were inspired by the realization that it is possible, through the analysis of experimental data, to gain insights into the rules governing a molecular machine’s function. In thermal equilibrium, molecular machines are constantly buffeted by the random motions of nearby solvent molecules, which deform them reversibly as they transition via a series of thermally-driven steps. State-of-the-art single-particle cryo-EM^5–7^ is now capable of providing large numbers of two-dimensional snapshots (i.e., projections) of a molecular machine undergoing this process. When the number of snapshots is sufficiently large – typically several hundred thousand – they capture virtually the entire range of conformations accessible in thermodynamic equilibrium. By virtue of the Boltzmann statistics, the relative number of sightings in each of these states can be translated into changes of free energy^8,9^. Thus, under assumption of thermodynamic equilibrium, the machine’s free-energy landscape can be obtained from an experiment. Accurate estimation of the free-energy landscape for molecular machines and other biological assemblies is of unparalleled importance in modern structural biology.

The way to utilize the data from a single-particle cryo-EM experiment is not easy, however. Ideally, we would wish to compare 3D structures, but only 2D projections are accessible experimentally. After each 2D projection is assigned angles to define its viewing direction on the 2-sphere (*S*^2^), a set of projections in close proximity to one another can be assigned to a unique *projection direction* (PD)^†^. For any PD ⊂ *S*^2^, the challenge is that the relationship among the *N* images therein, represented as a *P* pixel array, require an analysis of the point cloud formed in vector space ℝ^*P*^. Similarities between molecules captured in the same PD, but slightly different conformations, appear as closeness between corresponding points in this high-dimensional space. Thus, for a given PD, images of molecules captured in random states are arranged—by virtue of their similarities—according to the continuous motions of the molecule’s domains.

The geometric structure formed by such an ensemble is an *n*-dimensional manifold Ω embedded in a high-dimensional Euclidean space ℝ^*P*^, with an intrinsic dimension *n* equal to the number of the system’s independent molecular degrees of freedom. By choosing a suitable embedding that maps the data points in Ω into a low-dimensional Euclidean space, we create the foundation for the analysis of the molecule’s conformational spectrum and free-energy landscape by a machine learning approach. In the following, we use the term *PD-manifold approach* to refer to this strategy. Specifically, it entails the grouping of cryo-EM data into individual PDs, with the subset of images within each PD analyzed via manifold embedding, and the resulting representations combined into a consolidated conformational spectrum. This approach was first introduced by Dashti et al.^1^ and is now termed ManifoldEM^10,11^. Results from previous ManifoldEM studies on biological systems—including the ribosome^1^, ryanodine receptor^10^ and SARS-CoV-2 spike protein^12^—have proven its viability and its potential to provide new information on the functional dynamics of molecules.

As manifolds are encountered in many domains of mathematics, science and engineering^13^, the aim of dimensionality reduction has been widely pursued and given rise to a number of well-established techniques to analyze large and complex datasets. Representing data points on Ω in terms of leading eigenvalues and eigen-vectors gives valuable insights into its intrinsic structure, with these relationships having been well studied in the context of spectral geometry^14^. In the analysis of cryo-EM data, both linear^3,15–21^ and nonlinear^1,10,20,22^ dimensionality reduction methods have been applied, primarily principal component analysis^23^ (PCA) and diffusion maps^24,25^ (DM), respectively. Both approaches allow an analysis of the data points in Ω as embedded in ℝ^*N*^, whose entries are the first *N* eigenvectors of the respective graph, and noting that only a leading subset of these are needed for retrieving the conformational spectrum.

In the PCA approach, eigenvectors are obtained from the covariance matrix, whereas DM approximates the eigenfunctions of the Laplace-Beltrami operator (LBO) on Ω, sampled at the given data points. Some techniques are not so easily classified, however, such as the method of Laplacian spectral volumes^4^, which relies on both linear and nonlinear dimensionality reduction. The application of these methods can further be classified based on their type of data input – generating embeddings from either 2D projections straight from a cryo-EM experiment (i.e., the ManifoldEM approach), or 3D density maps which have been reconstructed from those projections^4,26–29^. Regardless of the approach, since the intrinsic structure of such manifolds formed by the data is unknown, these competing reconstruction methods cannot be immediately validated or informatively compared using experimental data alone. They instead require careful evaluation using appropriate synthetic ground-truth datasets.

The purpose of the present study is twofold. For our first endeavor, we provide a heuristic investigation of the manifolds obtained from synthetic quasi-continuous ground-truth datasets, with properties endowed as is anticipated from cryo-EM visualization of biological molecules. To this end, we create several state spaces by simulating a molecule with movable domains, each having undergone a series of independent *conformational motions* (CMs), with the number of CMs constructed for each state space defining the intrinsic dimensionality of the dataset. We then determine how these quasi-continuous motions are reflected in the low-dimensional representations of the manifold’s spectral geometry, obtained by linear or nonlinear dimensionality reduction using kernel methods. For either construction, we derive an explicit expression to account for the geometric structures observed, and describe how to interpret this information as it exists on a hypersurface spanned by multiple degrees of freedom. This heuristic analysis is introduced as a clean slate free from assumptions, aiming to further investigate—using ideal data—the feasibility of manifold embedding techniques under realistic experimental conditions, while exposing any intrinsic uncertainties that may arise. Recently, several issues and limitations have been documented^11^ in the ManifoldEM framework, further amplifying the motivation of our current pursuit.

For our second endeavor, we introduce a novel methodology (which we will term ESPER: “Embedded sub-space partitioning and eigenfunction realignment”) for extraction of conformational information from specific subspaces of PD-manifold (Ω_PD_) embeddings, which we use to generate the molecular machine’s free-energy landscape and corresponding 3D movies depicting its function. Whereas the previous approach^1,10,11^ reconstructs images via nonlinear Laplacian spectral analysis^30^ (NLSA) in an additionally embedded space spanned by one or more CMs, ESPER instead captures each CM directly from the initial embedding while retaining the original cryo-EM images. In addition, several novel operations and refinements to the existing PD-manifold approach are introduced, including a previously unaccounted-for high-dimensional eigenbasis transformation that we deem essential for correctly recapitulating ground-truth information, as well as identification of the proper 2D subspaces required to adequately capture each CM. We demonstrate that this alternative methodology provides conformational movies of significantly improved quality, further enabling the use of efficient strategies for generating multidimensional free-energy landscapes not accessible via the founding ManifoldEM framework. Ultimately, we will show that, when certain requirements are met in the quality and structure of a dataset, ESPER offers an alternative strategy with many benefits compared to the current ManifoldEM approach.

## METHODS

We first introduce a framework for the creation of synthetic ground-truth single-particle cryo-EM datasets in the form of 2D projections of 3D electron density maps arising from a quasi-continuum of atomic structures^31,32^. (Note that in reality cryo-EM data represent projections of the electrostatic or Coulomb potential distribution, which is distinct from the electron density distribution “seen” by X-rays. However, for the present analysis, this distinction is irrelevant). To begin, a suitable macro-molecule is chosen as a foundational model, defined by available structural information in the form of 3D atomic coordinates from the Protein Data Bank^33^ (PDB). Using this initial PDB structure as a seed, a sequence of states is generated by altering the positions of specific domains of the macromolecule’s structure. To mimic quasi-continuous conformational motions, we used equispaced rotations of the domains about their hinge-residue axes. The number of these mutually independent conformational motions^†^ defines the intrinsic dimensionality *n* of the system. By exercising these domain motions independently in all combinations, a set of atomic coordinate structures in PDB-format are generated. In sum, this quasi-continuum of states spans the molecular machine’s state space.

For this work, the heat shock protein Hsp90 was chosen as a starting structure due to its simple design, exhibiting two arm-like domains (chain A and B, containing 677 residues each) connected together in an overarching V-shape^34^. *In vivo*, these arms are known to close after binding of the molecule with ATP, with Hsp90 acting as a chaperone to stabilize the structures of surrounding heat-vulnerable proteins. During its work cycle, Hsp90 naturally undergoes large conformational changes, transitioning from its two arms spread open in a full V-shape (inactive state) to both arms bound together along the protein’s central line of two-fold symmetry (active state) following ATP binding. We initiated our workflow with the fully closed state via entry PDB 2CG9, whose structure was determined at 3.1 Å by X-ray crystallography^35^.

Casting Hsp90’s biological context aside, liberties were taken in the choice of the synthetic model’s leading degrees of freedom. Instead of a single conformational motion (arms open to closed, as *in vivo*), we decided to create three easily-identifiable and fully-decoupled domain motions, which we refer to as CM_1_, CM_2_ and CM_3_. Each CM was designed to cover a unique range of motions, with the cascade of overlaid states making up CM_1_ occupying the largest spatial region, followed in magnitude by CM_2_ and then by CM_3_. Using combinations of these CMs, three synthetic state spaces were generated, with intrinsic dimensionalities of *n* = 1, 2, 3. This was achieved by changing the positions of the first, the first two, or all three regions defined as rigid domains in their given ranges monotonically and (in the latter two cases) independently. Importantly, in view of a later discussion of boundary conditions, there is no steric hindrance between domains within the ranges of their motions.

In the following analysis, these state spaces are termed SS_1_, SS_2_ and SS_3_, and defined by: (1) 20 states exhibiting one degree of freedom (CM_1_); (2) 400 states (20 × 20) with two degrees of freedom (CM_1_, CM_2_); and (3) 1000 states (10 10 10) with three degrees of freedom (CM_1_, CM_2_, CM_3_), respectively. As a specific example of the ranges of motion present in SS_3_, the Root-Mean-Square Deviation (RMSD) was calculated^36^ for the differences of the atomic coordinates between neighboring states in each CM, yielding the values of 1.8 Å, 1.3 Å and 0.3 Å along CM_1_, CM_2_ and CM_3_, respectively; with the RMSD between the first and last state of each CM (representing its total span) yielding 15.3 Å, 11.3 Å and 2.4 Å. Altogether, the total spans of these synthetically-constructed CMs cover a wide range of motions, as one might observe in experiment. In-depth details for these datasets, such as exact atomic descriptions of each state, are provided in the supplementary material (SM) section SM-I^‡^. This section should also be consulted for its description of the indexing used for ordering images within each state space, which is essential for interpreting color maps in figures of embedded manifolds throughout this paper.

Our presentation showcases results from detailed evaluation of three data types—termed data-type I, II, and III—with each step incorporating image artifacts and ensemble statistics in our state-space models as is anticipated in a cryo-EM experiment. Detailed information pertaining to the construction of each of these three data types is provided in the supplementary material. We first investigate the pristine *data-type I*, which is given no simulated experimental artifacts or occupancy assignments. Within this construction, for five example PDs, we examine the manifolds (Ω_PD_) corresponding to each of the three state spaces as obtained via the DM framework, followed by a comparison with those obtained via PCA. Using the eigenfunctions of the LBO, we analytically quantify the trajectories of our simulated conformational changes as embodied by the spectral geometry of each Ω_PD_.

Next, in establishing *data-type II*, we vary the abundance of images per state in each dataset and add noise to the images with varying signal-to-noise ratio (SNR), so as to investigate the influence of statistical coverage on spectral geometry, and to quantify the robustness of this geometry in the presence of noise. Following this analysis, we further increase the presence of experimental artifacts through application of a contrast transfer function (CTF) with realistic microscopy parameters and random defocus variations (within the typical range expected in the experiment), and apply noise to obtain an experimentally-relevant SNR (*data-type III*). Based on the findings of our analysis, we finally introduce an overview of the ESPER method for reconstructing the conformational motions in the form of 2D and 3D movies as obtained from a collection of PDs along a great circle on *S*^2^, with occupancies assigned and transformed into a free-energy landscape (*final analysis*). All Python scripts for reproducing this workflow, including extensive documentation therein, have been made available online^32,37^.

## RESULTS

### I. Diffusion Maps

#### A. Data-type I in State Space 1

For its illustrative qualities, we first analyze embeddings constructed from the eigenvectors obtained from the DM framework for SS_1_, representing one degree of freedom sampled with 20 states in CM_1_. As the workflow for obtaining the general diffusion map for a given dataset has been described in several publications^24,25,38^, we summarize these procedures here while providing detailed information in the supplementary material (see section SM-X, where we define characteristic parameters such as the Gaussian bandwidth, *ε*). Using the DM framework, we ultimately generate a different embedding for each of the five PD manifolds, with each of the resultant point clouds containing 20 points, and each point therein corresponding to an image of a conformational state from CM_1_.

Upon inspection of each embedding (one per PD) for suitable *ε* within the range discovered, we found that the corresponding eigenvalue spectrum for each Ω_PD_ showed a staggered falloff in the significance of leading eigen-vectors, which decayed slowly to zero. Projecting the resultant set of eigenvectors (forming an *eigenbasis*) onto the leading eigenvector (Ψ_1_) alone presented a skewed version of the anticipated mapping, with the indices of states appearing in jumbled sequence near the boundaries. When we alternatively projected the data onto the first two eigenvectors {Ψ_1_, Ψ_2_}—forming a 2D eigen-vector subspace—the conformational signal followed a parabolic trajectory (as shown in the first subplot of Fig. 1-B), confirmed by the proper ordering of indices of points along this curve.

**FIG. 1:**
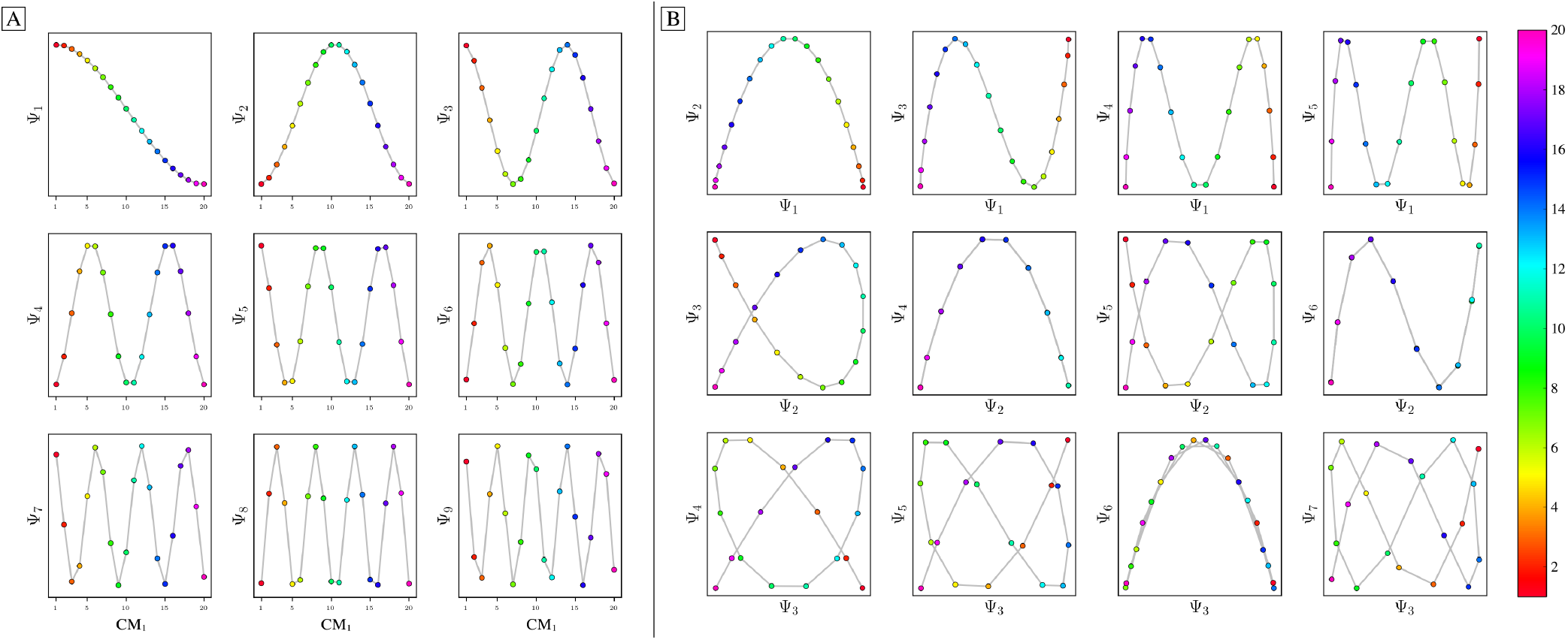
Analysis of eigenfunctions for PD_1_ in SS_1_ (i.e., 20 states total making up one degree of freedom). On the left [A] are the sinusoidal forms {cos(*kπx*) | *k* ∈ ℤ^+^ ≤ *N*} of each eigenvector Ψ_*k*_ that emerge when points (corresponding to images) in each Ψ_*k*_ are ordered precisely in the sequence in which their ground-truth images were constructed. Regardless of any knowledge of such a sequence, the composites of these eigenvectors will always form well-defined geometries (via the Lissajous curves), as shown in [B]. In the first row are the Chebyshev polynomials of the first kind, of which the parabola {Ψ_1_, Ψ_2_} is the simplest mapping of the conformational information present. As is explained below, {Ψ_2_, Ψ_4_} and {Ψ_3_, Ψ_6_} represent *parabolic harmonics* of the {Ψ_1_, Ψ_2_} parabola, which obfuscate the CM content. Finally, note the *nonuniform rates of change* along each Lissajous curve – where it can be seen, for example, that points along the {Ψ_1_, Ψ_2_} parabola are most densely packed near the boundaries and vertex.

Following this analysis, we next proceeded to investigate all other unique 2D combinations of eigenvectors. Mathematically, each such mapping to a 2D vector sub-space is the restriction to the *N*-dimensional embedding of the projection of ℝ^*N*^ onto ℝ^2^; given by {Ψ_1_, Ψ_2_,..., Ψ_*N*_} ↦ {Ψ_*i*_, Ψ_*j*_}, where *i < j*. (For expediency, we will use the term *subspace* to specifically refer to a subspace of an embedded manifold). As seen in Fig. 1-B, a subset of the canonical Lissajous curves^39^ emerged across the 2D subspaces of each Ω_PD_, with the curves in this set having the form

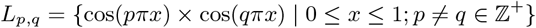

where the operator × denotes the Cartesian product^40^.

The appearance of these *L*_*p,q*_ curves—which are the composite of sinusoids—aligns with known attributes of the Laplace-Beltrami operator. Specifically, the functions *Ψ*_*k*_ = {cos(*kπx*) | 0 ≤ *x* ≤ 1; *k* ∈ ℤ^+^} are the canonical eigenfunctions of the LBO on the interval [0, 1] subject to Neumann boundary conditions^41^, with a metric tensor *g* equal to identity (see section SM-XIII and SM-XIV). By relying on our privileged knowledge of the ground-truth sequential ordering of CM states, we were able to further investigate these underlying sinusoidal forms. For demonstration, we plot each of the 1D points in a given eigenvector as a function of a uniform index *I* ∈ [1, 20] (for the 20 total states in SS_1_), making sure that the ordering of the points in 1D follows the sequence assigned by the ground-truth index of its corresponding image along CM_1_. As seen in Fig. 1-A, when the collection of points in Ψ_*k*_ are ordered appropriately, the eigenfunction’s sinusoidal form emerges along the full extent of the degree of freedom present (i.e., *I* ∈ [1, 20] ↦ *x* ∈ [0, 1]).

Of course, as the points in an experimental dataset naturally arrive in unordered sequence, one would have to properly sort the image indices to recognize these sinusoids; here, for example, there are 20! sequences to consider. In the application, even if an approximation of this sequence were obtained, then in the presence of duplicate CM states (which we anticipate in an experiment), each sinusoid would be irregularly stretched along the *x*-axis where those duplicate states occurred, forming an unwieldy distorted sinusoidal form. However, as the points in each Ψ_*k*_ are always scrambled in the same order in all eigenvectors, the composite of any two will always exist in a readily identifiable *L*_*p,q*_ form. For these composites, we found that CM information is portrayed most simply (without overlap) along a specific subset of *L*, here as seen across the set of 2D subspaces defined in pairwise combination with the leading eigenvector; i.e., ({Ψ_1_, Ψ_2_}, {Ψ_1_, Ψ_3_},..., {Ψ_1_, Ψ_*g*_}), where *g* is the index of the smallest non-zero eigenvalue. Specifically, this subset *T*_*k*_ ∈ *L* corresponds to the known Chebyshev polynomials of the first kind^42^, of which we observed that the parabolic form is the lowest-order member present in each Ω_PD_ embedding.

Given their significance, these 2D subspaces have several important properties worth highlighting for their eventual use (or avoidance). First, note that for each sinusoidal subplot in Fig. 1-A, points are equispaced along the *x*-axis while maintaining the proper sinusoidal form on the *y*-axis, in correspondence with the uniform rotations of the corresponding atomic-coordinate structures. However, due to the Cartesian product, only non-uniform spatial relationships exist between neighboring states in each *L*_*p,q*_. Analytically, this relationship is described by a non-isometric mapping, where lengths in the domain *X*_*a*_ are not preserved in the codomain *X* = Π_*a*∈*A*_*X*_*a*_, and naturally arises when taking a set (indexed via *A*) of Cartesian products (Π) operating on cosine functions *X*_a_ = {cos(*kπx*) | *x* ∈ [0, 1]; *k* ∈ ℤ^+^} that are each uniformly occupied with a finite number o f datapoints. As shown in Fig. 2-A, the spacing between points in *L*_1,2_, which is the composite of two such sinusoids, has an intrinsically nonuniform spatial distribution, with the density of points similarly arranged as seen in the corresponding point clouds.

**FIG. 2:**
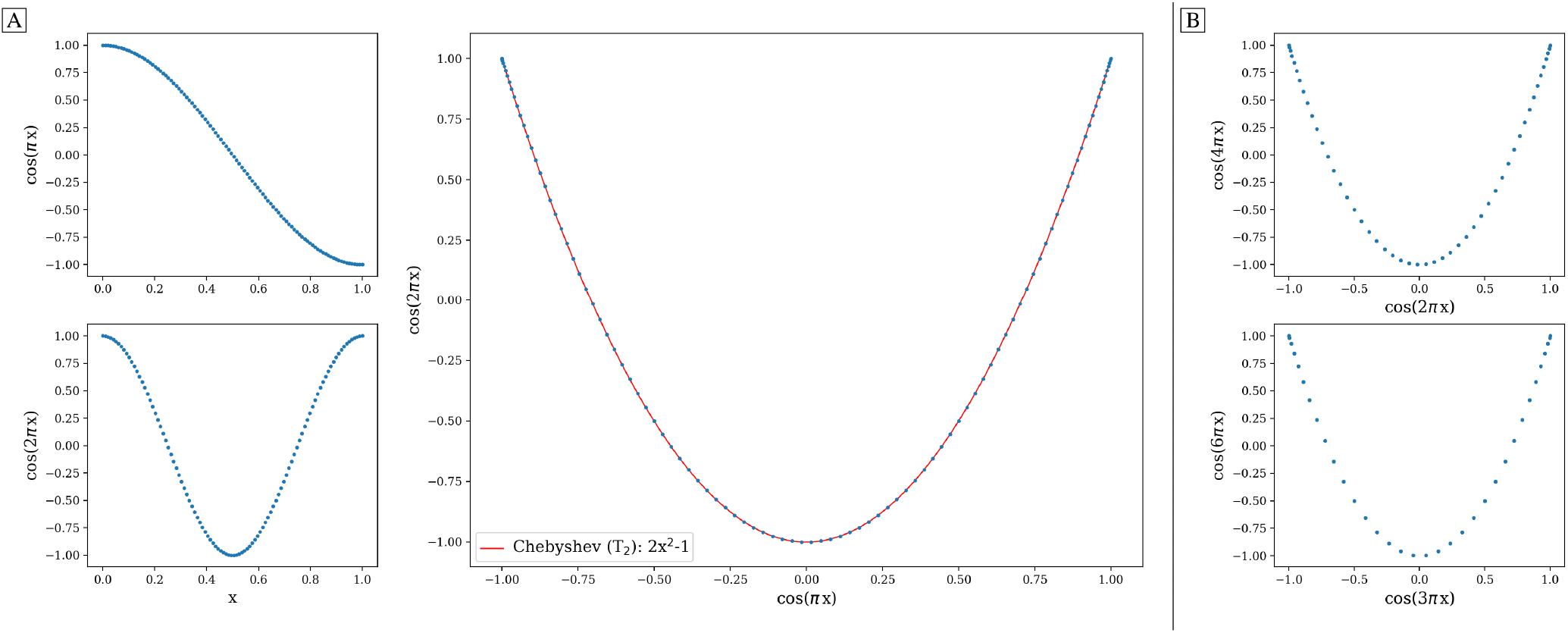
The analytical generation of the Lissajous curve *L*_1,2_ = {cos(*πx*) × cos(2*πx*) | uniform *x* ∈ [0, 1]}, where *L*_1,2_ ∈ ([−1, 1]×[−1, 1]) is shown in [A]. Note the naturally-induced nonuniform spacing between points near the boundaries and vertex of the parabola. As a simple demonstration, we also fit this curve with the Chebyshev *T*_2_ polynomial, which is a subset of the Lissajous curves; however, *T*_2_ does not share the same nonuniformity in spacing as *L*_1,2_. In [B], the parabolic harmonics are likewise generated for *L*_2,4_ and *L*_3,6_. While the same *x*-coordinates were used to generate all underlying cosines for parabolas in both [A] and [B], more than one point in the domain ends up mapping to each coordinate of these parabolic harmonics. As such, these harmonics obfuscate the true conformational information, which is intact on *L*_1,2_.

We denote this aspect with the term *nonuniform rates of change*. As a potential remedy, we investigated the use of an inverse-cosine mapping on each eigenfunction. Fig. S4 provides the results of this transformation on both (1) the analytically-derived cosine functions *k* ∈ {1, 2} and (2) the SS_1_ eigenfunctions obtained by applying DM on images in PD_1_. The first two subplots in Fig. S4 further highlight the remarkable fidelity of the DM eigenfunctions of the graph Laplacian to the analytical form of the LBO, while the third subplot illustrates the results of inverse-cosine transformation. As can be seen, this mapping presents the coordinates of each eigen-function in a space with uniform rates of change, consistent with the ground-truth relationships between atomic-coordinate structures. We will leverage this aspect later in our framework, and indicate any eigenvectors Ψ_*i*_ under this transformation with the insignia Φ_*i*_.

Next for consideration, as seen in Fig. 1-B, there exist several parabolic trajectories scattered throughout the 2D subspaces of a given Ω_PD_. As confirmed by the indices of points and the corresponding color map along each curve, only the first of these parabolas describes the full extent of the conformational motion present monotonically, while all other trailing parabolas display a non-monotonic signal. As a specific example, Fig. 2 shows that the first three such parabolas can be generated via *L*_1,2_, *L*_2,4_ and *L*_3,6_ – which repeat the conformational information once, twice, and three times, respectively, within one span of the parabolic trajectory.

As a consequence, only the mapping from the sinusoids to the first parabola in this set is bijective (injective and surjective)^40^, with all other mappings to higher-order parabolas non-injective surjections. Importantly, since the Cartesian product of continuous functions is continuous and projections from product spaces are also continuous, this bijection further meets the requirements of a *homeomorphism*: a bijective correspondence that preserves the topological structures involved^40^. We denote the higher-order parabolas (formed via the non-injective surjections) as *parabolic harmonics*, which do not preserve topological structure and must be avoided when mapping a given CM; a problem that becomes more challenging in the following subsections as more degrees of freedom are added to the system.

We next compared these sets of 2D subspaces among the five PDs, and found only subtle differences in the distribution of their point clouds. It is important to underscore here the natural discrepancies between each Ω_PD_ that should be expected, which will continue to manifest in several significant forms throughout this analysis. Naturally, as each 2D projection provides an incomplete representation of the underlying 3D density map, depending on the type of motion and its component along the PD under investigation, ground truth is preserved to different degrees. Going forward, we will refer back to this notion under the label *PD disparity*. This disparity affects all Ω_PD_ characteristics, and will become more relevant as we investigate the embeddings of datasets with multiple degrees of freedom.

#### B. Data-type I in State Space 2

To further understand these conformational-variation signals with increasing intrinsic dimensionality *n*, we next investigated the embeddings generated for SS_2_. As seen in Fig. 3-A, by plotting the points in each eigenvector in the specific ground-truth sequence constructed for CM_1_ against a uniform index (now for the 400 states in SS_2_; i.e., {1, 2, 3*,...,* 400}), a similar but now interspersed pattern of sinusoids appeared. Specifically, the appearance of the sinusoids (with increasing *k* ∈ ℤ^+^) only manifested in a subset of all eigen-vectors present, while for all other eigenvectors outside of this set, more arcane patterns emerged. Following this observation, we next reordered the indices of points within all eigenvectors to instead correspond with the specific ground-truth sequence constructed for CM_2_ (i.e., {1, 21, 41*,...,* 381} *,...,* {20, 40, 60*,...,* 400}). The output of this operation can be seen in Fig. 3-B, which manifested a new subset of interspersed sinusoids, with increasing *k′* ∈ ℤ^+^ independent from the previous subset; and inhabiting only those eigenvectors in the complement of the CM_1_ subset. By induction—based on these observations—we conclude that for *n* degrees of freedom in a given dataset, there must be *n* independent sets of these sinusoids {cos(*kπx_q_*) | *q ∈ n*}, with each set interspersed throughout the collection of available eigenvectors {Ψ_*i*_ | *i* ∈ *N*}.

**FIG. 3:**
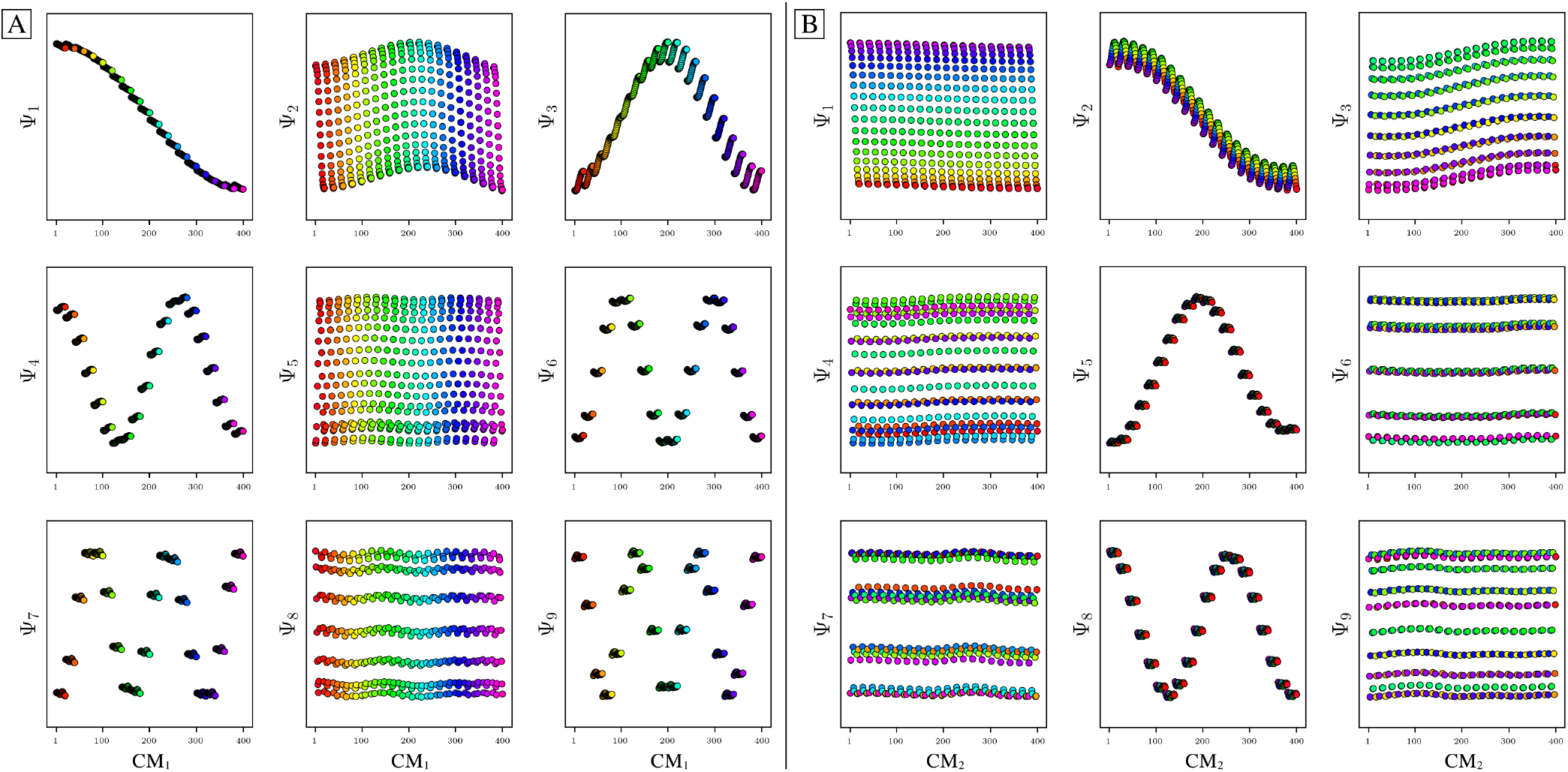
Visualization of eigenfunctions for PD_1_ in SS_2_ (i.e., 20 × 20 = 400 states total making up two degrees of freedom). On the left [A] are the sinusoidal forms {cos(*kπx*) | *k* ∈ ℤ^+^} that emerge for only a specific subset of eigenvectors {*k* = 1, 3, 4, 6, 7, 9*,...*} when points in each Ψ_*k*_ are ordered precisely in the sequence of CM_1_ (as assigned when the ground-truth images were initially constructed). Likewise, in [B], when points in each Ψ_*k*_ are instead ordered precisely in the sequence of CM_2_, a new set of sinusoids emerge {*k* = 2, 5, 8*,...*} precisely for those remaining Ψ_*k*_ not in the previous CM_1_ subset. Hence, it can be seen in [A] and [B] that by systematically ordering the points in each eigenvector in sequence along each degree of freedom present, the corresponding set of sinusoids emerge in the frame of reference of that degree of freedom. However, as stated for SS_1_, such frames of reference are unavailable *a priori*.

Following our previous discovery of a single set of orthogonal Chebyshev polynomials spanning specific 2D subspaces of SS_1_, we next investigated whether similar patterns existed in SS_2_. In doing so, we found that for every conformational motion present in a given state space, there exists a corresponding set of Lissajous curves interspersed across specific {Ψ_*i*_, Ψ_*j*_} projections of the *N*-dimensional embedding. Specifically, in the case of PD_1_, independently projecting the data for SS_2_ onto the planes spanned by its {Ψ_1_, Ψ_*i*_} and {Ψ_2_, Ψ_*j*_} combinations (where *i >* 1; *j >* 2) revealed a unique set of Chebyshev polynomials, with the sequence of points along these trajectories corresponding to CM_1_ and CM_2_ (Fig. 4). With this knowledge in hand, we can now compare the subset of eigenfunctions as obtained in either the reference frame of CM_1_ (Fig. 3-A) or CM_2_ (Fig. 3-B) with the Chebyshev polynomials in Fig. 4. Indeed, each Chebyshev polynomial mapping CM_1_ information in Fig. 4 (visualized with subplots enclosed by blue boxes) corresponds to the subset of sinusoidal eigenfunctions which emerged in the reference frame of CM_1_ in Fig. 3-A; with similar relations holding for CM_2_ in Fig. 3-B. (For convenience, we will refer to a set of Chebyshev polynomials corresponding to a given CM as the *conformational modes*). Thus, even though the knowledge required to view these CM sinusoids is unavailable outside of ground-truth studies, our analysis confirms that these CM relationships are ever-present, and further, that we can rely on their existence—via the composites of carefully chosen eigenvectors—to elucidate conformational type and order.

**FIG. 4:**
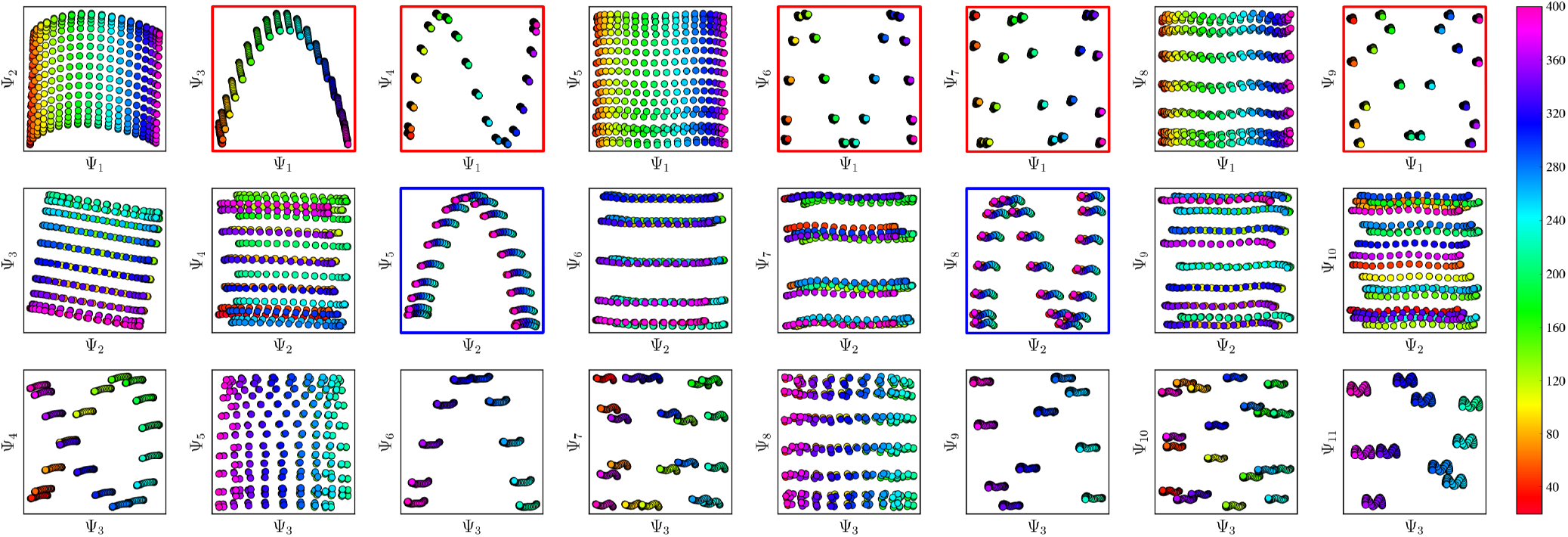
A subset of the space the 2D subspaces for PD_1_ in SS_2_. As demarcated in red and blue boxes, a set of conformational modes exists for both CM_1_ (red boxes, {Ψ_1_, Ψ_*i*_}) and CM_2_ (blue boxes, {Ψ_2_, Ψ_*j*_}; where *i >* 1 and *j >* 2), interspersed throughout each row. The indices for points in each set of polynomials can be visualized here via the corresponding color mapping, where CM_1_ points follow along the full spectrum of colors (i.e., a rainbow with indices 1-400) while CM_2_ points are approximately uniform in color map value (i.e., magenta with indices a multiple of 1-20, with all other colors similarly underlaid). Additionally, note the occurrence of the first parabolic harmonic for CM_1_ located at {Ψ_3_, Ψ_6_}. See Fig. S5 for similar plots obtained for the remaining four PDs.

Combining this empirically-obtained knowledge with our *a priori* understanding of the eigenfunctions of the LBO on known domains, we were able to intuit the analytical form of these Ω_PD_ eigenfunctions. A detailed exposition of this discovery is provided in section SM-XIV, showing how the canonical eigenfunctions on a rectangular domain transform as the data type is translated stepwise from atomic models to 3D density maps to 2D projections. In close approximation, the leading Ω_PD_ eigenfunctions are of the form

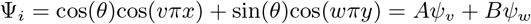

such that a given eigenvector Ψ_*i*_ may contain some linear combination of *n* canonical interval-like eigenfunctions {cos(*kπx*_*q*_) | *k* ∈ ℤ^+^} corresponding to each degree of freedom *x*_*q*_ ⊂ ℝ^*n*^. In Fig. S26, we use this explicit form to near-perfectly emulate the heuristic results obtained in Fig. 3 and Fig. 4. We also show that the these coefficients are conserved across pairs of eigenvectors (i.e., *A*^2^ + *B*^2^ = 1), such that the base functions Ψ′_*i*_ = *Ψ*_*v*_ and Ψ′_*j*_ = *Ψ*_*w*_ can be expressed as a rotation **Ψ** = **R**^*T*^**Ψ′**, having the form

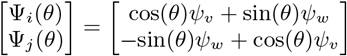

From our analytical expression, it is clear that depending on the PD, CM information – pertaining to each of the system’s degrees of freedom – will lie on some linear combination of the embedded manifold’s orthogonal eigen-vectors. This feature is seen most strikingly in {Ψ_3_, Ψ_4_} of PD_3_ (Fig. S5), where the parabolic surface described by the Chebyshev polynomial is significantly tilted out of alignment with the plane of the 2D subspace containing it. Similar instances, albeit in more subtle form, also arise for surfaces in the remaining three PDs of Fig. S5. In section SM-XIV-C, we demonstrate that this feature is a result of *PD disparity*. Specifically, we generate an embedding (via DM) of the SS_2_ collection of 3D electron density maps (EDMs, from which the PD datasets originate), and demonstrate near-perfect decoupling of all cosine eigenfunctions such that they become independent eigenvectors. Thus, it is clear that the need for eigenfunction realignment is due to the change in interatomic distances dependent on projection direction (Fig. S25). This disparity among PDs is inevitable, and poses a fundamental problem that must be addressed.

As a remedy to this problem, we aim to stitch the CM information of each Ω_PD_ together into one consolidated orthogonal coordinate system. As already shown, since each CM is represented by a set of orthogonal sinusoids (one per degree of freedom), we thus aim to isolate these sinusoids in their complete form within each PD-manifold eigenbasis. As detailed in section SM-XIV, by use of appropriate rotation operators *R*_*i,j*_, the summands within each eigenfunction pair can be maximally separated among a set of eigenvectors (e.g., Ψ_*i*_ = *Ψ*_*v*_ and Ψ_*j*_ = *Ψ*_*w*_) such that an ideal (i.e., canonical) eigenbasis is recovered. As a result of this decoupling of eigenfunctions onto a set of appropriate eigenvectors, each corresponding parabolic surface becomes manually aligned within its 2D subspace, such that the projected structure is again that of the 2D Chebyshev parabola carrying information about a single CM along its curve. In this projected view, states differing in coordinates that are orthogonal to the projection plane (and thus describe ulterior CM information embedded on a higher-dimension surface) overlap – a feature we will take full advantage of later when generating 2D conformational movies. Thus, as long as each parabolic trajectory corresponding to a given CM is aligned with the plane of an independent 2D subspace, we can restrict our study to an analysis of only a few essential subspaces.

To provide rationalization for this technique, Fig. 5-A shows the eigenvectors for the highly-misaligned PD_3_ eigenbasis, ordered along CM_1_. As seen in the first column of Fig. 5-A, while the sinusoids for Ψ_1_ = {cos(*πx*) | CM_1_}, Ψ_4_ = {cos(2*πx*) | CM_2_} and Ψ_5_ = {cos(3*πx*) | CM_1_} are in agreement with expectations, the graphs of Ψ_2_ = {cos(2*πx*) | CM_1_} and Ψ_3_ = {cos(*πx*) CM_2_} appear heavily deformed. As a direct consequence, any Lissajous curve that inherits one of these deformed sinusoids (e.g., any subspace composed in combination with Ψ_2_ or Ψ_3_) will be misaligned with respect to its ideal form (Fig. 5-B).

**FIG. 5:**
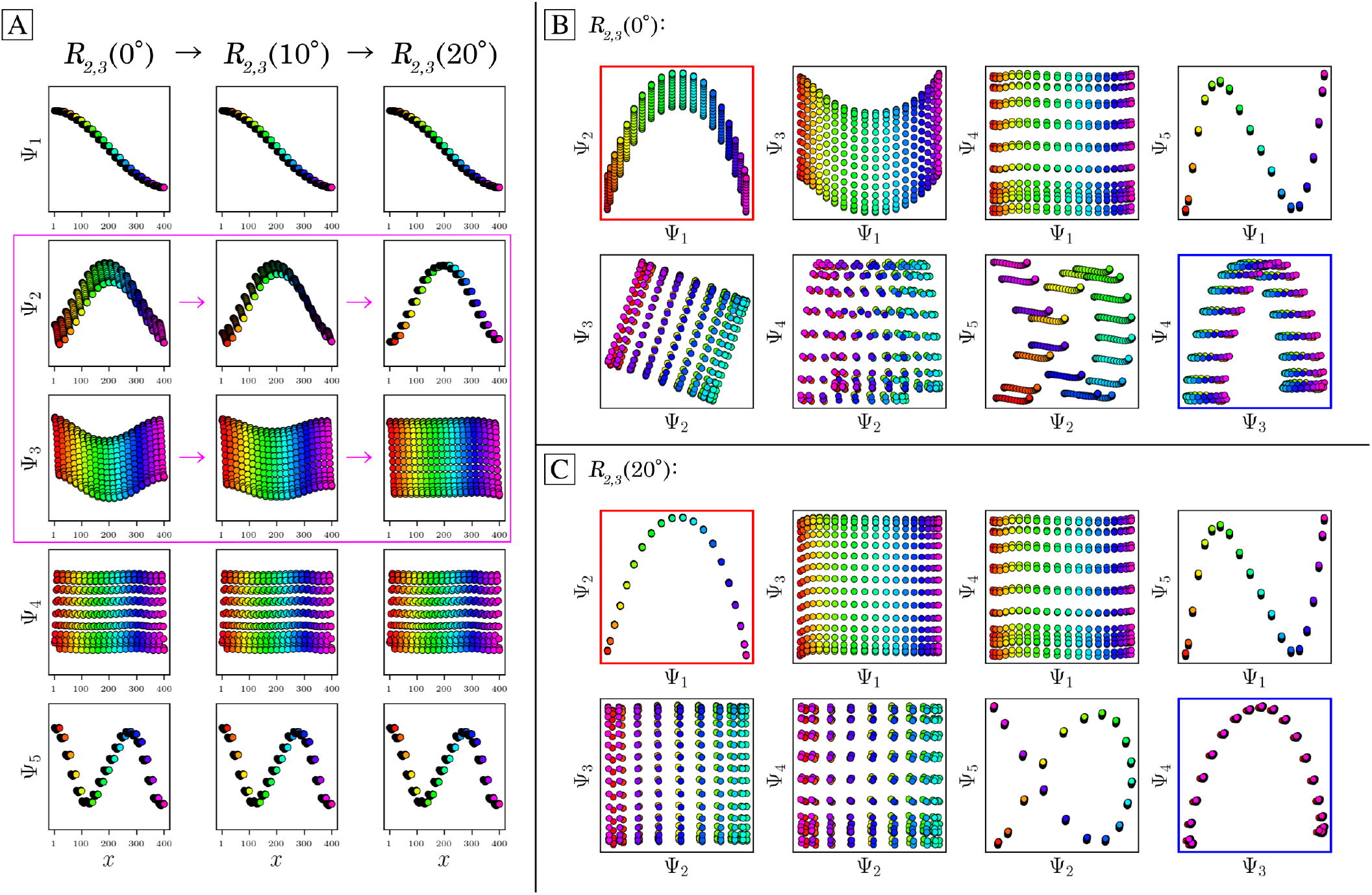
Application of a 5D rotation matrix *R*_2,3_(*θ*) on the embedding generated for PD_3_ from SS_2_. The three columns in [A] display the individual eigenfunctions (as plotted by indices corresponding to the CM1 frame of reference) before the rotation is applied, at *R*_2,3_(10°), and finally at *R*_2,3_(20°), respectively. Note that *R*_2,3_(20°) maximally decomposes Ψ_2_ and Ψ_3_ into unique sinusoids (recalling that the planar distribution in Ψ_3_ is in fact a sinusoid when visualized in the CM_2_ frame of reference, and vice versa for Ψ_2_). The before and after effects of these rotations on the Lissajous curves can likewise be seen in [B] and [C], respectively. Applying *R*_2,3_(20°) properly orients both parabolic surfaces corresponding to CM_1_ and CM_2_ (denoted with red and blue boxes, respectively), such that the eigenvectors are orthogonally aligned with the eigenbasis of the CMs.

Given this insight, we now introduce a method for correcting these misalignments using orthogonal transformations. Specifically, we apply a *d*-dimensional rotation operator of sufficiently large dimensions to single-handedly reorient all aberrant surfaces in their respective 2D sub-spaces. The results of this operation on the embedding associated with PD_3_ can be seen in Fig. 5-B and Fig. 5-C; before and after applying a 5D rotation matrix, respectively. Mathematically, this *d*-dimensional rotation is a subgroup of the orthogonal transformation in *d* dimensions with determinant one. These orthogonal transformations are linear and represented by a *d × d* matrix *O* with the property *O × O*^*T*^ = *I*, where *O*^*T*^ is the transpose of *O* and *I* is the identity matrix. As a consequence, orthogonal transformations leave lengths and angles between vectors unchanged. Each such matrix *O* can further be represented by *d*(*d* − 1)/2 rotation sub-matrices *R*_*i,j*_, with each sub-matrix parameterized by a unique angle and operating on a specific plane. For the specific case of the 5D rotation matrix used in Fig. 5, there exist 10 rotation sub-matrices in total, with each corresponding to a specific planar rotation on the eigenbasis. Of these 10 matrices, we found that only one had to be altered to achieve the results shown, having general form

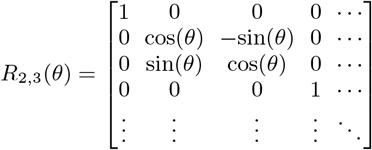

As this *R*_2,3_(*θ*) operator corresponds to transformations performed solely on Ψ_2_ and Ψ_3_ (row 2 and 3, respectively), eigenvectors previously identified as problematic in PD_3_ are thus isolated. The result of this transformation on the full set of eigenvectors can be seen in the three columns of Fig. 5-A, which visualize the *R*_2,3_(*θ*) rotation under 0°, 10° and 20°, respectively (where only Ψ_2_ and Ψ_3_ undergo change, as expected). Intuitively, the outcome of this operation is equivalent to the traditional notion of a vector rotation (for example, consider **e**_**1**_ = (1, 0) ∈ ℝ^2^: just as **e**_**1**_ is given some combination of the secondary dimension *y* with its initial dimension *x* during a rotation (via vector additions and scalar multiplications), so too is Ψ_2_ weighted with Ψ_3_, and vice versa. As seen in Fig. 5-A, along the way in reaching *R*_2,3_(20°), Ψ_2_ and Ψ_3_ have effectively transferred between each other an equal share of their initial content via a series of continuous deformations, with each initial eigenvector thus sharing some combination of the other’s initial sinusoidal form (as is also analytically demonstrated in Fig. S20-C). After this exchange, the initially overlapping sinusoidal information contained in part between Ψ_2_ and Ψ_3_ is maximally separated between both eigenvectors, ultimately resulting in the alignment of all corresponding Lissajous surfaces with their 2D subspaces (Fig. 5-C), as desired. Later in our analysis we will return to this topic under the moniker *eigenfunction realignment*, and describe a strategy for automating these corrective actions for noisy datasets.

#### C. Data-type I in State Space 3

We next investigated the 1000 states making up SS_3_. As before, the eigenvalue spectra were similarly found to be slowly decreasing, but falling off more gradually than in the SS_1_ and SS_2_ spectra. Additionally, for each conformational motion present in a given PD dataset (this time for CM_1_, CM_2_ and CM_3_), a set of unique Lissajous curves were again found spanning specific 2D subspaces of the embedded manifold, with the Chebyshev subset explicitly describing the corresponding CM along a 2D trajectory explicitly. Fig. S7 shows the set of 2D subspaces where these modes exist for PD_5_. To note, due to the increased complexity of the SS_3_ state space, these patterns were much more interspersed throughout the *N*-dimensional embedding, but still followed a similarly predictable ordering. In addition, due to the relatively small range of motion exhibited by the third conformational domain (as seen from these PDs and as designed in the ground-truth structures), all CM_3_ modes were found in higher-order eigenvectors; e.g., Ψ_5_ and higher for these five PDs. As similar patterns were identified in SS_3_ as in previous accounts, for the remainder of our paper, we will hone our focus on mapping datasets generated specifically from SS_2_.

### II. Principal Component Analysis

#### Data-type I in State Space 2

Following our analysis of manifolds using the DM framework, we next performed linear dimensionality reduction on the SS_2_ images in PD_1_ using PCA. Instead of defining the Gaussian kernel as previously used in the Markov transition matrix, we performed PCA on the array of all pixels, with dimension defined by the number of images and pixels in each image (i.e., on a dataset *Z* of dimension *P×N*). Before embedding, we standardized the images in each dataset by removing the mean and scaling to unit variance, and generated eigenfunctions of the resultant *N×N* matrix *Z*^*T*^*Z*. To note an important comparison between PCA and DM, the matrix *Z*^*T*^*Z* is symmetric and positive semi-definite (i.e., all eigenvalues are non-negative)^43^, which is also the case for the Markov transition matrix used in the DM framework.

A set of different projections of this embedding as obtained from selected eigenvectors (i.e., principal components, PC_*i*_) can be seen in the first column of Fig. S17, with results from DM similarly presented for comparison in the fourth column. As demonstrated, the eigenvalue spectra and eigenvectors obtained from performing PCA and DM are almost identical, except for subtle differences in the spacing between states and boundaries for the pristine case (SNR_∞_, i.e., data-type I). These similarities align with our *a priori* knowledge of the existence of quadric surfaces for positive semidefinite matrices, as described in the section SM-XII). The similarity between these manifolds holds for all subspaces explored, and, as will be seen in the next section, the distinction is diminished in the presence of noise. The results of PCA versus DM on SS_1_ and SS_3_ show similar behavior.

### III. Influence of SNR and Statistical Coverage

#### Data-type II in State Space 1 and 2

As SNR is an important attribute of any experimental dataset, we next sought to understand how the structure of these manifolds change with varying SNR and state space coverage. To this end, we first compared the manifolds from PCA and DM for PD_1_ with additive Gaussian noise (generated as described in section SM-IV), such that the images in each dataset had unique SNR ∈ {1, 0.1} consistently applied to all images in a set. The results of this procedure can be seen in the remaining columns of Fig. S17. For both dimensionality reduction techniques, the fidelity of the resulting spectral geometry to the state space ordering decayed with increasing noise level. Within low-SNR regimes, the behavior of the embeddings from PCA and DM were highly consistent, with DM generated within its optimal range of Gaussian bandwidth. Overall, the corresponding spectral geometry obtained from each framework became increasingly similar as the SNR was decreased (Fig. S18).

We next investigated the effects of varying state space coverage across several SNR regimes, and its effects on the robustness of the corresponding manifolds produced by PCA. As the choice of PCA or DM proved irrelevant in these low-SNR regimes, PCA was chosen here so as to bypass uncertainties introduced by the need of additional parameters in DM; i.e., the Gaussian bandwidth. For this study, we used the 20 images in PD_1_ representing SS_1_ (i.e., one full range of conformational motion), and varied both the number of times (*τ*) these *M* = 20 ground-truth states were duplicated as a group— with each instance having a different realization of additive Gaussian noise—and the SNR of each image therein. Here, Gaussian noise of constant variance was applied for each SNR regime and uniquely added to each of the *τM* = *N* images independently, as shown in Fig. S8. An excerpt from the results of our analysis is shown in Fig. 6, where a highly structured pattern emerged. Specifically, when increasing levels of noise was added to each image (decreasing SNR), increasingly larger values of *τ* were required to reestablish coherent structure in the spectral geometry; i.e., the set of Lissajous curves and corresponding Chebyshev polynomials.

**FIG. 6:**
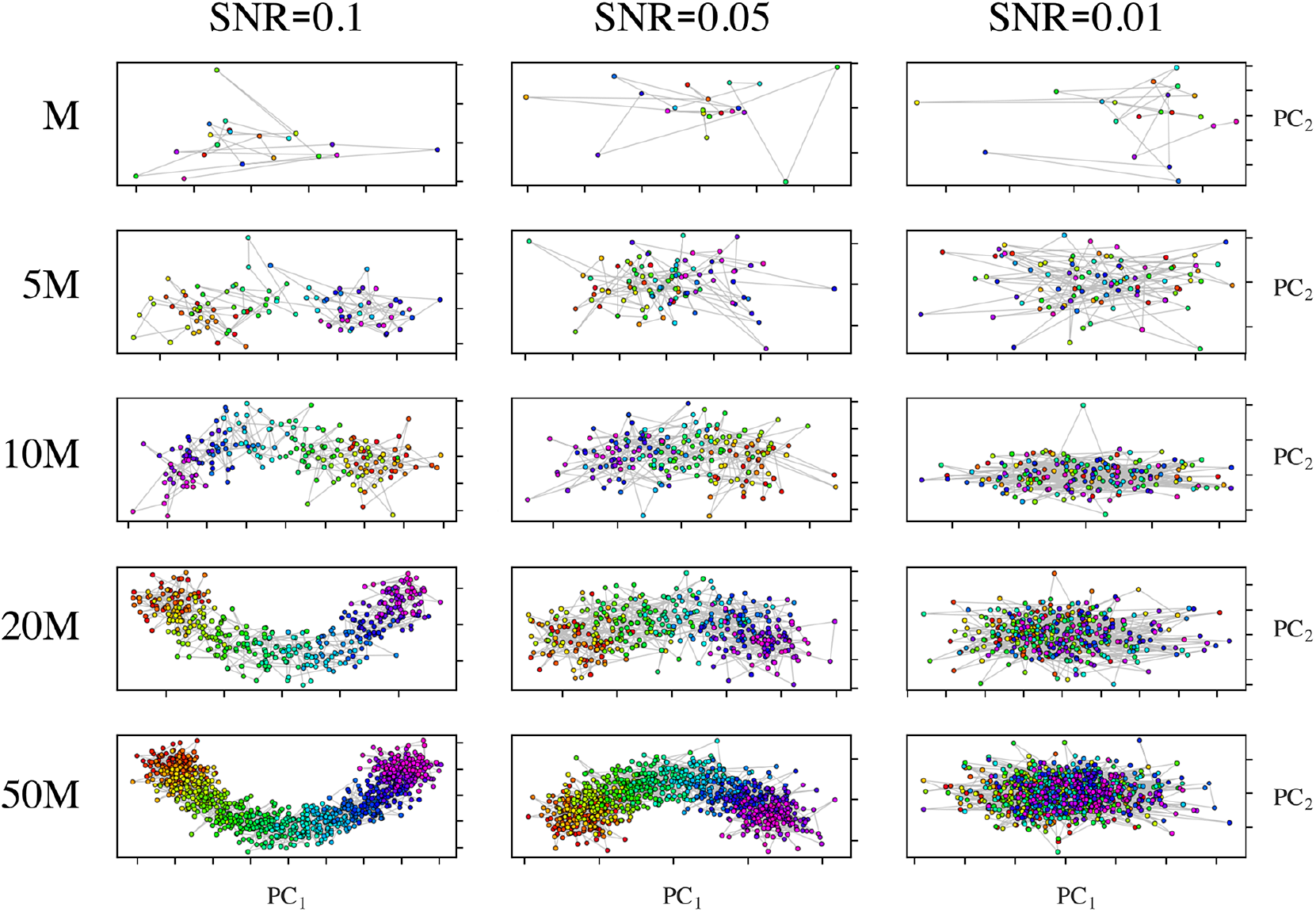
Set of {PC_1_, PC_2_} subspaces produced by PCA from PD_1_ images in SS_1_ over a range of SNR values and state space coverage. As can be seen in the columns, the fidelity of the point-cloud distribution in each subspace to the parabolic form increasingly deteriorated due to decreasing SNR regimes. However, as the *M* = 20 state space was populated by increasing values of *τ* in each of these SNR regimes, the intrinsic parabolic structure of the embedding reemerged. To be precise, 5*M* represents five exact copies of the 20 SS_1_ images (*τM* = 100 images), with unique Gaussian noise added to each image independently as prescribed by its SNR regime. It can be seen here that all values of *τ* shown (up to 50) in the SNR=0.01 regime are too low for recapitulating the intrinsic parabolic structure of the embeddings, and, as further illustrated by the color mapping of their points, no sensible ordering of snapshots can be ascertained within these subspaces.

To quantify these relations, each member of the set of PCA-embedded manifolds in a *τ*-series was fitted with a set of leading Chebyshev polynomials, as seen in Fig. S9 for SNR = 0.1. The coefficient of determination (*R*^2^), which can be interpreted as the proportion of variance in one variable accounted for by another^44^, was then computed for each mode therein. The resultant trends across several SNR regimes are plotted in Fig. S10. Our findings show that as *τ* is increased, the rate at which the geometry of each subspace reaches its most stable regime is dependent on SNR. A critical *τ*_*c*_ value was determined both visually and analytically by assessment of the asymptotes for each SNR regime, beyond which larger values of *τ* provided no further improvement to the spectral geometry.

Further, across all of these regimes, each subsequently higher-order Chebyshev polynomial required a larger value of *τ*_*c*_ to be properly resolved (Fig. S10-A), which is a consequence of higher-frequency patterns requiring more points to resolve when the number of their points (i.e., images) is held constant. For our purposes, recall that the accurate acquisition of only the parabolic trajectory is relevant. As *τ*_*c*_ fluctuates based on numerous unknowns in the experiment, determination of its value for a given experimental dataset is infeasible. Parameters influencing *τ*_*c*_ include not only unknowns such as the number of ground-truth states *M* and SNR regime, but also the intrinsic dimensionality of the dataset and the free energy of the system.

We next describe specific characteristics of CM sub-spaces obtained from datasets generated with these noisy-duplicate images. Specifically, we examine the parabolas generated via PCA from SS_2_ PDs with SNR = 0.1 and *τ* = 10, which will guide several choices made for our framework in the following section. Fig. 7 shows the composite parabolic trajectory and corresponding sinusoidal form of each eigenfunction for CM_1_ and CM_2_ of PD_1_, as well as a collection of similar CM subspaces from randomly-selected PDs. Each subplot has been assigned a color map matching the ground-truth sequence of states of the CM to which it corresponds with this sequence partitioned into 20 equally-occupied bins (i.e., CM states). As can be seen, while each of the two underlying point clouds corresponding to a unique eigenfunction maintains well-defined structure after introduction of noise, CM state partitioning becomes increasingly disordered in their composite parabolic point cloud.

**FIG. 7:**
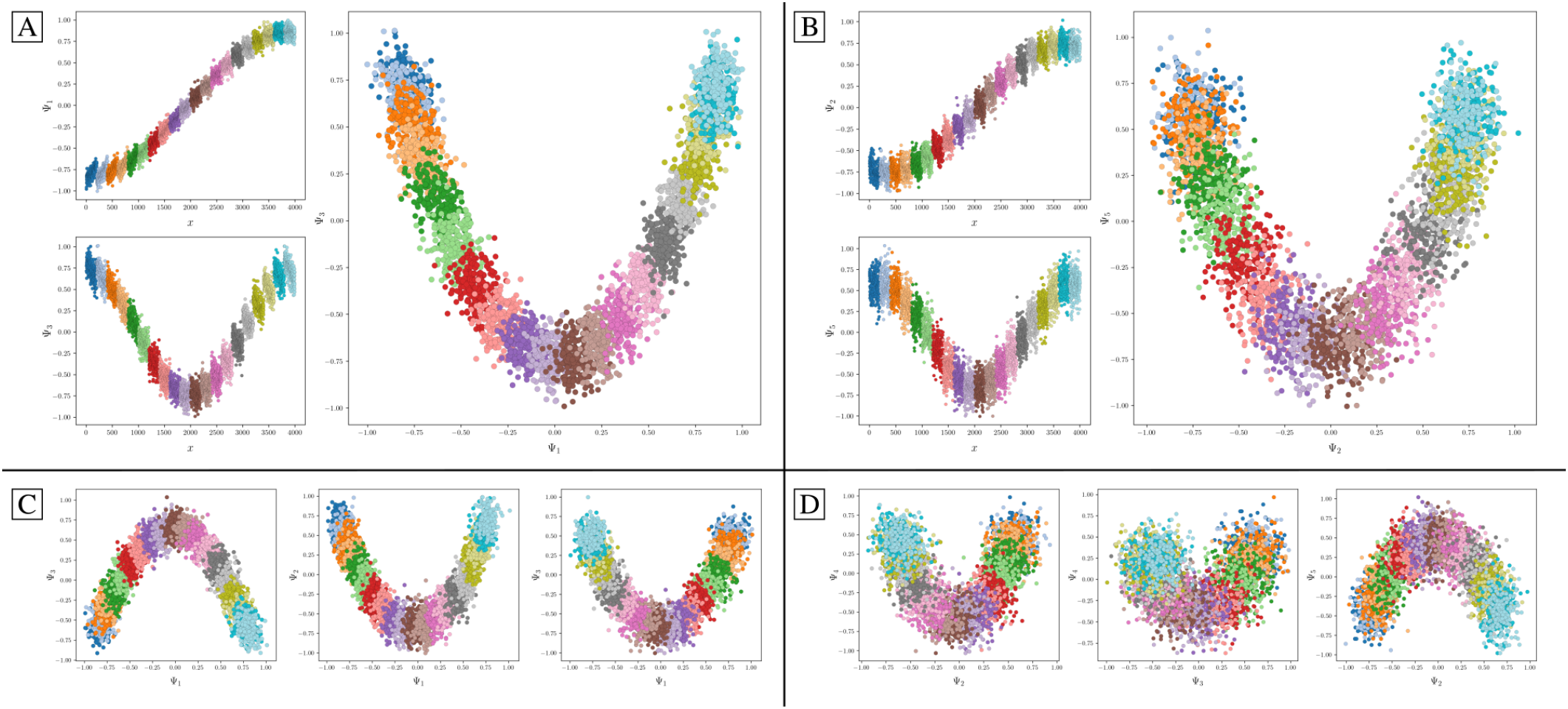
CM subspaces for a set of five PDs generated with SNR = 0.1 and *τ* = 10 and embedded via PCA. The coordinates within each point cloud are colored to indicate their ground-truth CM state assignment, such that each point belongs to one of 20 CM bins, and each bin contains 200 points (with the same coloring scheme used regardless of CM). In [A], the parabolic CM_1_ subspace of PD_1_ is shown along with its two leading cosine eigenfunctions (with each cosine ordered according to its ground-truth sequence). Similarly, in [B], the parabolic CM_2_ subspace of PD_1_ is shown with its own set of leading cosine eigenfunctions. The remaining subplots show a variety of CM_1_ [C] and CM_2_ [D] subspaces for three randomly-oriented PDs, so as to emphasize the variability in features prevalent in manifolds embedded from noisy images.

Additionally, due to PD disparity, the characteristics of each CM-parabola can be seen to vary significantly depending on viewing direction. These variations include average thickness, length, density, trajectory, and spread of data points in each parabolic point cloud, with aberrations occurring most frequently in CM subspaces generated from PDs where the apparent range of the given CM is diminished. As a result, while the CM subspaces for all PD manifolds carry reliable content for recovery of 3D density maps along a conformational trajectory, certain clusters of PDs ⊂ *S*^2^ offer less reliable geometric structure for accurately estimating occupancies of CM states therein. From these initial observations, it is clear that effectively delimiting states in these highly-variable sub-spaces will require robust solutions to be subsequently explored. As a final note for this section, all trends described here for PCA were likewise found to exist for embeddings of manifolds obtained using DM (Fig. S18).

### IV. Influence of TEM Contrast Transfer Function

#### Data-type III in State Space 2

Carrying forward our knowledge gained from evaluation of data-types I and II, we next turn to data-type III for analyzing the PD manifolds obtained from image ensembles generated with experimentally-relevant CTFs and SNR as is encountered in a Transmission Electron Microscope (TEM). For these trials, we first generate and apply a CTF to each image as described in section SM-VI. Specifically, using images from PD_2_ of SS_2_ with *τ* = 10, we assign to each image a random defocus value from the interval [5000, 15000] Å. Such a wide range is usually chosen to compensate for the zero-crossings of CTFs where no information is transferred (Fig. S12), with similar intervals typically used in modern cryo-EM experiments. Likewise for each image, constant values are used for voltage (300 kV), spherical aberration coefficient (2.7 mm), and amplitude contrast ratio (0.1) to emulate typical TEM conditions. These parameters are jointly used to construct a unique CTF for each image, which is applied via multiplication to the image’s Fourier transform. With the collection of images modified by unique CTFs, additive Gaussian noise is next applied such that the SNR of each image in the resultant ensemble is approximately 0.1.

We next set out to measure the extent of interference of the Contrast Transfer Function on the corresponding manifold for an example PD. However, since the collection of images are now sampled using a range of defocus values, they are no longer directly comparable using a standard distance metric. Instead, an adjustment to the kernel must first be made to account for our introduction of CTF. We show here the results of applying the previously established *double-filtering* kernel, which ensures a zero Euclidean distance between any two images that differ only in defocus^1^. In application, during each pairwise Euclidean distance calculation, the Fourier transform of each image is multiplied by the CTF of the image under comparison to compensate for the defocus difference. The corresponding manifold embedding is shown in Fig. 8, which juxtaposes these results with the same dataset generated without CTF and using a standard Gaussian kernel.

**FIG. 8:**
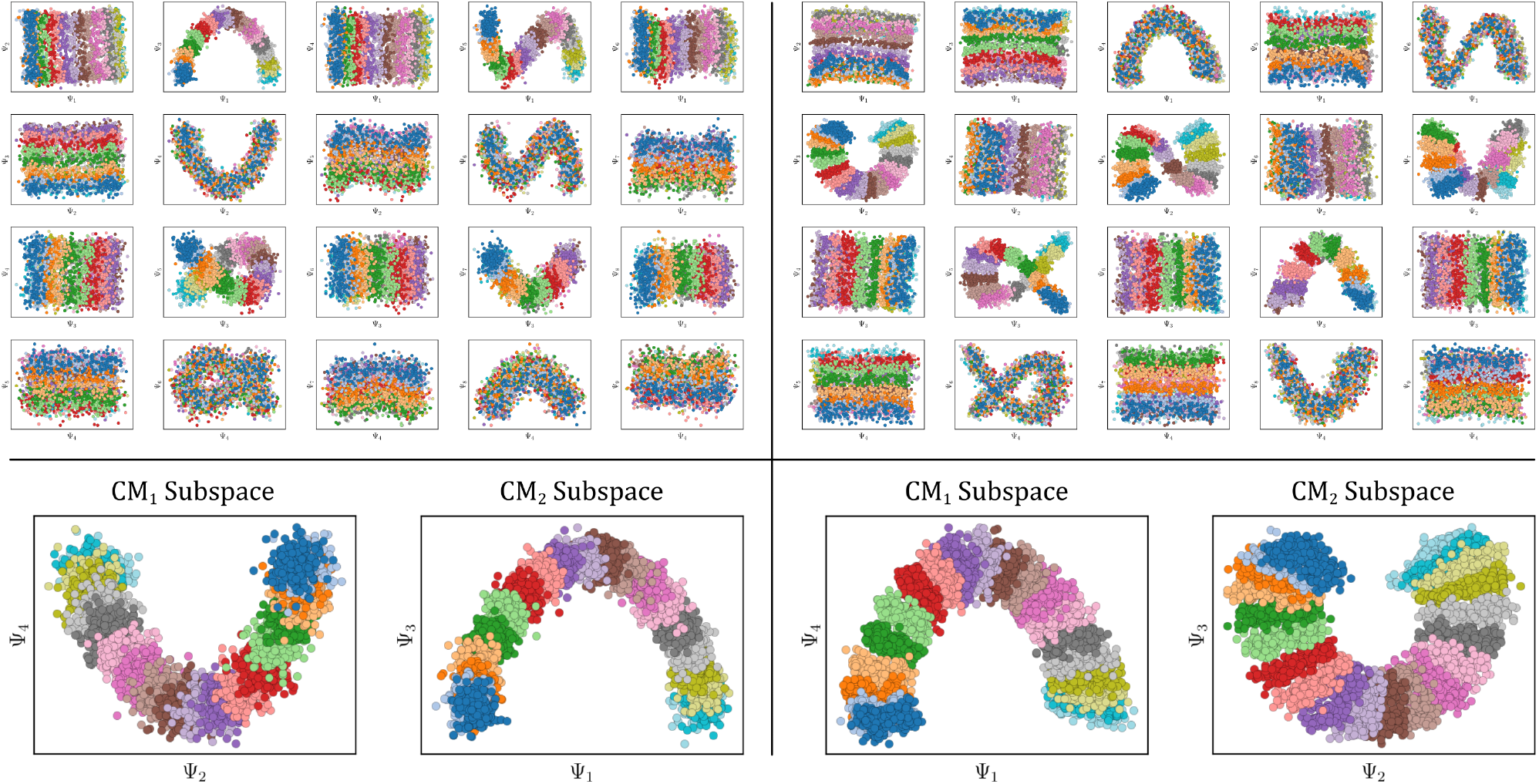
SS_2_ manifold embeddings (PD_2_, *τ* = 10, SNR = 0.1) obtained with and without CTF applied, as shown on the right and left-hand side, respectively. For the case of the embedding obtained from images without defocus, protocols for synthetic generation follow those established in Fig. S12 (A, B). Likewise, on the right, protocols follow synthetic generation of images with microscopy parameters as shown in Fig. S12 (D, E). The non-CTF manifold embedding was generated via DM with the standard Gaussian kernel, while the CTF-manifold embedding was obtained via DM with the double-filtering kernel. On the top insets, colors displayed represent the ground-truth CM_2_ bins, while for the bottom insets, for both sides, colors represent CM_1_ bins (left) and CM_2_ bins (right).

As seen on the right-hand side of Fig. 8, there is a noticeable inward-curling at the ends of the CM sub-space parabolas generated using the double-filtering kernel. Notwithstanding this artifact, we found that the double-filter kernel was successful in preserving the most important aspects of the manifold. This approach also proved superior to alternative techniques explored, such as embedding using a standard kernel from sets of CTF-corrected images. To note, perfect defocus assignments were used here for CTF correction, when in reality these values would be estimated first using established algorithms^45–47^.

### V. Overview of the ESPER Framework

Having explored all three data-types including CTF and noise, we now lay out the ESPER strategy for recovery of conformational motions in the form of 3D movies and a corresponding free-energy landscape. This methodology requires several steps that will be introduced in turn, with the entire process schematized in Fig. 9.

**FIG. 9:**
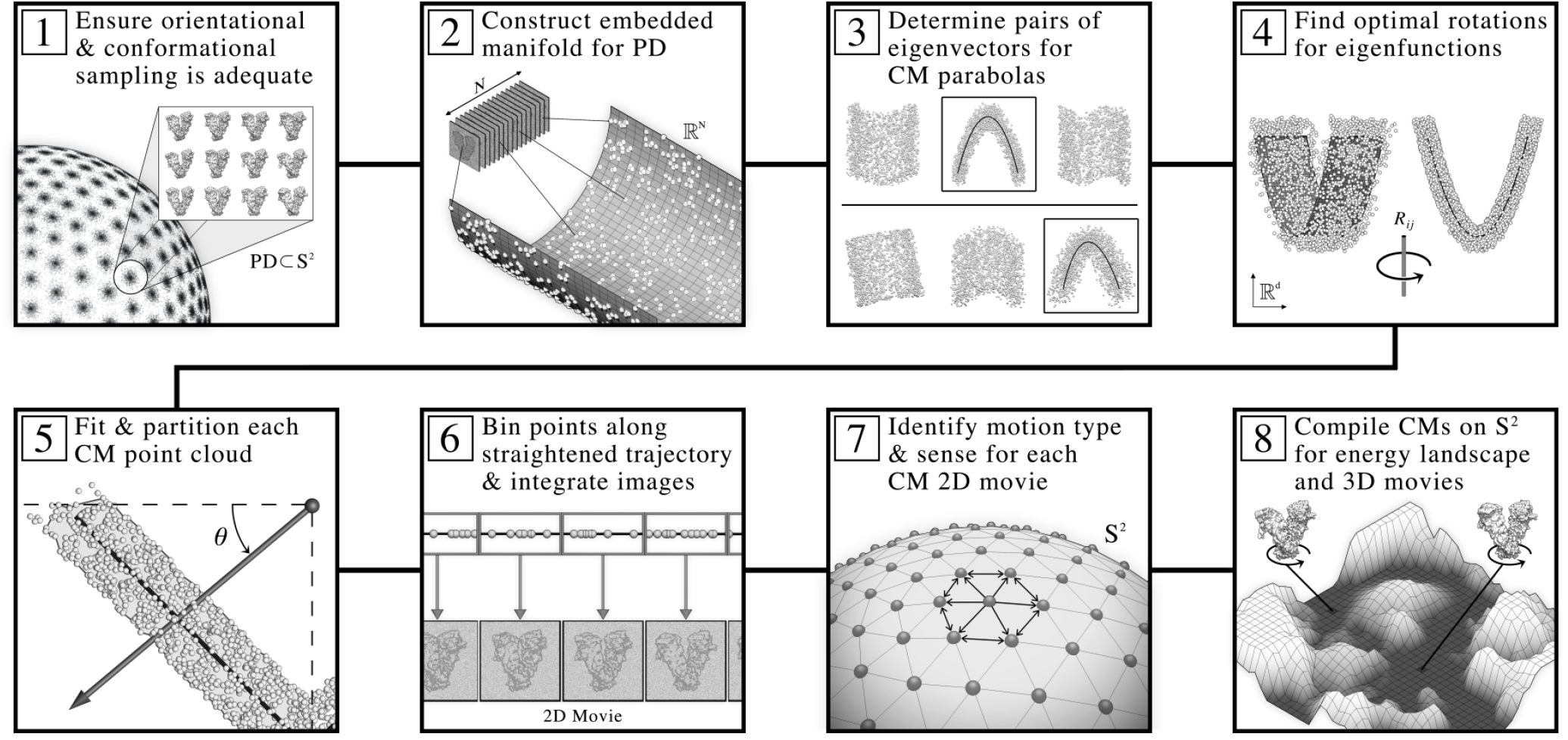
Schematic detailing the ESPER workflow for recovery of conformational continuum as informed by our heuristic analysis. Through this framework, 3D movies and corresponding free-energy landscapes are obtained for the set of conformational motions in a given dataset. Note that the previous ManifoldEM workflow branches off after completion of step 2 above, and after performing a series of alternative steps required by NLSA, then enters again with our pipeline at step 7, before splitting off again to form final outputs (as we achieve independently via ESPER) in step 8. As will be described fully in our Discussion, given certain requirements are met in the quality and structure of a dataset, our method provides an alternative avenue to NLSA for obtaining conformational outputs within the ManifoldEM framework.

The general intuition for our approach is as follows. Ideally, for each Ω_PD_ embedding, we first wish to translate the *n* conformational-variation signals residing along a high-dimensional parabolic surface into a rectilinear *n*-dimensional state space. To this end, one can imagine forming a coarse *n*-dimensional grid along this desired hypersurface—with each *n*-cube (bin) on the grid nonuniformly stretched to occupy an equal volume as required to account for nonuniform rates of change along its complex surface—and accruing the set of points (and thus indices of corresponding images) falling within each bin’s boundary. This procedure should then be repeated for each Ω_PD_ independently. To reconcile the contents of these PD manifolds on *S*^2^, which may contain conformational information along different coordinates due to PD disparity, the orientation of each *n*-dimensional grid (and thus ordering of bins therein) must be aligned so as to match across all PD manifolds. Next, the set of images belonging to each compiled bin can be combined to reconstruct a 3D density map of the molecule, with the total image count used to define a state occupancy. As a result of this construction, an *n*-dimensional occupancy map (and thus free-energy landscape) can be formed, along with a set of corresponding 3D density maps representing every state.

In application, however, there are many complications to this procedure. For one, the desired high-dimension parabolic surface presents difficulties in both discovery and direct mapping. Since there are many such potential subspaces housing parabolic surfaces (even for *n* = 2) within the embedding of a given Ω_PD_, there exists ambiguity as to which one contains the desired information, further exacerbated in the presence of harmonics and experimental artifacts. In addition, due to the complex nature of these hypersurfaces—which can vary in features ranging from elliptical, to parabolic, to hyperbolic, and each with boundary aberrations—it is much easier to instead fit and partition the set of its orthogonal components; i.e., the parabolas residing in easily identifiable 2D subspaces. Given this route, several operations can next be performed on these parabola-housing subspaces to approximate an idealized, straightened trajectory for each CM, ultimately allowing the formation of a rectilinear coordinate system when this set of straightened CM trajectories are recombined. Finally, these rectilinear co-ordinate systems must be organized in such a way that CM content is matched across all PDs, and compiled.

Following this rationale, the ESPER approach first aims to find the set of parabola-housing subspaces (via least-square conic fits) required for elucidating higher-dimensional CM information, while accounting for eigen-vector rotations and carefully eliminating harmonics [steps 3, 4]. Each parabola-housing subspace is next transformed (along with its conic fit) via the inverse-cosine mapping to account for nonlinear rates of change, with points partitioned into contiguous equal-area bins [step 5]. Each bin is then filled with a set of image indices corresponding to all points falling within its geometric bounds. Images belonging to each bin are next integrated to form the frame of a 2D movie [step 6], which is used to identify both the type of CM and its directionality (i.e., *sense*) of its motion [step 7]. As the location of each point (and thus image index) present in a given CM subspace is coupled to its coordinates in all other orthogonal CM subspaces (on the high-dimensional surface), we can reconstruct this joint geometrical relationship using only the intersection of image indices obtained in all pairwise combinations of bins spanning all CMs. By means of this approach, when this information is accumulated across all PD manifolds, the desired occupancy map and index sets required for full recovery of 3D electron density maps in all bins are thus obtained [step 8].

In the following three subsections, we provide a more detailed description of these steps. For the purposes of this exposition, we will use SS_2_ via data-type II for initial demonstrations of eigenfunction realignment and subspace partitioning (subsection A and B, respectively), followed by use of our final analysis data-type (i.e., SS_2_ via data-type III with free-energy landscape) in subsection C to furnish final outputs and assess their validity.

### A. Eigenfunction Realignment for Data-type II

We describe here our methodology for calculating the rotations required for eigenfunction realignment of each embedded Ω_PD_ in the presence of noise with experimentally-relevant SNR. We consider this calculation to be the first step in the ESPER framework (i.e., [step 3] in Fig. 9). To note, after generation of each embedding from PD-images as previously described [steps 1, 2], our methodology deviates here from the existing ManifoldEM method^1,10,11^, which would next move on to NLSA without accounting for eigenfunction realignment on Ω_PD_ subspaces. First, recall that depending on the PD, the observed conformational eigenfunctions may be misaligned with respect to the ideal eigenfunctions of the LBO, requiring application of a *d*-dimensional rotation matrix to align the subspace (Fig. 5). As an example, the effects of applying a 4D rotation to the 4D sub-space for data-type II corresponding to PD_2_ (containing all parabolic modes of CM_1_ and CM_2_ in SS_2_) can be seen in movie M5, where only one of the six required rotation matrices is altered by 28.65° (with the remaining five unaltered; i.e., 0°) to single-handedly realign both parabolic modes (one per CM) to the plane of their respective 2D subspaces.

Based on this behavior, we have developed a technique to automate the discovery of the rotations required to realign essential eigenfunctions in each Ω_PD_ embedding. This algorithm is informed, first and foremost, by our heuristic findings of the existence of parabolic surfaces in each embedding, which correspond to a specific CM. In the case of noisy data, as each corresponding 2D subspace is rotated, it exhibits a unique profile that can be characterized by a sequence of 2D histograms on that subspace, with one 2D histogram per each rotation angle corresponding to a given R_*ij*_ rotation operator. When we plot the number of nonzero bins in the corresponding 2D histogram as a function of rotation angle, the minimum in this distribution corresponds to the angle required to properly counter-rotate each 2D subspace by the current operator (movie M6). After careful observation of all PDs across numerous datasets, we have determined that the exact rotational operators R_*ij*_ required to adequately rotate each 2D subspace are linked to the indices of those eigenvectors housing each CM parabola. As a consequence, we need to first determine the 2D subspaces housing parabolas, and these are identified via the best least-squares fits in each eigenvector row (movie M7). A detailed description for the procurement of this information is available in the section SM-XVI, with an example visualization of this workflow provided in movie M7.

Once these CM subspaces have been isolated, a final 2D in-plane rotation still needs to be applied to orient the parabola into its canonical form. We thus perform a least-squares fit Ψ_fit_ using the implicit equation of a general conic defined by an irreducible polynomial of degree two

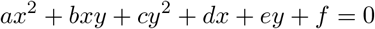

This general conic form can account for parabolas, ellipses or hyperbolas (discriminant *b*^2^ − 4*ac* equal to zero; less than zero; or greater than zero, respectively). As will be seen, this flexibility is essential for fitting parabolic-like point clouds with nonzero discriminant, which are encountered for point clouds with boundary aberrations, and especially those obtained from images modified by the CTF. In this form, the *xy*-term rotates the graph, providing for the possibility of encountering subspace trajectories with an axis of symmetry unaligned from the 2D eigenbasis. This equation can thus be rewritten with a new set of coefficients^48^ to effectively rotate the coordinate axes such that they come to alignment with the axis of symmetry.

### B. Subspace Partitioning for Data-type II

Once the required Ω_PD_ eigenfunctions are correctly rotated into a common eigenbasis as defined by the desired CMs, and each 2D subspace housing CM information is identified for each PD [steps 3, 4], we next partition these 2D subspaces into contiguous equal-area bins [step 5] representing a quasi-continuum of conformational states. Here ESPER differs decisively from the preexisting ManifoldEM workflow e ncompassing N LSA. To h elp distinguish between these strategies, a brief summary of the NLSA workflow h as been p rovided in section SM-XVIII.

The motivation for the ESPER subspace-partitioning approach stems from our analysis of PD disparity in the presence of noise (as shown in Fig. 7), where it is observed that the ground-truth bins and overall area of each point cloud manifest in a variety of sizes depending on PD viewing angle. These observations inspired an area-based point-cloud fitting approach able to correctly chart spatial discrepancies while remaining unencumbered by changing densities (i.e., occupancies) along each trajectory. Fig. 10 provides an overview of our novel strategy for splitting up each CM subspace into a sequence of equal-area bins, with subplots detailing recovery of CM_1_ states and corresponding occupancies for a single PD.

**FIG. 10:**
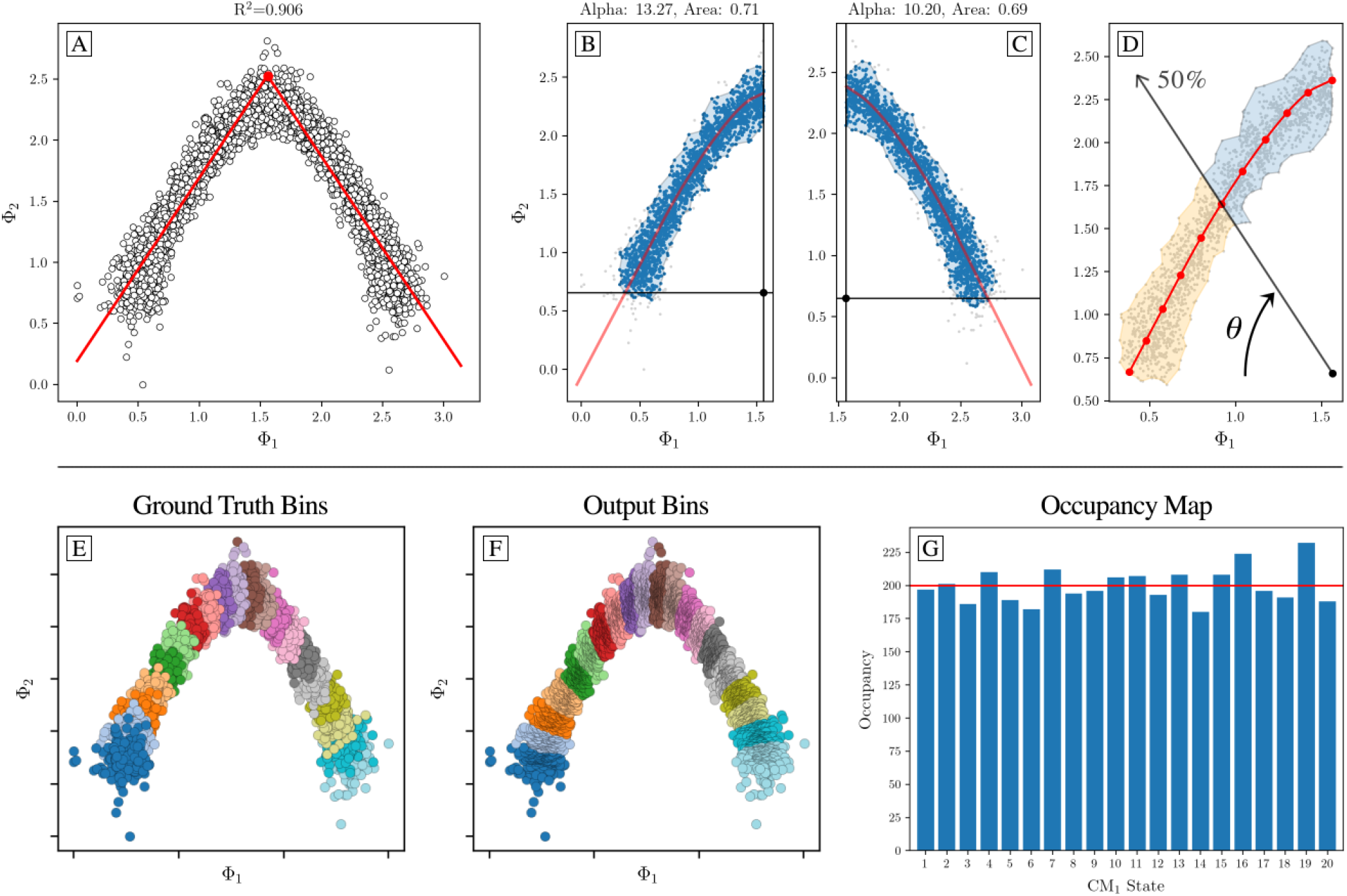
Overview of area-based method for extracting sequential conformational information from a given CM subspace. Subplots [A] through [G] display our algorithm’s outputs on the CM_1_ subspace of an arbitrary PD from data-type II. First, [A] shows the inverse-cosine transformation and corresponding preliminary fit using an absolute value function. Subplots [B] and [C] demonstrate the alpha-shape polygon and Φ_fit_ trajectory defined on each halved subspace, with the anchor-point designated within the central alcove. In [D], a ray is shown passing from the anchor-point through the point cloud. At the current angle *θ* shown, half of the area of the alpha-shape has been traversed, demarcating the boundary between the 5^th^ and 6^th^ (of 10) CM_1_ bins. Subplots [E] and [F] compare the ground-truth bins—as visualized via the known sequence of images in each state—with the final output bins defined via this framework. Finally, the 1D occupancy map is provided in [G], where the horizontal red line (200 images) represents the ground-truth occupancy assignment per CM_1_ state.

To initiate this procedure, for each CM subspace of a given Ω_PD_, we first scale both eigenvectors {Ψ_*i*_, Ψ_*j*_} between [−1, 1] and apply an inverse-cosine transformation on each: *f* : {Ψ_*i*_, Ψ_*j*_} ↦ {Φ_*i*_, Φ_*j*_}. As previously shown in Fig. S4, it is expected such a mapping will induce a space with uniform rates of change between states. To mitigate the overall complexity of operations, the axis of symmetry of the conic equation (with *b′* = 0) is then used to split the subspace into two halves, such that each half can be operated on individually. We next apply a ball tree algorithm^49,50^ to temporarily prune outlier points for heightened accuracy during subsequent steps. Specifically, the ball tree approach clusters points in a series of nesting 1-spheres based on the Euclidean metric, from which we select only those clusters having a minimum number of members.

Following this preparation, we define t he overall area of each halved subspace with a polygon enclosing a majority of our remaining points (Fig. 10-B and Fig. 10-C). This construction is achieved via the alpha shapes algorithm^51–53^: a generalization of the convex hull that defines the boundaries of the point cloud by a series of *α*-discs (1-spheres of radius 1*/α*), such that an edge of the alpha-shape (polygon) is drawn between two members of the point cloud whenever there exists an *α*-disc containing no members of the point cloud and carrying the property that the two points lie on its boundary. A family of alpha-shapes can thus be defined for each halved subspace via the *α*-parameter, ranging from coarse (a convex hull) to increasingly finer fits around the point cloud. Within this class of polygons, there exists a member providing an optimal level of refinement, in which the alpha-shape area and point-cloud area are equal^52^.

For our purposes, the determination of a suitable value for this parameter was automated by generating a sequence of alpha-shapes of increasingly finer complexity up until the resulting alpha-shape – previously defining one polygon – collapsed into two polygons. Through this construction, our point cloud is enclosed by a fine polygon representing the key features of its geometric shape.

Next, the general conic fit Ψ_fit_ is transformed by inverse cosine to form the trajectory Φ_fit_ and split along its axis of symmetry for use on each half of the {Φ_*i*_, Φ_*j*_} subspace. The intersection of Φ_fit_ with the outer boundary of the alpha-shape is used in combination with the position of the initial vertex to form a new anchor-point nested within the central alcove of the point cloud (Fig. 10-B and Fig. 10-C). For each image-point in the point cloud, a ray is next drawn connecting the anchor-point with the image-point, with the intersection of that ray with Φ_fit_ recorded. As a result of this construction, all image-points are uniquely projected onto the Φ_fit_ trajectory. With each image-point now assigned an index along Φ_fit_, a method is next employed to partition the trajectory into segments representing CM states, such that each image-point is ultimately assigned to a single state. Here, we define a ray emanating from the anchor-point, initiated with *θ* = 0° (Fig. 10-D). For each angle *θ* ∈ [0°, 90°], the area of the lower sub-polygon formed by the intersection of the alpha-shape with the ray is determined^54^ to record the overall ratio of that sub-polygon to the whole. We form 10 bins in this fashion along the Φ_fit_ trajectory (per halved subspace), making up 20 bins in total. Finally, we tally the number of image-points assigned to each of these 20 bins (as visualized in Fig. 10-F) to form the 1D occupancy map for the current CM. Importantly, we store the image indices belonging to each bin along the given CM for subsequent use in forming an *n >* 1 occupancy map (to be detailed in the following section C).

A comparison of our outputs with ground-truth is provided in Fig. 10-E and Fig. 10-G, showing an overall agreement with expectations. To note, we have programmed these steps to require no intermediate supervision by using a robust automation strategy, with details for specific subtasks available in the comments of our corresponding code^37^. As a result of this procedure, the points in the embedding corresponding to each conformational motion are independently lined up within a corresponding 2D subspace, such that averaging points together in that subspace only reveals the conformational-variation signal corresponding to the current CM. Hence, images in each bin can next be averaged to generate each frame of the respective CM’s 2D movie. This process is then repeated for the 2D subspace where the second parabolic mode resides (CM_2_), and so on for higher degrees of freedom. The results of this procedure can be found in supplementary movie M2, where we showcase 2D movies for both CM_1_ and CM_2_ as obtained from a subset of these 126 PDs.

### C. Final Analysis – Recovery of 3D Conformational Motions

We demonstrate here the efficacy of our entire framework with a comprehensive dataset of PDs occupying states in SS_2_, using ground-truth images modified with experimentally-relevant CTF and noise. We will additionally note slight alterations to the previously-described ESPER methodology [steps 3-6] that are required for handling our final analysis dataset, which now includes introduction of CTF as well as noise. Finally, once 2D movies have been obtained from each Ω_PD_, we describe here the conclusion of the ESPER framework via an efficient method for compiling CM information from all PDs [step 7] to create a free-energy landscape and corresponding set of 3D movies [step 8].

First, we note that the minimum number of equispaced PDs (PD_min_) on a great circle required for 3D tomographic reconstruction at a given resolution is defined by the Crowther criterion^55^ PD_min_ = *πD/r*. Here *D* is the particle diameter (120 Å, as measured in state 20_20 of SS_2_) and *r* is the targeted resolution of the reconstructed volume (for our purposes, 3 Å as chosen to match the resolution of our ground-truth maps). According to this criterion, we generated 126 equidistant PDs spaced approximately 1.5° apart along one half of a great circle (labeled as Great Circle 1 in figures that follow), chosen so as to avoid redundant information due to diametric mirroring. Each of the 400 SS_2_ states in each of these PDs was then duplicated based on assignments imposed from a fictitious occupancy map (see section SM-VII) resulting in 4000 images per PD, with each particle modified by an individual CTF having randomly-assigned defocus [5000, 15000] Å and the same microscopy parameters as previously described. Finally, additive Gaussian noise (SNR = 0.1) was applied to each image such that 504,000 unique images were created in total. Euclidean distances among the 4000 images within each PD were then calculated using the aforementioned defocus-tolerant kernel with matching CTF assignments. Finally, following the DM framework, a Markov transition matrix was produced for each distance matrix and diagonalized for subsequent Ω_PD_ analysis.

As was discovered for data-type I and II, we found that a substantial number of these 126 PD-manifold embeddings had misaligned eigenfunctions from our preferred, common coordinate system, with the magnitude of counter-rotations required varying significantly from one PD to the other. For all 126 PDs, our previously-described rotation-automation strategy (see ESPER sub-section A) correctly isolated CM_1_ and CM_2_ and counter-rotated each CM trajectory into the plane of its 2D sub-space. As one small adjustment, we substituted the use of the general conic fit for constrained parabolic fit at the beginning of our eigenfunction-rotation algorithm, which proved essential for evaluating initial subspaces housing inward curling parabolas due to the presence of CTF (Fig. S14). A histogram of the magnitudes of rotations used across all rotation operators—which varied nontrivially among the PDs—is provided in Fig. S28.

With the required CM eigenfunctions in these 126 PD-manifold embeddings correctly rotated into a common eigenbasis and each 2D CM subspace identified, we next proceed to partition each 2D subspace into bins representing different conformational states. This procedure follows our previous description (see ESPER section B), with only slight modifications required for the case of images modified with CTF. First, to combat the effects of inwards curling (Fig. 8) that now encumber ray projections, the location of the anchor-point was moved closer towards the central alcove of the Φ_fit_ trajectory, such that it comes to reside halfway between the center of the {Φ_*i*_, Φ_*j*_} subspace and its previous position on the *y*-axis. Additionally, the emergence of a 20-PD “blind spot” on our 126-PD great circle—where images of CM_2_ were highly obfuscated while CM_1_ remained pronounced— inspired the creation of an alternative branch in our partitioning procedure. In the presence of CTF, we found that while CM_1_ subspaces remained highly parabolic for these 20 PDs, the parabolas corresponding to obfuscated CM_2_ signals were much less appreciable in structure. These instances could be easily predicted by simple evaluation of each subspace’s coefficient of determination (*R*^2^), with a low score indicating that the conic fits and corresponding anchor-points had proven suboptimal. Using an *R*^2^ cutoff (0.6) as a criterion, we devised an alternative strategy for dealing with these aberrant subspaces, where, instead of a general conic fit, the subspaces were fit using an absolute-value function (as is shown, for demonstration only, on a more well-behaved subspace in Fig. 10-A), and anchor point re-assigned such that nearly vertical projections were taken across the point cloud. In effect, of the two eigenvectors making up the subspace coordinates, the influence of the leading eigenvector was made more prominent, such that the contiguous bins were delimited with near-vertically aligned borders. This added flexibility made our partitioning algorithm robust in the presence of less-structured distributions, with results validated via examination of ground-truth bins.

Finally, we note that when CTF-modifications are present, we first individually CTF-correct a nd Wiener-filter^5^ each image before integration within each CM bin. A subset of the final 2D movies produced through this framework are available to view in supplementary movie M3. We have also provided a comparison of these final ESPER outputs to NSLA for one degree of freedom in the section SM-XIX. Through analysis shown in several movies, we found that ESPER generates 2D movies of noticeably higher quality than NLSA, and with significantly more accuracy in occupancy assignments for corresponding states. We further highlight problems that can emerge in the alternative NLSA pipeline, such as the inability to capture certain conformational motions captured properly by ESPER, and the possibility of delivering nonsensical (i.e., physically impossible) results. In addition, we found that the overall computation time of NLSA far exceeds the one needed by ESPER.

After generating all 2D movies (one per CM for each PD), both the type of conformational motion present in the 2D movie (e.g., CM_1_ or CM_2_) as well as its *sense* must be determined individually for each PD. As to the definition of the *sense* of each movie, it cannot be said *a priori* in what direction (i.e., the sequential ordering of states) a CM trajectory is following along any path^56^. For example, for CM_1_, the parabolic mode could either be charting the trajectory from states 1 to 20 or from 20 to 1. This uncertainty is due to arbitrary eigenfunction polarity which naturally arises via eigendecomposition^57^. Although a comprehensive method has been developed to solve this problem for datasets with large numbers of sufficiently occupied PDs using optical-flow and belief-propagation algorithms^56^, considering we only had 126 PDs to decipher, we opted to instead determine the type of CM and its sense with perfect accuracy by visual inspection of the 2D movies. With CM types and senses assigned, the 2D movie of a given CM—housing indices of all images within its frames—can next be compiled together with all other 2D movies (and corresponding 1D occupancies) of that same CM across all PDs. If we desired only one degree of freedom as output, our task would next be complete after reconstructing 3D density maps from the images accumulated in each frame of the given 2D movie (and similarly, as it applies, for the 1D NLSA approach).

For presentation of the intermediate 1D occupancy results, we performed this compilation on both CM_1_ and CM_2_ independently, with corresponding occupancy statistics accumulated for each state therein, and compared with ground-truth knowledge (as shown on the left in Fig. 11). For our comprehensive dataset, we found that the 1D occupancy map distributions were in strong agreement with ground-truth knowledge, with states on average monotonically captured along each subspace trajectory and having a relatively small spread of uncertainty for each bin. Still, noticeable disagreement emerged for both CMs near the boundaries of these distributions, where inward curling of the parabolic point cloud due to CTF is most prominent. As a result, 1D occupancy assignments are slightly skewed (overestimated) near the boundaries in comparison to ground-truth expectations (Fig. S13).

**FIG. 11:**
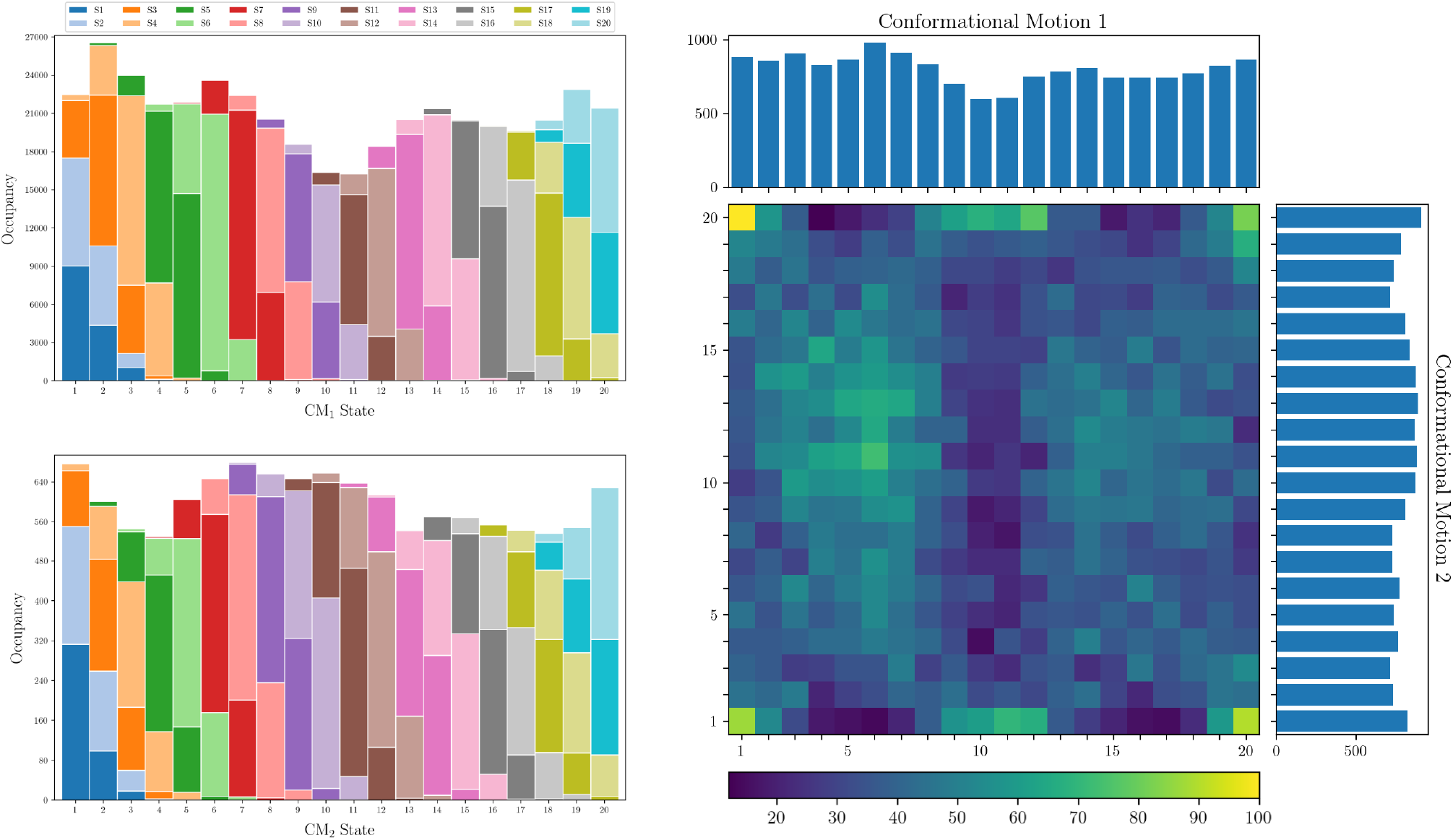
On the left, the final occupancy maps for the 20 states in CM_1_ (top) are shown alongside an equivalent representation for the 20 states in CM_2_ (bottom). Each plot was obtained by integration of the corresponding 20 bins (corrected for sense) in each of the 126 PDs. The total number of images as assigned to each state via our subspace fitting procedure is shown by the height of the 20 bars. Within each bar, the different colors represent how many of the assignments therein belonged to which ground-truth states (as seen in the legend), allowing an assessment of the True Positive rate. On the right, the final 2D occupancy map for the 400 states formed by CM_1_ and CM_2_ is shown; obtained via the intersection of image indices in all pairwise combinations of CM_1_ and CM_2_ bins (corrected for sense) in each of the 126 PDs. Refer to Fig. S13 for a direct comparison with ground truth. Finally, to circumvent issues stemming from inclusion of CM subspaces with poor geometric structure, we note that while all images are used for subsequent 3D reconstructions, only those occupancy assignments for CM subspaces above an *R*^2^-threshold value (0.7) were integrated during this analysis. We additionally note that all results shown are the product of our robust automation strategy involving no case-by-case intervention; assuredly, these deviations could be further mitigated by enforcing key parameter choices with supervision.

As an aside, in order to further investigate these trends, we repeated this analysis for data-type II independently along three orthogonal great circle 126 PD-trajectories. The results of this analysis are provided in Fig. S11, where we plot 1D occupancy maps for CM_1_ and CM_2_ along with corresponding histograms detailing the *R*^2^ values of all 126 respective CM subspaces. First we note that occupancy statistics become aberrant near the boundaries to different extents in all occupancy maps observed, a problem seemingly unavoidable based on our observations of corresponding embedded geometries. As this problem exists, to some lesser extent, even without CTF modification of the data, it can likely be attributed to Neumann boundary conditions^24,57^ (i.e., vanishing normal derivatives). It is also clear that the overall geometric quality of the collection of CM subspaces present (described here via *R*^2^) is a defining f actor affecting the fidelity of the corresponding occupancy assignments to ground truth. Artifacts from PD disparity aside, all occupancy maps remained in excellent sequential agreement, with the significance of occupancy surface fluctuations highly dependent on the quality of CM subspaces available. With this understanding, we now return to our final data-type analysis to conclude with the remaining steps of the ESPER framework.

As previously pointed out, these two CM coordinates are intrinsically linked by the independent occurrence of image indices from the same PD image stack. This fact is used by ESPER to generate a 2D occupancy map, and likewise for any number of degrees of freedom present in a dataset. Specifically, this operation was performed by taking the intersection of image indices (overlap) corresponding to each pairwise combination of bins in the CM_1_ and CM_2_ trajectories: effectively reconstructing the hypersurface on which they jointly reside. For more information on this procedure, see section SM-XVII. The simplicity and efficiency of this operation is based on our method’s continued use of the raw cryo-EM images and initial manifold embedding. In contrast, the original ManifoldEM^1,10^ workflow requires a radially dense set of 1D profiles derived via N LSA (180 in total for *n* = 2, each obtained from an independent NLSA analysis) from the *n*-dimensional subspace formed by the previously-selected set of *n* eigenvectors. Naturally, this operation rapidly increases in computational time as *n* is increased. An inverse Radon transform is next applied to these NLSA profiles to reconstruct an *n*-dimensional occupancy map. The result of our ESPER operation is shown on the right in Fig. 11, which encompasses occupancies for all 400 (20×20) bins in a 2D state space. As each of these 400 bins contains the indices of images sharing a given bin coordinate in the {CM_1_, CM_2_} plane, 3D density maps can next be produced for each state in the state space.

Following this assessment, image stacks, one for each state, were generated and paired with an alignment file that carried the input alignment and microscopy information for each image therein, as initially defined for each PD along the 126-PD great circle trajectory. This file was then used as input for the `relion_reconstruct` module^46^ to create a 3D reconstruction for each of the 400 states. These 3D density maps were loaded in sequence to create 3D movies seen from different views using Chimera^58^. As shown in Fig. 11 and qualitatively expressed in movie M4, these 3D density maps uphold the spatial relationships in the ground-truth CMs with striking accuracy. Of most importance, and presenting a key distinction from NLSA results, is the fact that the mobile domains in all states are as well resolved as the immobile domains.

As a quantitative validation, we calculated the Fourier shell correlations^5^ (FSC) curves between several ground-truth 3D density maps and their corresponding ESPER recovered 3D density maps. FSC curves are a routinely used tool for map validation in cryo-EM and suited to provide a global indicator of agreement between two 3D density maps. The FSC curves confirmed a good recovery of the different states up to a resolution near 3 Å, the ground-truth value. Additionally, Q-scores^59^ were used, not to estimate the resolution, but as a local quantitative validation of the structural fidelity of these outputs. Using this approach on the ground-truth atomic-coordinate structures and their corresponding ESPER recovered 3D density maps, we found highly favorable agreement across all residues for each state. Results for an example state are available in section SM-XX.

## DISCUSSION

Through our analysis, we have identified the way sets of images originating from a varying atomic structure are represented in the manifold embeddings obtained by DM or PCA dimensionality-reduction techniques, and how this information can be used to retrieve the original, ground-truth conformational motions. Our findings on synthetic noisy datasets provide a number of new insights, and emphasize the need for a refined workflow when analyzing the eigenvectors from embeddings of single-particle cryo-EM datasets of molecules exhibiting conformational motions. Several of the operations introduced in this study offer straightforward improvements on the founding PD-manifold approach ManifoldEM^1,10,11^, such as our informed subspace fitting procedure using specific combinations of eigenfunctions, exclusion of parabolic harmonics, and a novel direct retrieval of each CM using the raw cryo-EM snapshots as arranged within the initial embedding. In the last case, the use of the raw images improves both the accuracy of occupancies and final resolution of 3D structures, while providing a vastly simplified workflow for generation of multidimensional free-energy landscapes. Further, we found the corrections for previously unaccounted *d*-dimensional rotations to be essential; the absence of awareness of which can lead to serious systematic errors downstream (see section SM-XIX).

All of the findings within this study are based on heuristic information obtained from ideal, controlled datasets, analyzed so as to maximize the fidelity of our final outputs with ground truth, while uncovering key limitations and uncertainties that could potentially emerge within this unsupervised machine-learning framework. It is important to be aware that results from synthetic data will always be superior to experimentally-obtained data, since even the most sophisticated simulations will be unable to capture all complexities existent in an experimental system. These complexities can be considered as introducing higher-order terms in our parameter space, which has been designed to emulate all lower-order terms up to a threshold deemed satisfactory. Any limitations or uncertainties that do emerge using synthetic data should be anticipated to arise in real-world data, and potentially in exacerbated form.

Contextually, this heuristic analysis is focused on data models originating from molecules undergoing collective rigid-body motions, which we believe are sufficient for most molecular machines, but may fall short of addressing instances involving more complex motions. This is especially the case for those machines entailing the concerted binding and release of ligands, which naturally require a separate state space for each possible combination of the machine with each of its binding partners. For such a situation, a study of ligands has been performed with the founding ManifoldEM approach^10^ using two state spaces, which could similarly be explored in an extension of the ESPER framework.

It is also worth noting that while the results of our heuristic analysis are most relevant to machine-learning methods dealing with projection data (i.e., requiring the PD-manifold approach), several portions of our analysis can be extended to other experimental techniques dealing with alternative manifold inputs, such as the use of atomic models in molecular dynamics and 3D density maps in cryo-electron tomography. Specifically, in section SM-XIV, we detail how the structure of manifolds obtained from a conformational state space transforms as the data type is translated stepwise from atomic models to 3D density maps to 2D projections. Broadly, we believe that there is a potential for the application of our methodology to a wide range of experimental datasets, and particularly those obtained from systems exercising multiple, continuous degrees of freedom.

With that said, we return to the flowchart presented in Fig. 9, to discuss each step of the analysis in detail, focusing on both implementation and existing limitations.

### 1. Ensure that orientational and conformational sampling is adequate

Before initiating this workflow, it is first necessary to ensure that adequate coverage has been obtained via cryo-EM imaging of the heterogeneous ensemble of molecules. We have identified two main categories of coverage: (i) *coverage of S*^2^: overcoming the effects of missing orientations with sufficient sampling of viewing directions; and (ii) *coverage of states*: imaging all available conformational states present and in sufficient abundance for the given SNR to obtain robust statistical coverage in the corresponding manifold. For datasets with low image counts in many PDs, the corresponding final 3D reconstructed volumes will ultimately suffer a loss in resolution. This loss is manifested when the CMs in each Ω_PD_ are stitched together, as not enough 2D CM information is present to properly depict its corresponding aspect of the 3D structure. For assessing coverage of *S*^2^, we have confirmed that within our framework the availability of data over a great circle is sufficient for recapitulating ground-truth conformational information (movie M4).

As for state space coverage, we found that the combination of two parameters—image SNR and state-space occupancy (e.g., *τM* = *N* for uniform distributions)— together served as a strong indicator for the ability of each Ω_PD_ embedding to render coherent conformational-variation signals. Specifically, for sets of images in SNR regimes of approximately 0.05 to 0.1, as encountered in low-exposure cryo-EM^60,61^, multiple noisy duplicates (as defined by *τ*) of the fundamental state space (having *M* states) were required to recapitulate ground-truth structure within the spectral geometry (Fig. S9 and Fig. S10). To note, for experimental conditions where each state occurs with a given frequency as dictated by its underlying free energy, this condition must be met for the least abundant states in the dataset. This extended coverage effectively serves to drown out the experimental noise up to the point where only the conformational-variation signal remains. Without this additional coverage, in the worst-case scenarios, the ordering of the ground-truth points within the embedding will be jumbled in an un-interpretable form, with the distribution of these points closely resembling a Gaussian distribution (e.g., as seen in the last column of Fig. 6).

However, even embeddings of manifolds with adequate state space coverage can still appear globular in structure while retaining proper arrangement of states, with this trend decreasing as coverage is increased (as shown in the second and third columns of Fig. 6). We found that as the *τ*-value is increased, there exists a lower *τ*-threshold (*τ*_*c*_) such that the arrangement of points in the embedded manifold is in highest achievable consistency with its ground-truth state space. In other words, there is a fixed amount of coverage that is sufficient. Unfortunately in an experiment, since the number of ground-truth states *M* for a given molecular machine is unknown, so too is this threshold particle count (*τ*_*c*_), which can be affected by the sample characteristics as well as the quality of the collected data. This threshold was additionally found to vary depending on both the intrinsic dimensionality and energetics of the ground-truth data – with knowledge of either of these parameters not immediately clear for a given dataset. In practice, then, the insight gained about threshold particle count is of no help in guiding the experimental design, and hence the decision on adequacy of state space coverage must be made by trial and error.

One would first aim to collect as many particles as possible during the experiment, and, after generation and embedding of manifolds, make a decision on the adequacy of that collection based on the presence of robust parabola-housing subspaces. Such subspaces were directly observed and showcased, for example, in the conformational embedding of the ribosome^1^. As one potential scheme, for each Ω_PD_ individually, after 2D subspaces have been fit and *R*^2^-values computed (Fig. S14), an *R*^2^-threshold can be used across all subspaces to assess geometric conditions and demarcate use of either NLSA or ESPER on that manifold. As seen in movie M8, these two frameworks share considerable overlap in their 2D movie outputs, and may need only their final PD-outputs subsequently combined for production of sensible 3D movies. Additionally to consider, while it has been shown that ESPER is by far the better choice for manifolds with geometrically-structured subspaces, only NLSA can be applied in the regime of manifold embeddings completely lacking discernible form, while still potentially incurring its known limitations and uncertainties^11^.

As a final note for this section, experimental data suffer from a wider range of nuisances than we have accounted for in our simulation, including the occurrence of aberrant particles (such as ice shards or foreign bodies); uncertainty in CTF estimation; and uncertainty in angular alignments, which are more pronounced in heterogeneous data. While numerous preprocessing algorithms exist to handle each of these instances^27,46,47^, it is obvious that such artifacts and errors, if left unaccounted for, can have detrimental consequences for the fidelity of the corresponding manifolds to the rules we uncovered using the simulation.

### 2. Construct embedded manifold for PD

The choice of method for obtaining eigenvectors— through either linear or nonlinear dimensionality reduction frameworks—proved to have relatively minor consequences compared with many other choices in our workflow. As previously noted, both PCA and DM approaches aim to achieve a description of the dataset’s most fundamental form, defined by the multidimensional relationship among all images in a PD. Surprisingly, PCA and DM produced almost identical eigenvectors for all data examined in this work, particularly in the presence of noise (as seen in Fig. S17 and Fig. S18). The high similarity in the performance of these methods did not conform to previous expectations, as the superiority of nonlinear dimensionality reduction frameworks to linear ones is a belief often cited in the field^1,4,24,38,62^. In contrast, however, a comparative review of twelve prominent dimensionality frameworks^13^ has also shown that, despite their ability to learn the structure of complex manifolds, most nonlinear techniques are unable to out-perform PCA on experimentally-obtained datasets. In the presence of noise, our discoveries indicate that biological objects (such as macromolecules) undergoing combinations of rigid body conformational motions fall in the latter camp and can be effectively studied using either linear or nonlinear techniques.

Given such a choice in embedding strategies, it would still make sense to give preference to DM, as it leaves room for the possibility of cases yet to be encountered, such as those where PCA may have difficulty in untangling certain types of complex conformational relationships. The DM framework additionally offers a reduction in computational load of the eigendecomposition due to its sparse matrices, which becomes increasingly relevant as the number of snapshots increases. However, certain caveats must still be considered. For one, discretion is introduced in all nonlinear frameworks through the fitting of additional parameters required for their optimal performance, failure of which can lead to systematic errors^13^. Using the right strategies, these uncertainties can be minimized. In the case of DM, while incorrect choice of the Gaussian bandwidth had drastic consequences, this parameter was consistently put into the correct ballpark using bandwidth estimation plots. It is also necessary to point out that while DM and PCA examined in detail here are two of the most prominent dimensionality reduction techniques, it might be well worth investigating the performance of other approaches^13^ using simulated datasets such as ours.

Finally for this section, we discuss the use of the defocus-tolerant kernel^1^ for calculating Euclidean distances between images on which a CTF was imposed. We found that its success dwarfed that of two other techniques explored. For comparison, we first observed that manifold embeddings obtained using the standard kernel without any CTF correction were completely incoherent in structure, as expected. Better behaved but still flawed were embeddings obtained from sets of CTF-corrected images (Fig. S12-G) using the standard kernel, which ultimately provided structure suboptimal to those obtained using the double-filtering kernel. We consider this study the first demonstration that the previous double-filter kernel introduced^1^ is more effective than CTF-correction of individual images. However, as previously noted, this kernel is not free of artifacts.

Most significantly, CM subspaces typically had a strong proclivity for curling inwards near the outer edge, with states clumped more densely in these regions (Fig. 8). This trend was apparent enough to require the introduction of general conic fitting strategies for elucidating CM subspaces (Fig. S14). Future studies could further examine the effects of this defocus-tolerant kernel on a wide range of defocus-value intervals and relative magnitudes. As a note, when altering the defocus range, one must carefully choose the particle box size. If the box size is too small to fit the broadening due to the point-spread function (i.e., the Fourier transform of the CTF), the presence of inward curling on each CM subspace is greatly exacerbated. There are additional higher-order aberrations of the CTF relevant for high-resolution cryo-EM^63^ (such as astigmatism and beam tilt) which can be easily incorporated in the CTF of the double filter. While not explored here, these may further affect the spectral geometry if not accounted for properly.

### 3. Determine pairs of eigenvectors for CM parabolas

Specific techniques must next be applied to discover the set of conformational modes corresponding to each CM within each Ω_PD_ embedding, with the number of these sets defined by the dataset’s intrinsic dimensionality. However, as described in section SM-XI, there exists an initial uncertainty about how to best determine the intrinsic dimensionality of any given dataset^64,65^ (i.e., the number of CMs present to search for), for which both an evaluation of the eigenvalue spectrum (for both PCA and DM) and bandwidth estimation strategy (for DM) have proven unsatisfactory for our purposes. To circumvent this uncertainty, we have introduced an elimination procedure to locate subspaces where conformational information most likely resides, and based on information gleaned from those findings, eliminate unsuitable sub-spaces from further study.

Our analysis demonstrated that the minimum information required during this discovery was the attainment of the lowest-order Chebyshev polynomial (*T*_2_) for each degree of freedom, with all other features in the embedding irrelevant to our needs. For any number of CMs defined by the dataset’s intrinsic dimensionality, the point-cloud resemblance to these Chebyshev polynomials can be found spanning specific 2D subspaces. However, care must still be taken to prevent overfitting, since blindly performing the subsequent steps in this framework on subspaces that do not unequivocally represent one of the dataset’s CMs (note that there are principally many such subspaces, such as those housing harmonics) can produce nonsensical conformational movies. For instance, we have provided several 2D NLSA movie outputs corresponding to independent degrees of freedom in movie M8 showcasing physically impossible motions (further descriptions are provided in section SM-XIX). Likewise, we have demonstrated how sets of parabolic harmonics for each CM naturally exist in all Ω_PD_ embeddings and, due to the overlapping nature of their point clouds, are unviable for mapping conformational information.

In contrast, the ESPER algorithm automatically fits each 2D subspace with parabolas, and signals the best-fit subspace for each eigenvector row for use in subsequent steps. Further, the eigenvector indices of these best-fit subspaces can be used to procedurally eliminate the use of CM harmonics, as is described in section SM-XVI. The availability and use of this information acts to circumvent the potential for contextual confusion in outputs, and is a direct remedy to one of the most prominent uncertainties^11^ in the founding ManifoldEM framework (see section SM-XIX). However, in practice, the ability to strategically avoid harmonics is only applicable up to the number of CMs present having pronounced geometric structure viable for reliable parabolic fits. It i s likely that more advanced strategies could be applied to optimize our routine to additionally eliminate a larger subset of higher-order harmonics. Finally, it is important to carefully examine the 2D movies produced for each potential CM by eye. The emergence of nonsensical (i.e., physically impossible) patterns should be seen as a strong indicator for overfitting of the Ω_PD_ embedding. In a similar vein, the possibility of underfitting (analyzing too few CM subspaces) must also be taken into account, though it carries significantly less risks.

### 4. Find optimal rotations for the eigenfunctions

As we have seen to some extent in every embedding explored, depending on the choice of PD, the conformational modes previously discovered can appear mis-aligned with the plane of their 2D subspace. As shown in section SM-XIV, this effect arises from the introduction of altered, foreshortened distances in each 2D projection (i.e., PD disparity), and could be countered via rotation of specific eigenvectors to ease retrieval of the CM. Given this knowledge, we have presented an automated procedure for the discovery of these optimal rotation angles for each PD. Despite its remarkable performance on our four 126-PD datasets (Fig. 11 and Fig. S11), we believe there is still room for this technique to be further developed in future works so as to account for rare events such as complex boundary conditions (e.g., due to steric hindrance between domains, as discussed in section SM-XV) as well as datasets with *n >* 2 degrees of freedom (see section SM-XVI). In the prior case, the use of additional rotation operators may be required, creating a more complex collection of decisions, possibly well-suited for a maximum-likelihood approach. Furthermore, for noisier, less-structured embeddings, we have observed that the 2D histogram method may provide close but not perfectly-aligned counter-rotations. It is possible that this method can be further elaborated by using additional eigenfunctions, which would allow the fitting with parabolic sheets instead of—or perhaps alongside with— histogram distributions.

These findings have directly established a new set of requirements in the types of datasets most viable for PD-manifold studies of conformational continuum. If no geometric form can be deciphered in all or even a significant number of Ω _PD_ embeddings, we claim it is effectively impossible to find the proper counter-rotation such that all PD eigenbases are displayed in a common coordinate system defined by CMs. We have further observed that the inability to align eigenfunctions onto this common eigenbasis can to varying extents subvert the quality of 2D movies and occupancy maps (see PD_33_ in movie M8), and thus ultimately the quality of the final 3D reconstructions. This fact explains several of the limitations^11^ recently documented in the founding PD-manifold approach^1,10^, since the previous ManifoldEM approach does not account for the tendency of Ω_PD_ embeddings to be unpredictably unaligned from a common coordinate system (e.g., Fig. S28). For a more detailed analysis of limitations and uncertainties observed, see section SM-XIX. Thus, in order to maximize fidelity of final outputs with ground truth, a dataset must first fulfill a minimum set of requirements that ultimately provide a well-structured spectral geometry when embedded. The performance of ESPER hinges on the presence of this geometric information, and as we have shown and will continue to discuss in the sections that follow, a great number of benefits emerge when it is available.

### 5. Fit and partition each CM point cloud

Once the set of conformational modes are identified and counter-rotated for each Ω_PD_ embedding, a further obstacle is encountered in the heightened variability of these point clouds (Fig. 7) – which vary in average thickness, density, length and type of trajectory, as well as spread of data points – depending on CM and PD. These complications were addressed through use of a robust automation strategy, including least-squares fitting and area-based partitioning of the essential subspaces. Overall, the area-based procedures employed—including the ball tree and alpha shape frameworks—are strongly affected by large changes in parameters and thus may require initial supervision on externally-obtained datasets. For the purpose of this study, we investigated over 500 subspaces, with each of our parameters broadly tuned for robust, high-quality performance on the SS_2_ CM_1_ and CM_2_ subspaces from data-type II and III. Detailed notes on these procedures have been provided in our published repository^37^, which includes comments describing less-significant decisions not explicitly noted in our main text. As a final note, we analyzed the outputs of these procedures on both the parabolic {Ψ_*i*_, Ψ_*j*_} and transformed {Φ_*i*_, Φ_*j*_} CM subspaces, and found consistently better agreement with ground-truth occupancies in the latter kind.

### 6. Bin points along straightened trajectory and integrate images

After achieving these prior conditions, points (one corresponding to each image) projected onto their respective least-squares fit must be binned, with bins created via the aforementioned area-based approach. Each of these bins represents one of the system’s unique states along the CM corresponding to the current 2D subspace. While we used a bin size as informed by our ground-truth knowledge to enable a direct comparison between inputs and outputs, it is another issue entirely to decide on the proper bin size for real-world data. As detailed in section SM-XVII, note that the bin size effectively controls the precision to which we can locate each point in the higher-dimensional surface, and it influences the range of images falling within each state that we group together as virtually identical for means of our final outputs. Naturally, the use of this optimal value should maximize the amount of information ascertainable in our system. In theory, we desire a minimum number of snapshots in the lowest-occupancy bin, such that every frame of the subsequent movie has significant content; e.g., as possibly defined via both the average SNR of each image and the number of images in the lowest-occupancy bin.

For each CM in a given dataset (regardless of intrinsic dimensionality: 1, 2, and 3 were tried), we were able to project the set of our images onto curves determined by least-squares fits, organize them into bins, and produce near-perfect 2D movies displaying each conformational motion independently of all others (movies M2 and M3). The corresponding occupancy maps showed a favorable agreement with our ground-truth knowledge (Fig. 10-G, Fig. 11, Fig. S11), and were significantly more accurate than those produced by NLSA for all datasets explored (movie M8). Overall, the procurement of accurate occupancy maps for each PD was by far the most trying endeavor, requiring a robust workflow able to account for large variations in a wide range of manifold characteristics. While it is very easy to split the CM subspace crudely into only two or a handful of states, as the number of states requested is increased, the degree of sophistication in mapping and segmenting each point cloud must also increase in turn. As previously noted, when dealing with experimental data, these occupancies contain vital information for the energetics of the ensemble, and are directly linked to the free-energy landscape spanned by the conformational degrees of freedom. We will return to this topic in the next section when discussing the final occupancy maps generated via integration of CM content across all PDs on *S*^2^.

### 7. Identify motion type and sense for each CM 2D movie

With the acquisition of each isolated conformational motion for each PD, lastly in such a framework, these CMs must be stitched together^1,56^ using equivalent CM information from each PD across *S*^2^. For our workflow, this procedure required manually solving two subproblems: (i) identifying each set of CMs across *S*^2^ (e.g., such that CM_1_ in a given PD is matched with CM_1_ in all other PDs, etc.) and (ii) identifying the sense of each CM in each PD. We draw attention again to the external, comprehensive method developed to solve this problem with heightened accuracy, which uses optical flow and belief propagation algorithms^56^, and strongly advise against making such assignments arduously by hand.

### 8. Compile CMs on S^2^ for free-energy landscape and 3D movies

After CMs have been properly identified and matched among all PDs, the indices of points (designating images) and related statistics can be organized to produce an *n*-dimensional occupancy map, with the images assigned to each *n*-dimensional bin therein accumulated to form a corresponding 3D density map. Once these previous steps were performed for our dataset, we compared the state assignments of all images from *S*^2^ to their ground-truth indices. We found that the majority of images in each bin (state) were correctly assigned, with each bin also encompassing a small set of images that were actually ground-truth members of neighboring bins (Fig. 11 and Fig. S11).

Our final 2D occupancy map (also seen in Fig. 11), formed by the intersection of snapshots in both 1D occupancy distributions, showed similar consonance with ground-truth expectations (Fig. S13). As a result of these accurate occupancy assignments, we ultimately observed a remarkable fidelity between each of the 400 reconstructed 3D density maps obtained by ESPER (movie M4) and their respective ground-truth 3D density maps. In addition, the resolution of all volumes were close to 3 Å, matching the resolution of the ground-truth structures. Overall, the small spread of states into neighboring bins appeared to have only a marginal effect on the quality of final 3D density maps. Through our analysis, it is apparent that the majority of aberrant information in each bin (corresponding to neighboring states) was effectively averaged out, such that only the conformational-variation signals corresponding to the current state dominated in each subsequently-generated 3D reconstruction. These findings were further supported quantitatively by an assessment of the structural fidelity of ESPER outputs with ground-truth atomic-coordinate structures and 3D electron density maps using Q-score calculations on residues and FSC curves, respectively (see section SM-XX).

As a final note, since our method retains the original image content for each image index assigned to a given bin, it is possible to further improve these image assignments after they have been generated by our sub-space fitting routines. Our ability to leverage the final image content to further improve 3D density maps and corresponding occupancy distributions stands in contrast to the founding PD-manifold approach^1^, which relies on histogram equalization (technically *histogram matching*) to match the occupancy distributions across PDs. Although not pursued here, one possible way is a maximum-likelihood approach aimed at comparing images within each bin and reassigning erroneously-assigned subpopulations into the neighboring bin in which they most likely belong. To note, a maximum-likelihood approach does already exist that aims to extract such granular conformational heterogeneity^66^, as does a method based on neural networks^67^.

## CONCLUSIONS

An ensemble of synthetic cryo-EM projections of the Hsp90 protein undergoing quasi-continuous conformational changes has been generated as an exemplary ground-truth model to determine how macromolecular motions appear in low-dimensional representations of their respective dataset using the PD-manifold approach. Based on knowledge obtained by this analysis, we have introduced a novel, unsupervised workflow with several enhancements that substantially improve the ManifoldEM framework and its ability to recover continuous conformations from single-particle cryo-EM data. These include essential eigenvector rotations to consistently align the spectral geometry of all Ω_PD_ embeddings, use of the original cryo-EM images to form high-quality 2D and 3D movies, and efficient strategies for building multidimensional free-energy landscapes. Along with the introduction of this workflow, we have also pointed out challenges, fundamental limitations and uncertainties that emerge in geometric machine learning of heterogeneous cryo-EM data. Finally, we hope that the insights gained from these machine-learning heuristics will be useful not only in cryo-EM, but also in the development of other techniques aimed at untangling complex systems exercising multiple, continuous degrees of freedom.

## ACKNOWLEDGMENTS

We would like to express our gratitude to Abbas Ourmazd, Ali Dashti and Ghoncheh Mashayekhi for their insights and expertise, providing a number of stimulating conversations throughout this development. This research was supported by the National Institutions of Health Grants GM29169 and GM55440 (to J.F.) and by the U.S. National Science Foundation Award No. STC1231306 (to P.S.).

## CONTRIBUTIONS

Author contributions listed alphabetically below. All authors reviewed the final manuscript.

- Conceptualization: ES, JF
- Formal analysis: ES
- Methodology and software: ES, FAR, PS, SM
- Validation: ES, FAR, PS
- Direction and project administration: JF
- Manuscript draft: ES
- Manuscript review and editing: ES, FAR, JF, PS, SM

Detailed contribution notes for the ESPER algorithm suite have been supplied in the header of all scripts in our online repository^37^.

## DATA AVAILABILITY STATEMENT

The Python software repository^37^ is available at the following link: https://github.com/evanseitz/cryoEM_ESPER This repository includes several pristine image stacks and algorithms to modify them with different occupancy assignments, CTF and SNR. In addition, a collection of pre-generated manifolds are supplied for immediate use in computations employing the eigenfunction realignment and subspace partitioning algorithms presented here.

## Appendix: Description of symbols and abbreviations

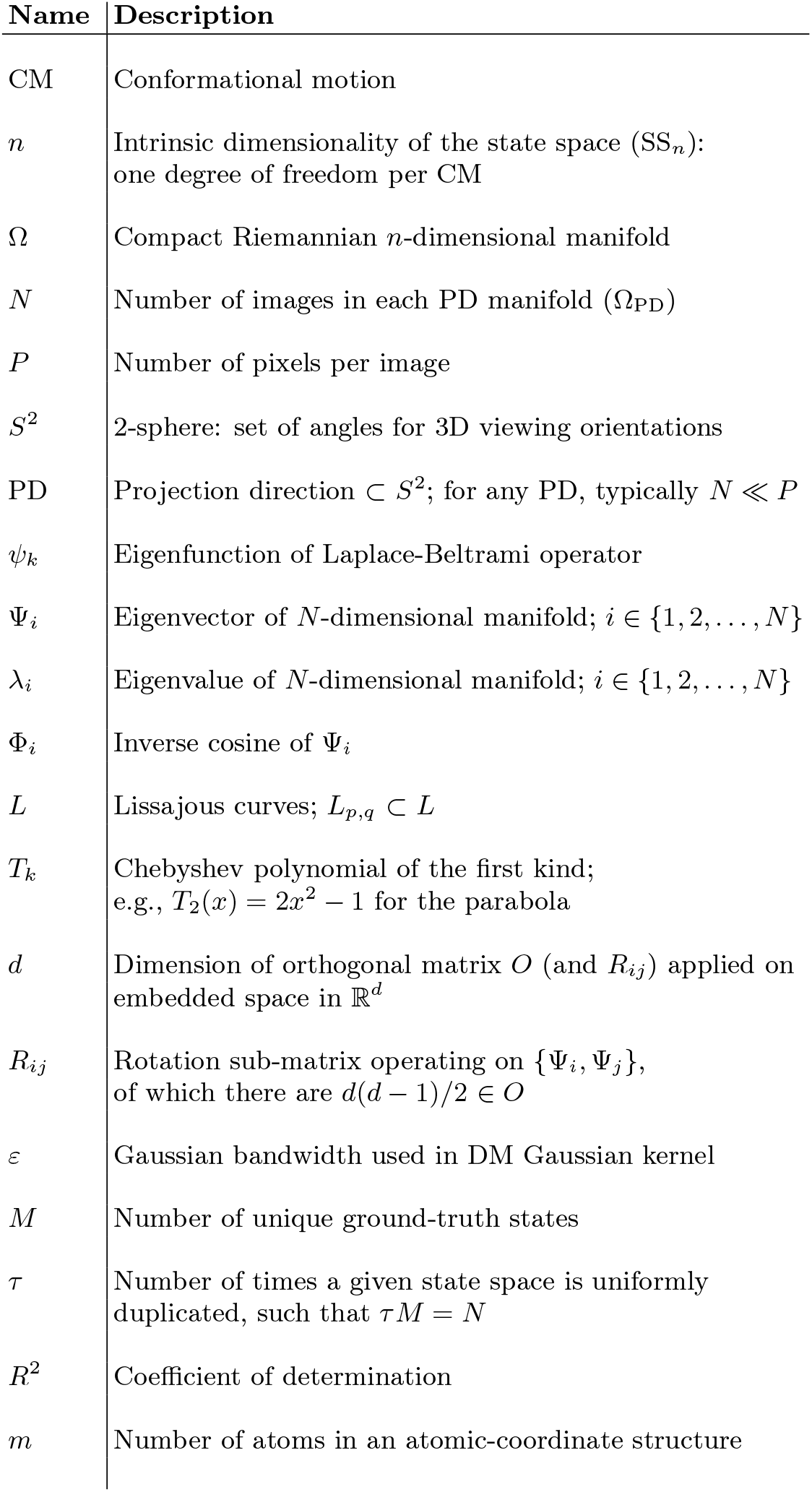

## Footnotes

† A tabulated description of symbols and abbreviations used throughout this document is available in the Appendix.

† There is a wide range of nomenclature used here between fields and, in some instances, works by the same author. The following are interchangeable: *conformational motions*; *conformational coordinates*; *reaction coordinates*; *collective motion coordinates*.

‡ All supplementary material sections will be referenced throughout this document in form SM-{Roman numeral}. The ordering of sections in SM is arranged to form a cohesive narrative, separate from the order each section is introduced in our main text.

