## Supplementary Material for "Geometric machine learning informed by ground truth: Recovery of conformational continuum from single-particle cryo-EM data of biomolecules"

Evan Seitz,[1] Francisco Acosta-Reyes,[2] Suvrajit Maji,[2] Peter Schwander*,[3] and Joachim Frank*[1,2]
[1)] *Columbia University, Department of Biological Sciences, New York, NY 10032, USA*
[2)] *Columbia University Medical Center, Department of Biochemistry and Molecular Biophysics, New York, NY 10032*
[3)] *University of Wisconsin-Milwaukee, Department of Physics, 3135 N. Maryland Ave, Milwaukee, WI 53211, USA*

#### SM-I. SIMULATION OF CRYO-EM ENSEMBLES FROM ATOMIC MODELS

This section provides additional information on the construction of our synthetic data, with more detailed documentation[1] and a repository of code for generating custom datasets available online[2]. In the time since its conception, this synthetic framework[1,2] has already been used as a benchmark in one external cryo-EM study on free-energy landscapes[3]. Here we repeat much of the previous presentation of our methodology to provide a description of the atomic-coordinate displacements between states for the specific case of our Hsp90 model with $n = 2$ exercising two conformational motions ($CM_1$ and $CM_2$). These CMs were designed so as to be fully decoupled from each other, such that no overlap occurs between the two sets of distinct atomic displacements (Fig. S1).

Atomic manipulations of the original PDB[4] coordinate file were done using PyMOL[5]. For $CM_1$, the chain A arm was rotated outwards (directly away from chain B) from its central hinge at residue 677, in 1° increments until a series of 20 rotational states were defined (Fig. S2-A). For each of these 20 $CM_1$ states, all residues above chain B's elbow (residues 12-442) were then rotated perpendicularly to the $CM_1$ motions in 2° increments until a series of 20 rotational states were defined for $CM_2$ (Fig. S2-B). These operations resulted in the creation of a total of 400 unique conformational states. The ensemble of these states can be organized in a 20×20 state space, where each entry represents one of the possible combinations of $CM_1$ and $CM_2$. This state space represents our synthetic model's complete ensemble of physically allowable conformations (i.e., the quasi-continuum). The specific size of the state space (400 states) was chosen based on the relative scale of the protein and range of its motions, providing approximately 3 Å and 2 Å gaps between states over a total arc length of 60 Å and 40 Å for $CM_1$ and $CM_2$, respectively (as visualized in Fig. S3). To note, geometry correction and energy minimization of generated states were skipped to avoid introduction of unintentional coupling of CMs.

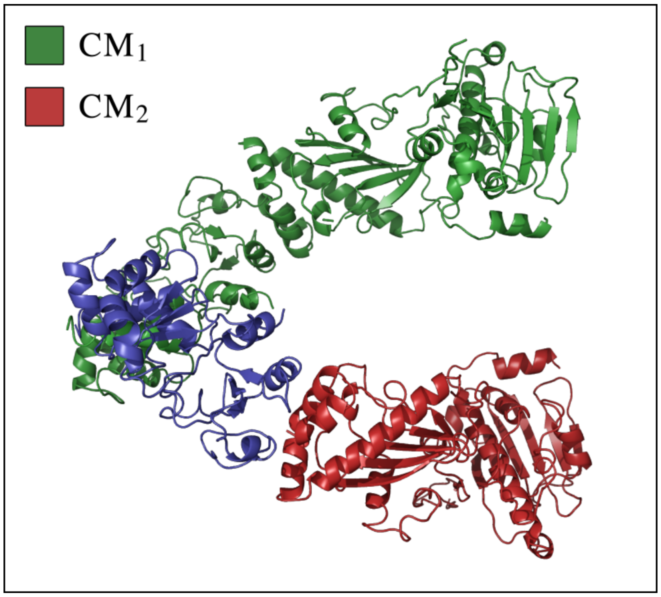


FIG. S1: Cartoon representation of atomic coordinates for state 20_01 with residues demarcated for $CM_1$ (green) and $CM_2$ (red). The $CM_1$ central hinge can be found near the intersection of the blue and green regions at residue 674, while the $CM_2$ hinge is found near the intersection of the blue and red regions at residue 443. To note, for $SS_2$, the residues making up the blue region are immobile throughout the entire state space.

These 400 structures represented by PDB files were then each transformed into 3D electron density maps (EDMs, formatted as MRC volume files) with a sampling rate of 1 Å per voxel and simulated resolution of 3 Å using the EMAN2[7] module `e2pdb2mrc`, where each atom is represented by a Gaussian with radius defined by the appropriate number of electrons (Fig. S3). Projections of each 3D density map can then be obtained via standard parallel line integrals along selected directions so as to simulate images generated in the transmission electron microscope (TEM) operated in the bright-field mode[8]. This core framework is used as a basis for the creation of images that are noisy duplicates or, finally, incorporate a CTF in their simulation:

- **Data-type I**: one copy of each state with pure signal and no CTF
- **Data-type II**: uniformly duplicated states with experimentally-relevant SNR
- **Data-type III**: uniformly duplicated states with experimentally-relevant SNR and CTF

An additional alteration on Data-type III is applied for use in our *final analysis*, namely the introduction of a nonuniform occupancy map.

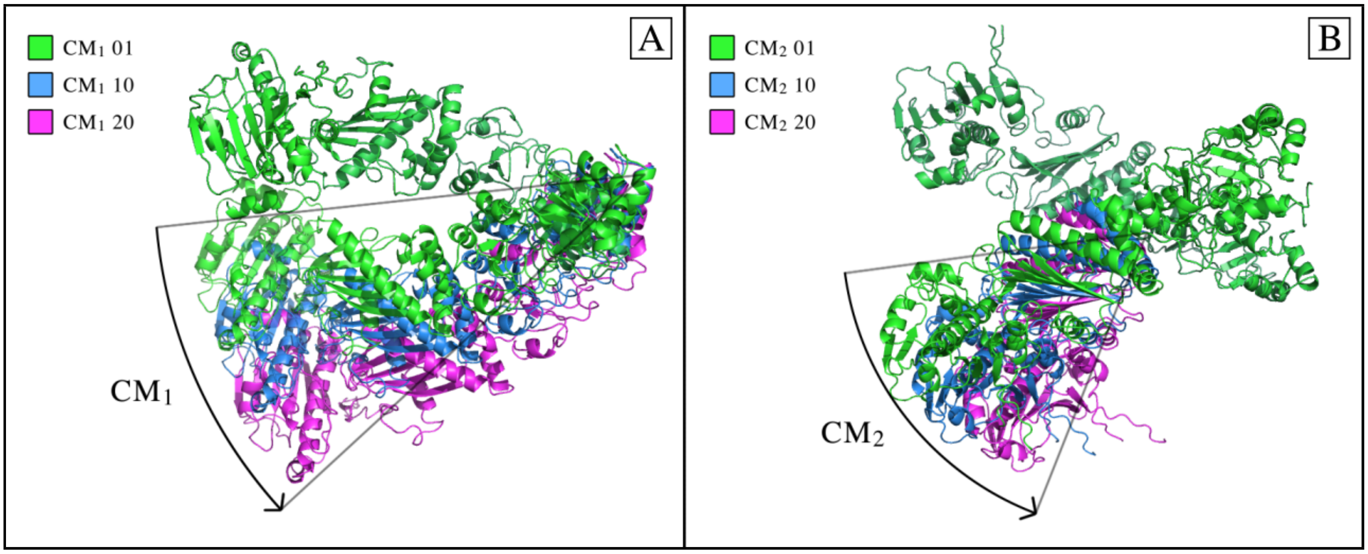


FIG. S2: In [A], a comparison of the $CM_1$ first, middle and final state is shown. Likewise, in [B], a comparison of the $CM_2$ first, middle and final state is shown. The $CM_2$ rotation was performed perpendicularly to the $CM_1$ rotational hinge where only those atoms above chain B's elbow-region were displaced. As a note, to remove the potential for overlapping residues within certain states, residues 1-11 (making up a loose tail) were removed from both chain A and B.

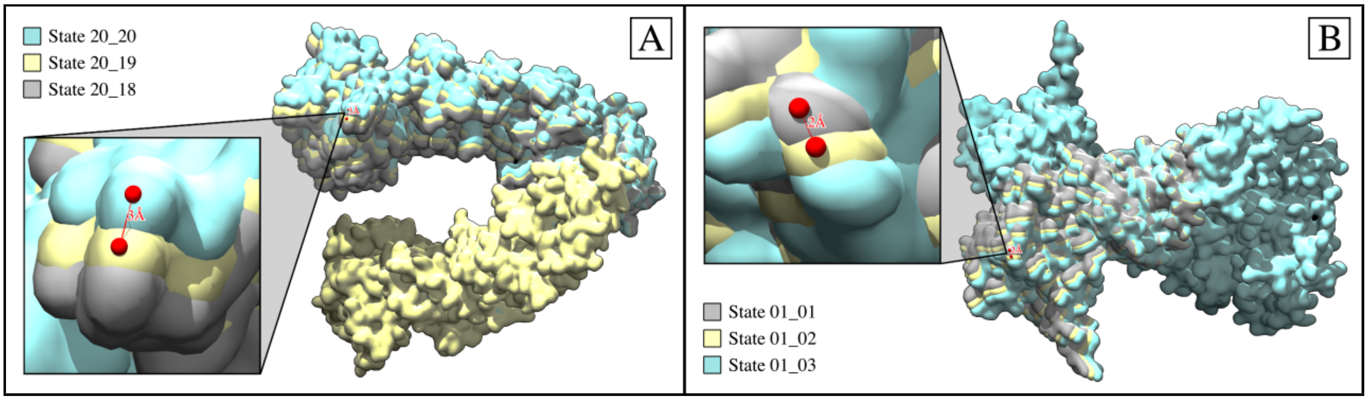


FIG. S3: In [A], a volumetric overlay of the first three rotational states of $CM_1$ (state 01_01, state 02_01, state 03_01) is presented, visualized as electron density maps (MRC format) via Chimera[6]. As can be seen, only the upper arm (chain A) has been rotationally displaced along $CM_1$, with 3 Å gaps measured between each consecutive state at the peripheral ends of this rotated region. In [B], a volumetric overlay of the last three rotational states of $CM_2$ (state 20_18, state 20_19, state 20_20) is shown. Only the upper region of chain B (above the elbow) has been rotationally displaced along $CM_2$, with 2 Å gaps measured between each consecutive state at the peripheral ends of this rotated region. As a note, the EMAN2 module e2pdb2mrc excludes calculations such as atomic form factors and molecular orbitals. While more accurate maps can be constructed using other methods, this does not affect the results of our heuristic analysis.

### SM-II. SIMULATING CRYO-EM ENSEMBLES FOR DATA-TYPE I

In the pristine scenario, for each electron density map within each state space ($SS_1$, $SS_2$ and $SS_3$), sets of images in five chosen projection directions (PDs) are first obtained. The first of these five PDs was chosen to be normal to the plane of the $CM_1$ rotation, such that all $CM_1$ motions from that perspective only underwent changes in the plane of the projection. A similar choice was made for $PD_2$, which was projected into the plane of $CM_2$ motions, with the remaining three PDs chosen at arbitrary positions in angular space. All projections were generated using the EMAN2[9] module `e2project3d` with no CTF applied, so as to maximally conserve the information contained in each projected EDM. (See movie M1 for a conformational animation of the molecule as viewed in each of these five PDs). For each state space, the set of these images as generated from one of these five PDs forms a high-dimensional manifold $\Omega_{\text{PD}}$. In all, this initial procedure resulted in the creation of 400×5 unique 2D images for each of our 400 3D density maps.

As pointed out in the main text, it is important to describe the order in which the states for each SS are indexed, with this ordering repeatedly visualized by color maps throughout this work and ultimately used to locate each state's coordinates within the embedded manifold. The ordering of $SS_1$ is trivial, with states following a sequence showing $CM_1$ transition from closed to open ($CM_1\{1\} \rightarrow CM_1\{20\}$; as described by the "arm" motion in Fig. S1-S3), while the ordering of $SS_2$ and $SS_3$ follow a clockwork pattern. For $SS_2$, the ordering of the states of $CM_2$ progresses like the second hand of a clock, with each progression from $CM_2\{1\} \rightarrow CM_2\{20\}$ iterating $CM_1$ (akin to the minute hand) forward once.

Thus, of the 400 states in $SS_2$, the first index corresponds to state $CM_1\{1\}\_CM_2\{1\}$, the second to state $CM_1\{1\}\_CM_2\{2\}$, the 21$^{\text{st}}$ to state $CM_1\{2\}\_CM_2\{1\}$, and the 400$^{\text{th}}$ to state $CM_1\{20\}\_CM_2\{20\}$. A similar pattern holds for $SS_3$, which now additionally includes an hour hand in this analogy ($CM_1$), followed by a minute hand for $CM_2$ and a second hand for $CM_3$. An animation of the molecule transforming through these sequences has been provided in movie M1. Of course, the resulting manifold embedding is independent of the ordering of the data points[10]; we have merely chosen one such ordering that heightens our awareness of trends in the subsequent outputs.

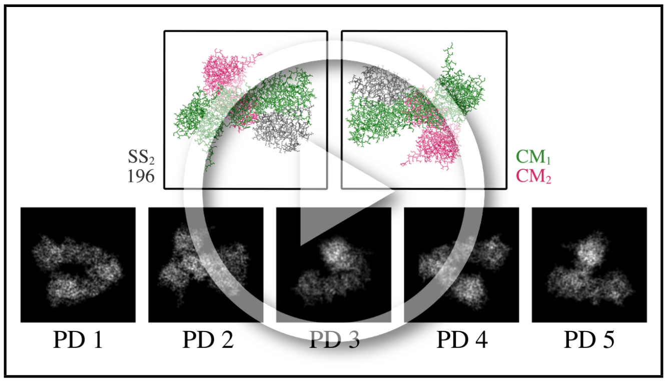


MOV. M1: An animation cycling through the 400 $SS_2$ states as represented by the Hsp90 atomic structures (top) and corresponding PDs (bottom), as analyzed in the $SS_2$ section of Results. The ordering of these states chronologically follows the indices described using our second hand and minute hand analogy. `https://www.dropbox.com/s/2qzhrzu0jzc8bbu/M1_SS2_GroundTruth.mp4?dl=0`

### SM-III. ANALYSIS OF MANIFOLD SUBSPACES FOR DATA-TYPE I

The following figures display several subspaces obtained from embeddings of pristine (noise-free) datasets. First, in Fig. S4, we provide an example of the inverse-cosine mapping $\{\Psi_i, \Psi_j\} \mapsto \{\Phi_i, \Phi_j\}$, as described in the main text. Next, we show the location of each set of Chebyshev polynomials within several exemplary subspaces, and demonstrate how each set corresponds to a unique conformational motion.

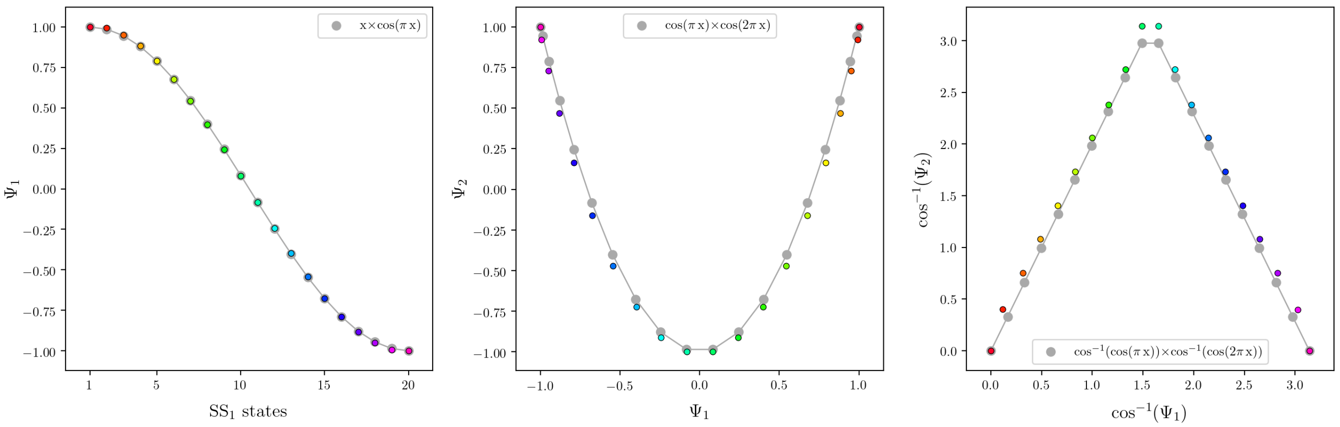


FIG. S4: For each of the three subplots, (1) analytical cosine functions are colored in gray and overlaid with (2) the coordinates of corresponding $PD_1$ eigenvectors, shown in different vibrant colors denoting the $SS_1$ sequence of states. To compare these two representations within the same coordinate system, the DM eigenvectors have been scaled to match the range of the analytical cosines. Specifically, for $x \in [0,1]$, 20 $x$-values were used to generate each $\cos(k\pi x)$ function and subsequently scaled to match the number of $SS_1$ states; i.e., $x' \in [1,20]$. Similarly, each eigenvector generated by DM was scaled to match the range of each cosine, such that $\Psi_k \in [-1,1]$.

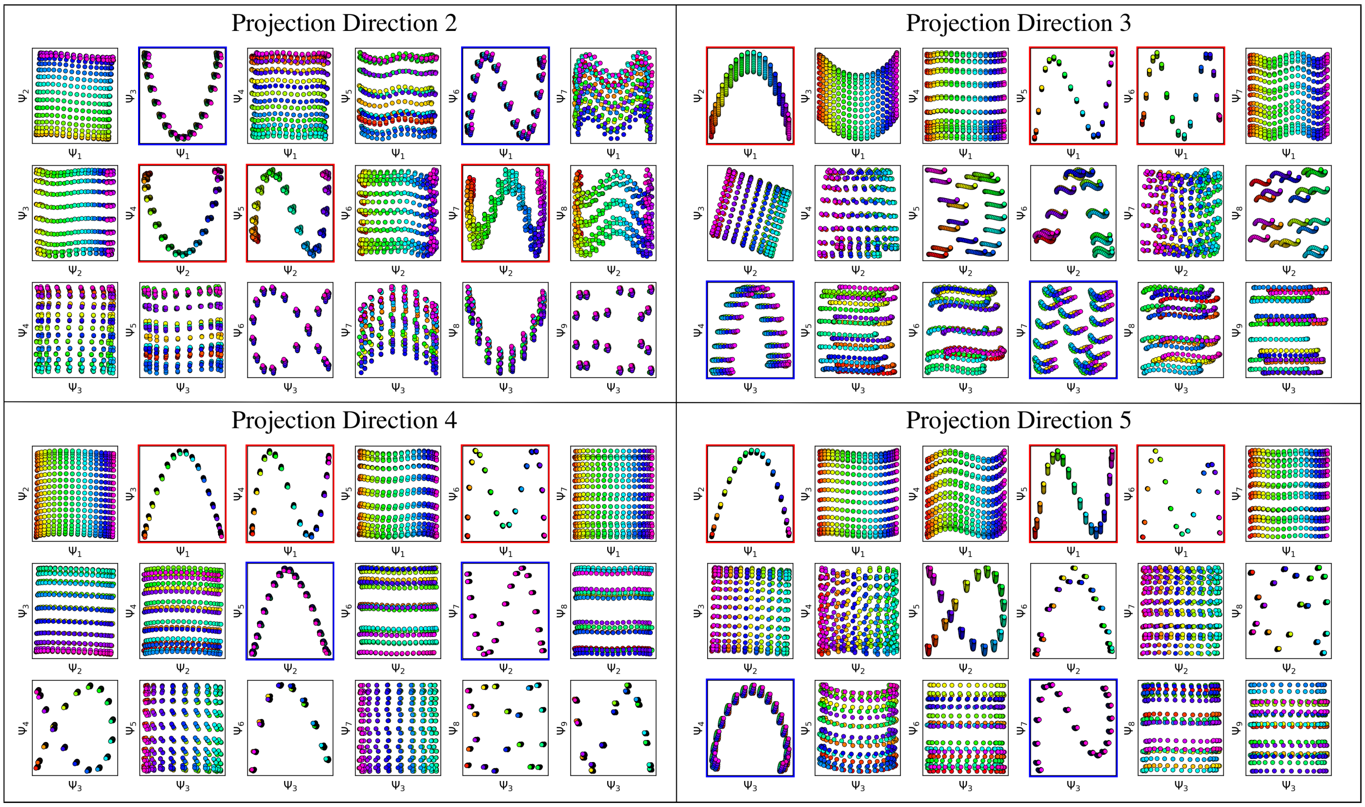


FIG. S5: A similar presentation as is shown in Fig. 4 for the remaining four PDs. Again, for each PD, red and blue boxes indicate the unique set of modes for $CM_1$ (red) and $CM_2$ (blue), which are interspersed throughout each row in specific 2D subspaces obtained from their respective $N$-dimensional embeddings. Subspaces requiring eigenvector rotations (e.g., both parabolas in $PD_3$) and housing subtle boundary problems (e.g., the curling inwards of the point-cloud trajectory in $\{\Psi_3, \Psi_4\}$ of $PD_5$) can also be seen in certain 2D subspaces. Note that in $PD_2$, the hierarchy of CM information is actually reversed from those seen in the other 4 PDs, with the $CM_2$ Chebyshev polynomials instead present along $\{\Psi_1, \Psi_i\}$ combinations (in the first row) and $CM_1$ Chebyshev polynomials instead along $\{\Psi_2, \Psi_j\}$ combinations (in the second row). The ordering of sinusoidal sets in each PD is related to the magnitude of the CMs present as seen from the given viewing direction, and not necessarily defined by the CM undergoing the largest change in the ground-truth atomic structures. Therefore, since $CM_2$ is projected in the plane of $PD_2$, the magnitude of its apparent motion is visually greater than $CM_1$, hence the reversed ranking.

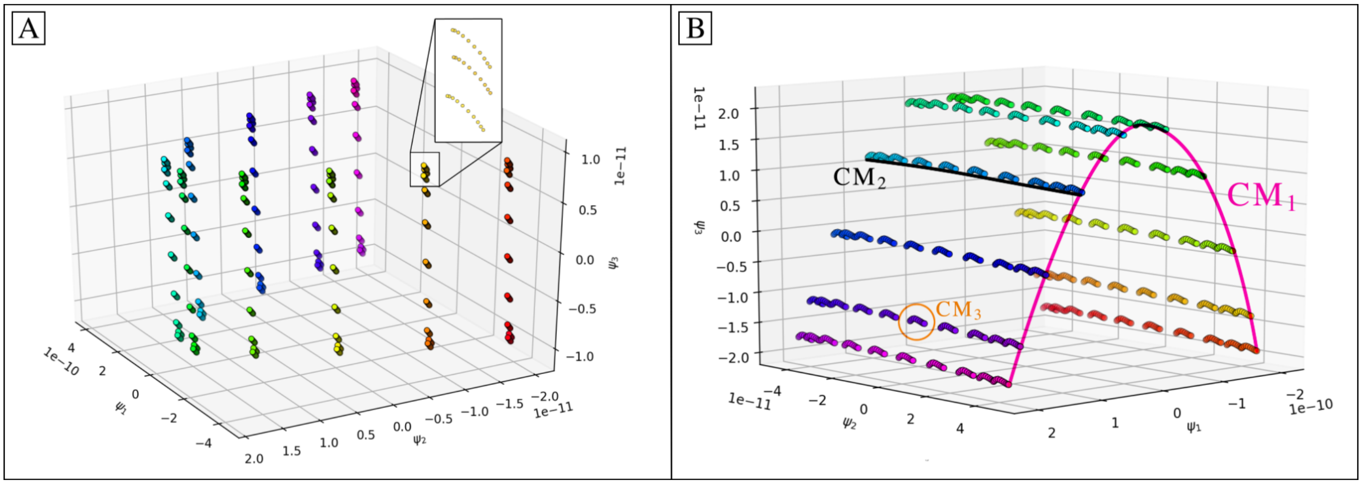


FIG. S6: Comparison of 3D subspaces obtained for images in $PD_5$ [A] and $PD_4$ [B] from $SS_3$ (1000 states arranged in a 10×10×10 state space). The inset in [A] illustrates the fractal pattern that emerges when more than two conformational motions are present, where $CM_3$ can be seen forming a series of mini-parabolas about each of the points in the larger 10×10 parabolic surface. In [B], the hierarchy of these structures is more pronounced, with exemplary CM trajectories depicted by the plotted lines. In section SM-XIV, we demonstrate that these patterns arise due to the additions of orthogonal cosines.

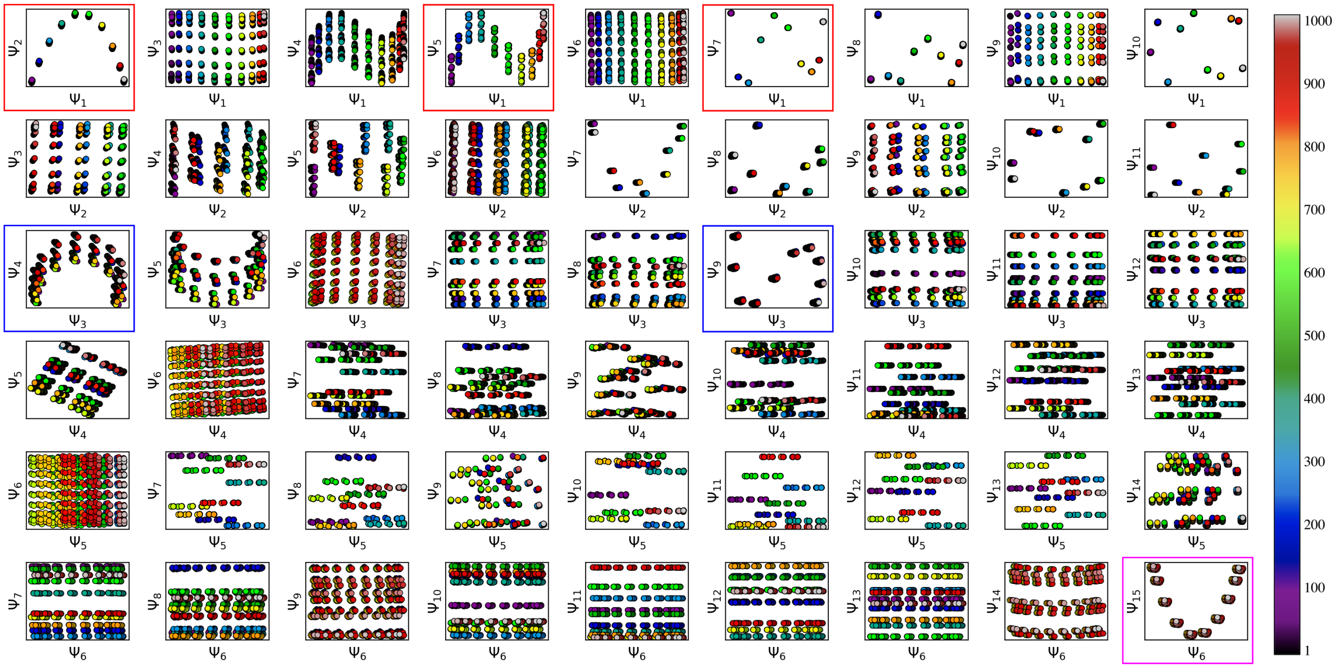

FIG. S7: A set of 2D subspaces projected from the $N$-dimensional embedding obtained from $PD_5$ in $SS_3$. The set of conformational modes corresponding to $CM_1$ is demarcated by the red boxes around interspersed $\Psi_1$ plots, and occupy specific $\{\Psi_1, \Psi_i\}$ combinations (where $i > 1$). Likewise, $CM_2$ and $CM_3$ are both separately represented by a set of their own conformational modes; demarcated by blue boxes around interspersed $\Psi_3$ plots and magenta boxes around interspersed $\Psi_6$ plots (of which only the first is displayed), respectively. Specifically, $CM_2$ modes span a set of $\{\Psi_3, \Psi_j\}$ combinations (where $j > 3$) while $CM_3$ modes span a set of $\{\Psi_6, \Psi_k\}$ combinations (where $k > 6$); with the eigenvector depth specified by the apparent span of each conformational motion as seen from the given PD. As expected, points on the trajectories defined by $CM_1$ modes follow along the full spectrum of colors (i.e., indices 1-1000), while $CM_2$ and $CM_3$ points cover a span of 100 and 10 colors, respectively. Note the presence of parabolic harmonics in neighboring rows (for each CM), which are characterized by the presence of $\alpha$-shaped trajectories (e.g., as seen in $\{\Psi_2, \Psi_5\}$, albeit slightly rotated here from the plane of its 2D subspace).

### SM-IV. SIMULATING CRYO-EM ENSEMBLES IN DATA-TYPE II

Data-type II uses images generated for data-type I as a base, from which images are uniformly duplicated $\tau$ times. Additive Gaussian noise is next applied to each image individually to grant it a specific SNR that is the same for all images in the set. We define the SNR by the ratio of each image's signal variance ($\sigma^2_{\text{signal}}$) to its noise variance ($\sigma^2_{\text{noise}}$)[8]. Here, signal represents the 2D region of pixels corresponding to the average area occupied by the macromolecule. This region is obtained by masking out all pixels within one standard deviation of each image's mean intensity value; in effect, excluding the approximately uniform-intensity background. This process was thus performed by first finding the mean pixel intensity ($\mu_{\text{signal}}$) and variance ($\sigma^2_{\text{signal}}$) of the *signal*, and then calculating $\sigma^2_{\text{noise}} = \sigma^2_{\text{signal}}$ / SNR. Using this parameter, we then apply additive Gaussian noise to each image in order to obtain an output image having the desired SNR. In this process, a sample from the Gaussian distribution is added to each pixel's intensity. Each resulting image was then normalized such that the average pixel intensity and standard deviation of pixel intensities was approximately 0 and 1, respectively.

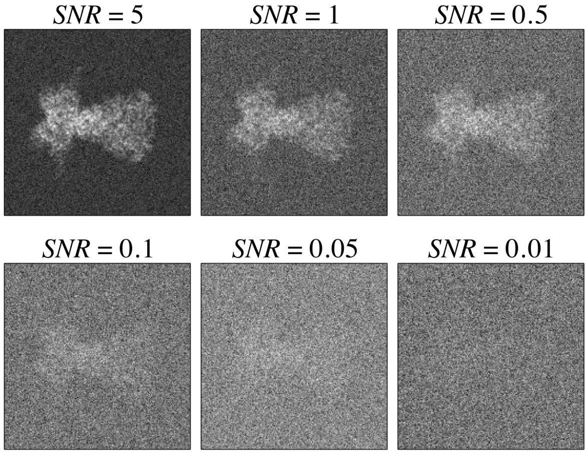


FIG. S8: First image in $SS_1$ from $PD_1$ with different values of SNR via additive Gaussian noise. As a note on experimental relevance, SNR = 0.1 has been previously established as a suitable choice for experimental SNR in images obtained by cryo-EM, and its low value can be attributed to the low contrast between macromolecules and their surrounding ice, as well as the limited electron dose required to avoid radiation damage[11].

### SM-V. ANALYSIS OF SUBSPACE FITTING AND PARTITIONING FOR DATA-TYPE II

In this section, we first present a set of 2D movies captured from $SS_2$, as seen in movie M2. We then analyze the robustness of recovering several conformational modes while varying $\tau$ values and SNR regimes. In Fig. S9, we investigate the robustness for the leading conformational modes (i.e., Chebyshev parabolas and higher-frequency oscillations occupying specific 2D subspaces of PCA embeddings) by fitting across a range of increasing $\tau$ values. The coefficient of determination ($R^2$) is calculated for each mode to provide a measure for the quality of its corresponding fit.

Fig. S10-A shows the $R^2$ trend for each mode as $\tau$ is incrementally increased. In Fig. S10-B, only the $R^2$ values for the first mode are similarly calculated across several SNR regimes. The asymptotic behavior of each plot is expected, demarcating regions along this trajectory where the intrinsic geometric structure of each mode is optimally reinforced against the background noise. By visual assessment of consecutive $\tau$-defined embeddings, for all asymptotic plots (those in both Fig. S10-A and Fig. S10-B), we found that a suitable value of $\tau_c$ could be estimated at approximately half the corresponding plot's maximum value of $R^2$. For example, in Fig. S10-A, the value of $\tau_c$ for the first mode is approximately 15; in agreement with the first emergence of a robust parabola seen in Fig. 6.

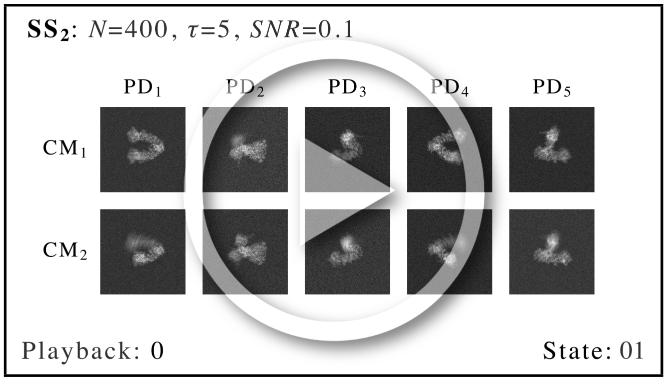


MOV. M2: Set of 2D movies captured along $SS_2$ subspaces, with each corresponding manifold generated from images with SNR of 0.1 and $\tau = 5$. As seen in both the first and second row, as a result of the integration procedure on each corresponding CM parabola, there is a significant difference in resolution between the desired CM and the CM orthogonal to it. For example, in the first row, the arm motion appears crystal clear in each PD, while the (orthogonal) elbow motion appears blurry; and vice versa for the second row. When 3D movies are eventually constructed from the full set of these 2D movies on $S^2$, pairwise information from both CMs is incorporated into each volume such that these anomalies resolve. `https://www.dropbox.com/s/z27lpl4orpli1xt/M2_2D_Movies_DT2.mp4?dl=0`

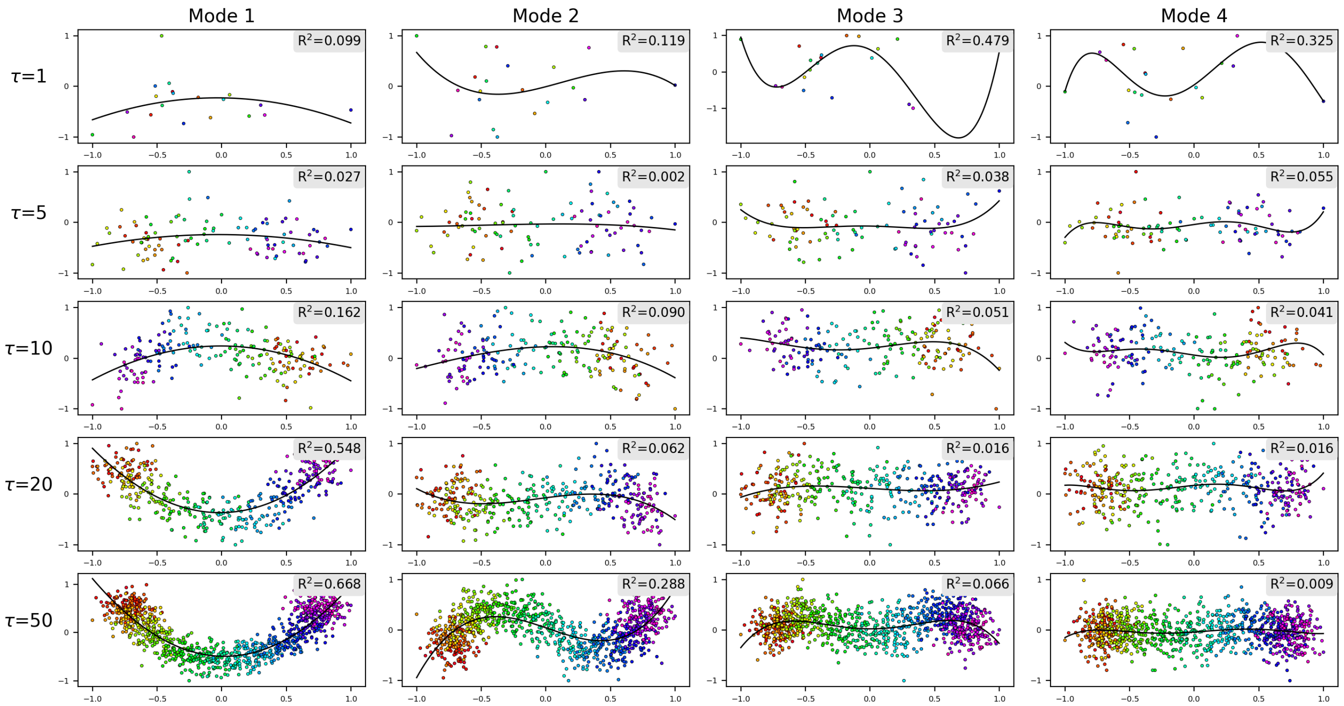


FIG. S9: Each of the above rows show a set of 2D subspaces from a $\tau M$-dimensional embedding (in $SS_1$ where $M = 20$, as obtained from an ensemble of images created with that row's given $\tau$-value and SNR = 0.1). Each 2D subspace within each row displays one of the CM's leading Chebyshev modes $m$ via $\{PC_1, PC_{m+1}\}$ (e.g., all 2D subspaces in the first column have $x$-axis and $y$-axis defined by $PC_1$ and $PC_2$, respectively). Note that each of these principal components has been scaled to have matching bounds (i.e., [-1, 1]). Lines of best fit are then computed with $R^2$ values recorded, as displayed in the corner of each subplot. As an aside, note the inconsistent orientation of each mode (e.g., upwards or downwards concavity for the parabolic mode), which is due to arbitrary eigenfunction signs which naturally arise during eigendecomposition[10].

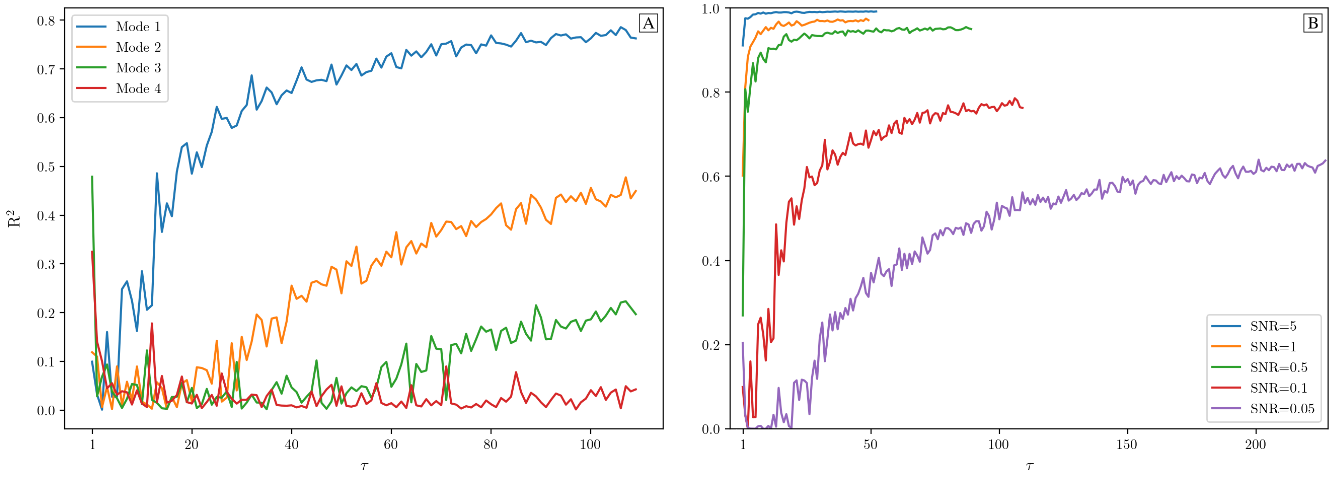


FIG. S10: In [A], the $R^2$ values are plotted for leading modes with constant SNR. The first mode corresponds to the parabolic Chebyshev mode defined via the projection $\{PC_1, PC_2\}$, with the second Chebyshev mode defined via $\{PC_1, PC_3\}$, et cetera (with exemplary best fits for all modes illustrated in Fig. S9). In [B], only the $R^2$ values for the first mode are shown while SNR is altered.

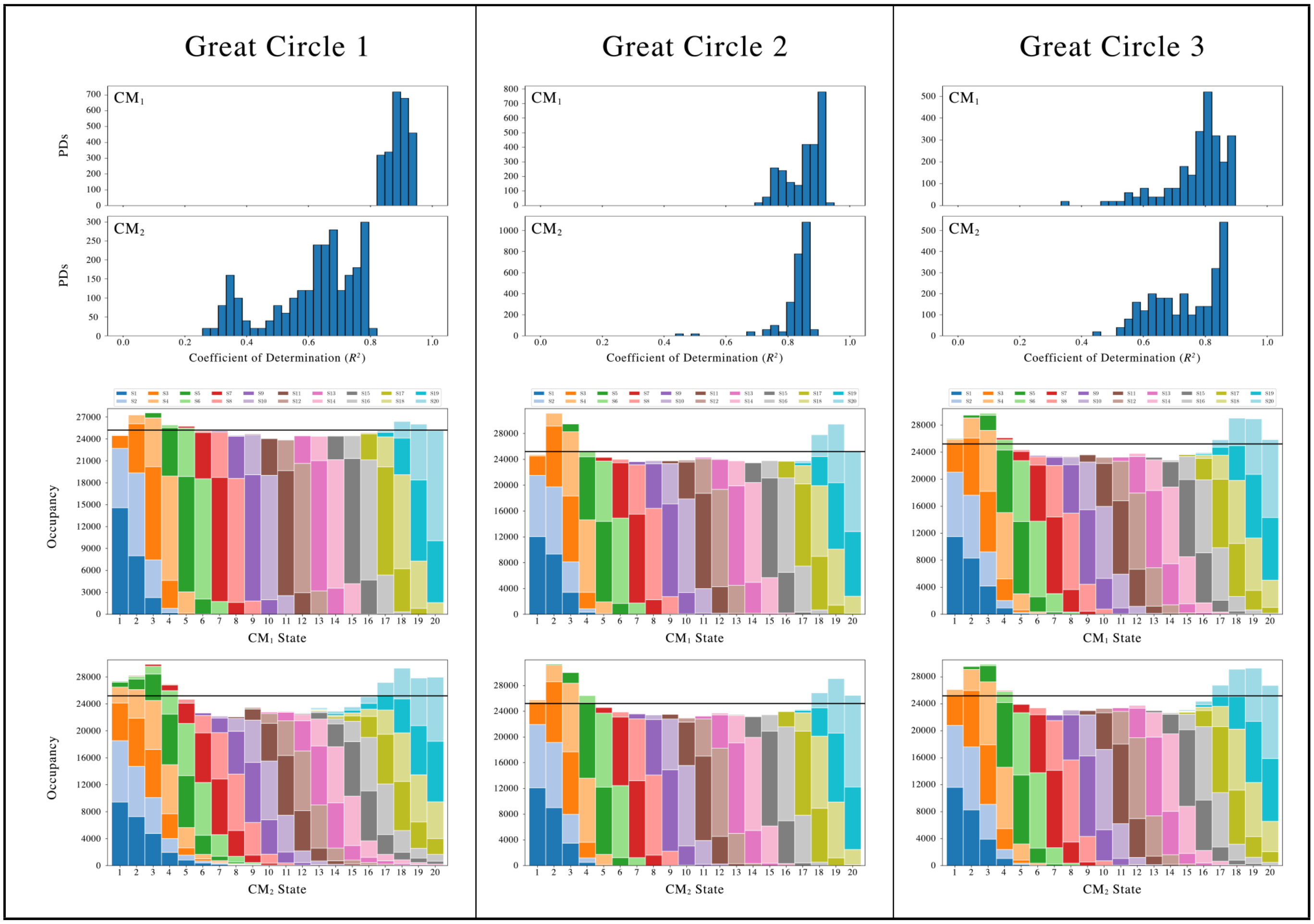


FIG. S11: Each column shows the final outputs of our workflow for one of three equispaced 126-PD great circle trajectories, with each trajectory computed in isolation from all others. To note, each of the great circles used are orthogonal to the other two. In every column, total occupancy maps for $CM_1$ and $CM_2$ from $SS_2$ with $\tau = 10$ are shown, as obtained by integration of the corresponding 20 bins for each CM (corrected for *sense*) in each of the 126 PDs. The total number of images as assigned to each state via our subspace fitting procedure is shown by the height of the 20 bars. Within each bar, the different colors represent how many of the assignments therein belonged to which ground-truth states (as seen in the legend). Note that the black, horizontal line demarcates the expected value of each bin for the ground-truth (flat) distribution; i.e., states $\times$ PDs $= 20 \times 10 \times 126 = 25,200$ images. The subplots above each of the great-circle occupancy maps show the corresponding histogram of $R^2$ scores for each CM as observed in the respective 126 PD-manifold subspaces.

### SM-VI. SIMULATING CRYO-EM ENSEMBLES FOR DATA-TYPE III

Data-type III again uses images generated for data-type I as a base. Similar to data-type II, $\tau = 10$ duplicates per state are first generated. However, instead of applying additive Gaussian noise to each image, as was done for data-type II, each image is first filtered by the electron microscope's contrast transfer function (CTF) in an experimentally relevant range. For this task, CTF is generated via the form CTF = $\sin(\chi) - A\cos(\chi)$, where $\chi = (-\pi\Delta z\lambda k^2 + \frac{1}{2}\pi C_s\lambda^3 k^4)$, $\Delta z$ is a defocus value randomly assigned in the interval [5000, 15000] Å (positive is underfocus); $\lambda$ is the wavelength of the electron (calculated via known microscope voltage); $k$ is the spatial frequency; $C_s$ is the spherical aberration; and $A$ denotes the fraction of amplitude to phase contrast. Once the CTF is generated with a choice of relevant microscopy parameters, it is next applied through scalar multiplication with the Fourier transform of the image, followed by an inverse Fourier transform of the product. After this procedure, additive Gaussian noise is uniquely applied to each image (SNR = 0.1) following previous protocol laid out for data-type II. Fig. S12 on the right provides a demonstration of this workflow, which further includes results of CTF correction. It should be noted that we use exactly the same CTF for initially modifying each image as we use later in our pipeline for CTF correction, making no allowance for experimental inaccuracy of CTF estimation.

Upon first performing these operations on the pristine states in $SS_2$, we found that, in the presence of CTF for certain PDs, gaps emerged between clusters of points, with points within each of these clusters representing an identical ground-truth state. For this experimentally most relevant dataset, it was clear that we needed better sampling of the quasi-continuum conditions. Thus, we shortened the total span of each CM while keeping the number of states constant, effectively increasing the density of states for data-type III. Specifically, we lowered the rotational distance between neighboring states in each CM until the corresponding point-cloud distributions appeared virtually continuous (i.e., without clusters or discernible gaps). The resulting RMSD between consecutive $CM_1$ states (e.g., state 01_01 and state 02_01) and $CM_2$ states (e.g., state 01_01 and state 01_02) was 0.4 Å each. In comparison to previous atomic relations (see caption of Fig. S3), the distance between the peripheral atoms of adjacent $CM_1$ states decreased to 1 Å and for $CM_2$ states to 1.5 Å.

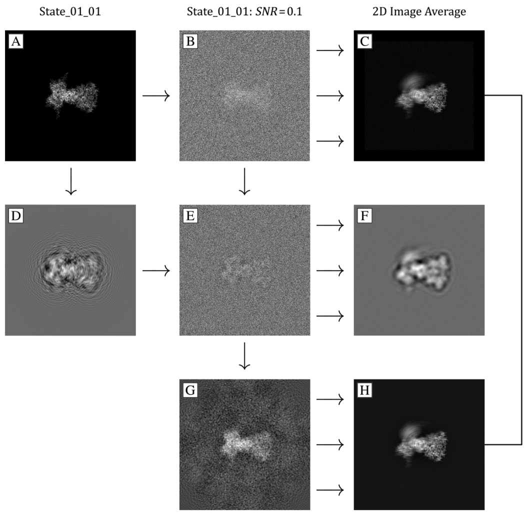


FIG. S12: In the first column, image [A] is a 2D projection of state 01_01 of $PD_2$ taken without CTF parameters using the EMAN2[9] module `e2project3d`. Image [D] is that same 2D projection but generated with CTF signal using the following parameter values: defocus 13,500 Å; amplitude contrast ratio 0.1; spherical aberration 2.7 mm; and voltage 300 kV. For both the first and second row, there are 399 other images each, created with respective attributes corresponding to the remaining $SS_2$ states. All of the 399 images generated similar to [D] were likewise given a random defocus value in the interval [5000, 15000] Å. Image [B] and [E] are the results of adding Gaussian noise to each image [A] and [D], respectively, such that the SNR becomes 0.1. As depicted with three cascaded arrows in the diagram, image [C] is the 2D average of all states generated similar to image [B], and likewise for the relationship of image [F] to image [E]. The image size is 320 × 320 pixels, with this decision guided by [D] such that the broadening of the point-response function stays within the image bounds. Image [G] is the CTF-corrected version of image [E], as depicted in previous work[8] using a Wiener filter with SNR = 0.1 and exact assignment of known CTF parameters. Finally, image [H] is the 2D average of all CTF-corrected images similar to image [G]. Note the similarity of [H] with [C], which is a result of filling the missing CTF zero-crossings during the averaging of all CTF-corrected images.

### SM-VII. OCCUPANCY ASSIGNMENTS FOR FINAL ANALYSIS

For our *final analysis* dataset, a nonuniform occupancy distribution was created for the $\{CM_1, CM_2\}$ map, instead of constant $\tau = 10$ (Fig. S13). As noted earlier, in thermal equilibrium, an occupancy map can be transformed into a corresponding free-energy landscape via the Boltzmann factor[12]: $\Delta G/k_B T = -\ln(n/n_0)$, where $n$ is the occupancy of the current state and $n_0$ is the occupancy of the maximum occupancy state in the state space. The lowest allowable occupancy for our map was chosen by taking the analysis in section SM-V in consideration.

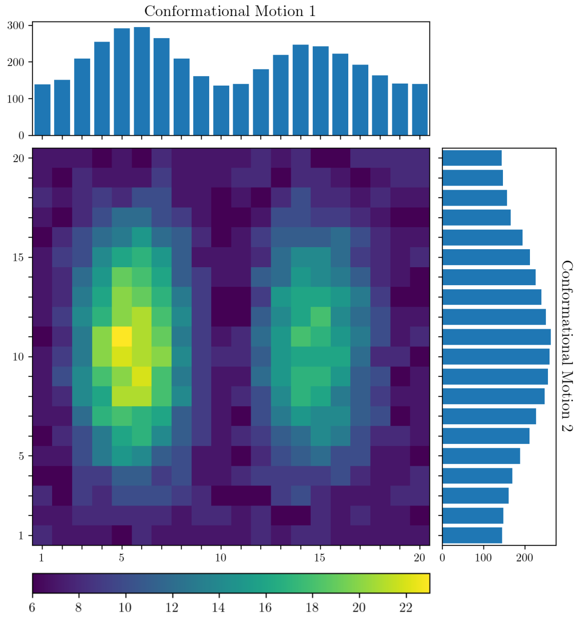


FIG. S13: 2D distribution of occupancies for all 400 states in $SS_2$, assigned equally for each PD. The net occupancy of this state space is 4000, such that the total number of images in the complete dataset is the product of 4000 and the number of PDs. The characteristics of this occupancy map were chosen to provide easily distinguishable features along both 1D and 2D conformational motions: specifically, bimodal and unimodal distributions for $CM_1$ and $CM_2$, respectively.

### SM-VIII. ANALYSIS OF 2D AND 3D MOVIES FOR FINAL ANALYSIS

The following two movies M3 and M4 show a selection of ESPER 2D and 3D sequences obtained from our final analysis.

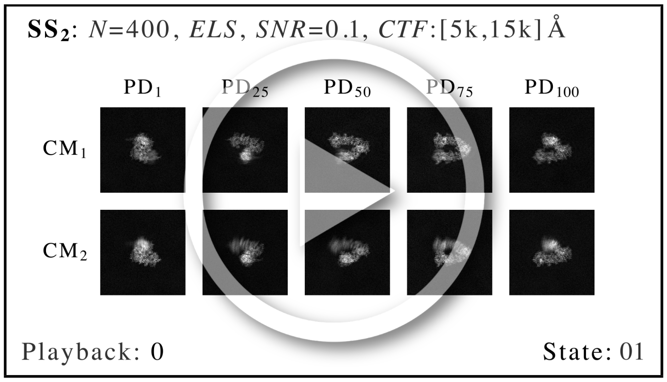


MOV. M3: Set of five 2D movies from $SS_2$ PD-manifold subspaces equispaced on a great circle. Each manifold is generated according to data-type III, with the exception that images are sampled from a nonuniform 2D occupancy map (Fig. S13). The set of CTF-corrected and Wiener-filtered snapshots within each bin are integrated, as opposed to the raw images themselves as previously shown in movie M2. Again, there is a significant difference in resolution between the desired CM and orthogonal CM, with the resolution of the desired CM superb across all states. See movie M4 for 3D movies from the full set of these 2D movies on $S^2$, where pairwise information from both CMs is incorporated to fully resolve all molecular domains in each volume. `https://www.dropbox.com/s/351288brw25si85/M3_2D_Movies_DT3_ELS.mp4?dl=0`

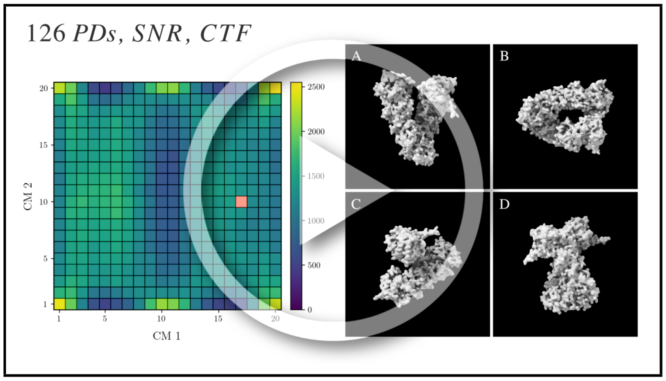


MOV. M4: Output volumes from our final-analysis dataset: 126 PDs, SNR = 0.1 and CTF with microscopy parameters as previously described. A sequence of 69 3D density maps is shown as seen from four orthographic views [A-D] animated along a chosen trajectory in the retrieved 2D state space. Here the 2D occupancy map before $R^2$-thresholding has been supplied (in contrast to Fig. 10), with all volumes reconstructed using RELION[13], without removal of any images in the original ensemble. Post-processing steps for display of each of these volumes included removal of dust (via Chimera's `hideDust` command with size 10) and application of a Gaussian filter (via Chimera, using 1 Å standard deviations of the 3D isotropic Gaussian function). https://www.dropbox.com/s/kfsddid347zb0j9/M4_3D_Movies_DT3_ELS.mp4?dl=0

### SM-IX. ANALYSIS OF SUBSPACE FITTING FOR FINAL ANALYSIS

In the following figure, we provide a collection of 2D subspaces for two PDs (left and right) for the final-analysis data type (i.e., data-type III with nonuniform occupancy assignments). As denoted in the blue boxes, the parabolic CM subspaces tend to curl inwards substantially near their boundaries. This inward-curling effect varies depending on the CM subspace and thus on the type of motion as visualized from the corresponding PD. For all PDs explored, our use of the general conic least-squares fit proved highly robust to these changes.

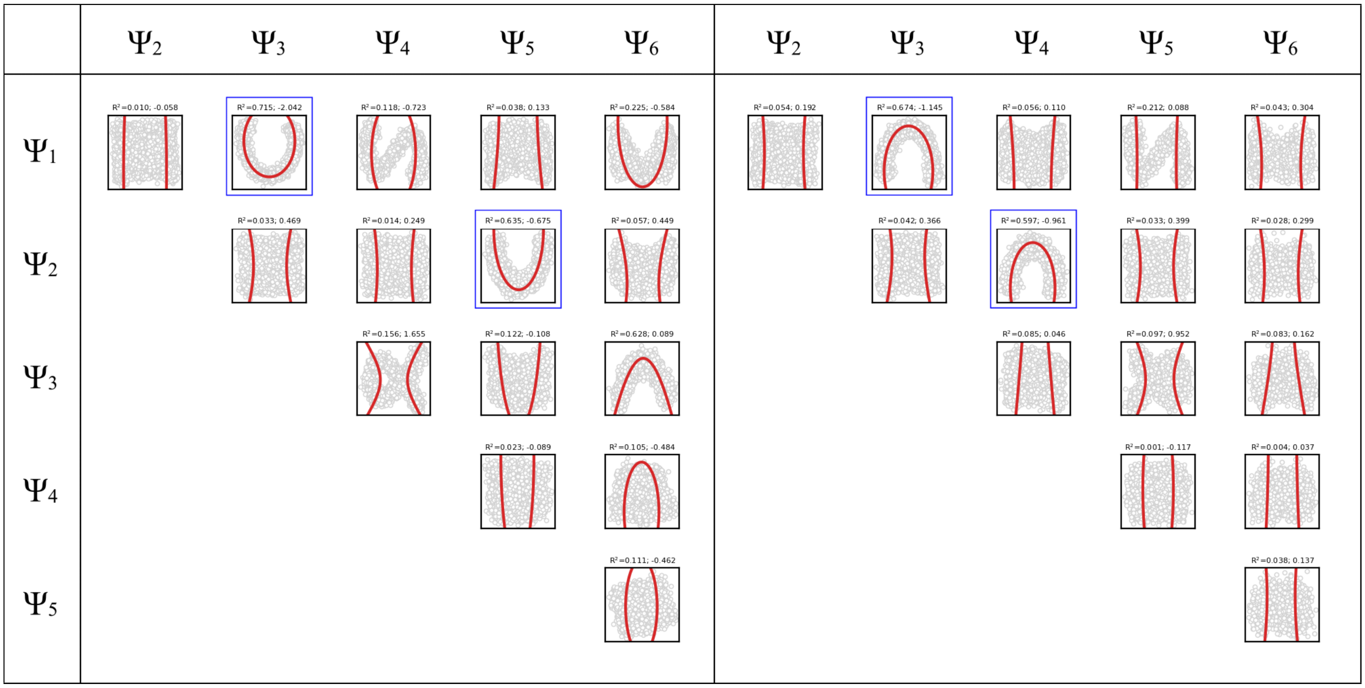


FIG. S14: Here we show the (1) coefficient of determination ($R^2$); and (2) discriminant of the implicit conic equation above each subplot. The $R^2$-value obtained for each 2D subspace is initially used before $d$-dim rotations to establish the location of parabolic modes. As can be seen above, this score is significantly higher for these parabola-housing subspaces. Across all 126 PDs, non-harmonic parabola-housing subspaces were predominantly defined by a negative (elliptic) discriminant, while harmonics were often associated with a positive (hyperbolic) value. A comparison of these plots with data-type II can be found in movie M7.

### SM-X. OVERVIEW OF DIFFUSION MAPS

Given a set of $N$ images, the diffusion map (DM) approach seeks to generate an optimal embedding of the data in a low-dimensional space, so as to preserve all relevant information. Below, we outline the DM framework with considerations taken for the synthetic data explored in our main text. As a preliminary step, we normalize each of the images by removing the mean and scaling to unit variance. As the kernel required for the DM framework is formed using pairwise distances, we first create the distance matrix $\boldsymbol{D}$ (for each PD independently) by calculating the Euclidean distance for every pairwise combination of its $N$ images. The Euclidean distance between two images $X = (x_1, x_2, \ldots, x_P)$ and $Y = (y_1, y_2, \ldots, y_P)$, where $x_i$ and $y_i$ denote the intensities at pixel $i$ in images $X$ and $Y$ (each having $P$ pixels), is defined[14] by

$$D_{X,Y} = \left( \sum_{i=1}^{P} (x_i - y_i))^2 \right)^{1/2}$$

The pairwise distances form a symmetric $N \times N$ square matrix, where a single row represents the distance of the corresponding row-indexed image to each column-indexed image. Next, an isotropic Gaussian kernel is applied to these distances to create a real, symmetric similarity matrix

$$A_{ij} = \exp\left( -\frac{D_{ij}^2}{2\varepsilon} \right)$$

The similarity matrix $\boldsymbol{A}$, calculated using a suitable $\varepsilon$ value, is then divided by a diagonal matrix of its row sums to construct a symmetric, positive semidefinite stochastic Markov transition matrix $\boldsymbol{M}$, representing the relative pairwise affinity between all images[15]. Eigendecomposition of the matrix $\boldsymbol{M}$ is then performed to retrieve an ordered set of $N$ eigenvalues and corresponding eigenvectors (leaving out the steady-state $\{\lambda_0, \Psi_0\}$ eigenvalue-eigenvector pair, where $\lambda_0 = 1$ due to the graph being fully connected[16]), which define a nonlinear spectral embedding of the data[10].

The Gaussian bandwidth ($\varepsilon$) in the above expression has a strong influence on the definition of similarity between our simulated images. At small Gaussian bandwidths, the system takes on a relatively fine-grained definition of similarity (i.e., data points only see their direct neighbors). Increasing $\varepsilon$ transforms this relationship into a more coarse-grained notion of similarity. These notions of similarity govern the behavior of all subsequent steps, and ultimately impact the geometric structure of the resultant manifold embedding, and thus the DM eigenfunctions. Particularly, in the limit $\varepsilon \mapsto 0$ and $N \mapsto \infty$, and with an appropriate normalization of the similarity matrix[16], the diffusion map eigenvectors converge to the eigenfunctions of the LBO[10].

### SM-XI. ESTIMATION OF GAUSSIAN BANDWIDTH AND INTRINSIC DIMENSIONALITY

A detailed analysis of the preferred Gaussian bandwidth regime for both pristine and noisy PD datasets is available in section SM-XIV-D. In summary, for all PD datasets explored, the choice of appropriate Gaussian bandwidth $\varepsilon \in [a,\, b]$ proved highly flexible, presenting embeddings with virtually identical cosine eigenfunctions across several orders of magnitude. For values slightly below this range, suboptimal yet structured properties emerged. Even farther below this range, the corresponding eigendecompositions did not converge, such that the structures of these embeddings became jumbled in nonsensical patterns. Similar disruptions occurred for all embeddings generated with $\varepsilon > b$, with such anomalies likely arising due to arithmetic underflow encountered during computation of the Gaussian kernel. For noisy datasets, the range $\varepsilon \in [a,\, b]$ coincided with the prescribed range of $\varepsilon$ values defined via a prominent routine for automating this decision (Fig. S15), which uses the correlation dimension as a measure of fractal dimensionality[10,15,17] (henceforth referred to as the bandwidth estimation method). We have provided an automation strategy[18] in our online repository[19], which uses the inflection point of a fitted hyperbolic tangent to select this value. This automated value was often just shy of retrieving optimal results on pristine datasets, while excelling for cases involving experimentally relevant SNR and $\tau$.

While proponents of the bandwidth estimation method also claim to predict the intrinsic dimensionality $n$ of the dataset[15], we found this recipe to be inconsistent with ground truth (as detailed in Fig. S15 and Fig. S16). The following two figures show the results of using the bandwidth estimation plots as procured following the fractal dimensionality method[10,15,17], from which it is expected that (1) the linear region of each plot delimits the range of optimal $\varepsilon$ values, and (2) twice its slope defines the intrinsic dimensionality of the dataset. While this method's first prong proved accurate in discovering a suitable $\varepsilon$ value for all datasets observed, the second prong always proved highly inaccurate for PD datasets. For example, in Fig. S15, the slopes of each PD within each SS grouping ($SS_2$ grouped at top and $SS_1$ at bottom) differ significantly (e.g., within $SS_1$, the slopes for $PD_1$ through $PD_5$ signified a dimensionality range of 1.55 to 1.95). As the ground-truth dimensionality of $SS_1$ defined via the rotation of a domain of an atomic structure is one, these trends demonstrate that the bandwidth estimation method—while sufficient for measuring the proper $\varepsilon$ value—is insufficient for correctly determining the dataset's intrinsic dimensionality. Fig. S16 provides further investigation of these results in the presence of noise, where the intrinsic dimensionality was still significantly overestimated.

Given these results, there remains uncertainty on how to best determine the intrinsic dimensionality of any given dataset (i.e., the number of CMs present to search for). The estimation of the intrinsic dimensionality of a manifold is a longstanding mathematical problem[20,21], which we consider an open issue. As it stands, this information must instead be intuited via a careful analysis of final outputs (i.e., 2D and 3D movies) generated from our framework.

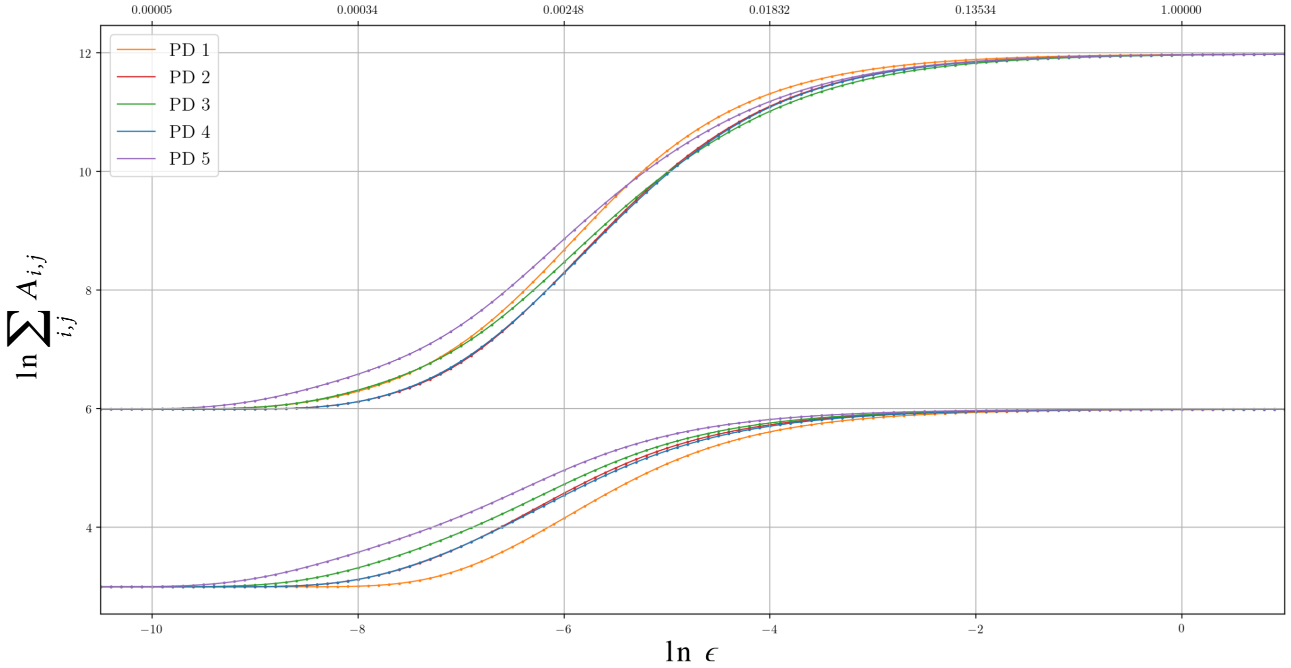


FIG. S15: bandwidth estimation plots as obtained via the fractal dimensionality method. For all PDs, the optimal $\varepsilon$ values were manually checked through trial and error, with the optimal range of $\varepsilon$ values approximately $[10^{-4}, 10]$. It can be seen that the linear portions of all bandwidth plots for each PD cover a similar $\varepsilon$ range, regardless of $SS_1$ or $SS_2$ (bottom group and top group, respectively). Instead, PDs from either state space differ mainly by their vertical position in the plot.

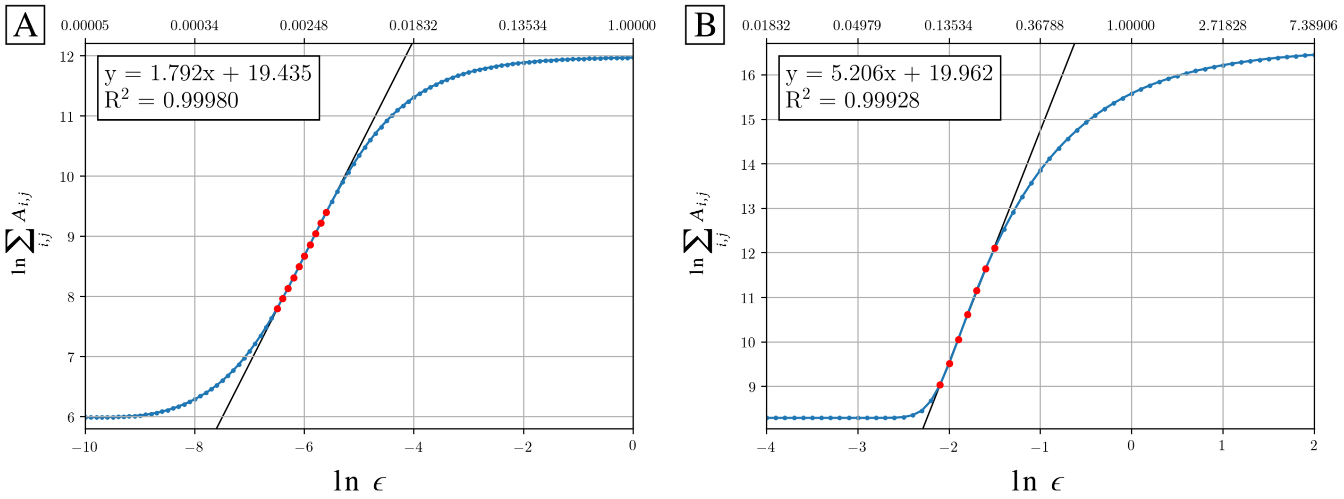


FIG. S16: Bandwidth estimation plots are shown for [A] $PD_2$ in $SS_2$ ($SNR_\infty$, as previously shown in Fig. S15) and [B] $PD_2$ in $SS_2$ ($SNR = 0.1$ and $\tau = 10$). As can be seen via twice the slope of the fit for each linear region, the ground-truth intrinsic dimensionality of these datasets ($n = 2$) is erroneously defined via this method, and increasingly so with the introduction of noise. However, even in the presence of noise, the criterion for the optimal $\varepsilon$ range still proved suitable, with the corresponding eigendecomposition converging for $\varepsilon \in [10^{-11}, 1]$. Note also the asymmetry of each curve and the magnitude of their curvature, which leads to uncertainties in fitting these regions with the prescribed linear model.

### SM-XII. COMPARISON OF PCA AND DM

In this section, we complete our comparison of diffusion maps with principal component analysis. First, we briefly readdress our *a priori* expectations as informed by spectral theory[22]. Both DM and PCA are kernel methods which entail the use of a symmetric matrix (i.e., a Markov transition matrix and a covariance matrix, respectively). Symmetric matrices have many convenient properties, and, particularly for our interest, are diagonalizable with mutually orthogonal eigenvectors, and corresponding real, non-negative eigenvalues. As shown via the principal axes theorem, this diagonalization is determined by the matrix's eigenvectors, which are used to align the innate principal axes of the graph into standard position. When organized along these principal axes, distinct classes of geometries associated with the quadratic form of the symmetric matrix (called "quadric hypersurfaces") clearly emerge, with the characteristics of each surface specifically defined by its corresponding eigenvalue[22].

When a symmetric matrix has only positive or null eigenvalues, the matrix is "positive semidefinite". Low-dimensional quadrics generated by positive semidefinite matrices include elliptic cylinders, parabolic cylinders, hyperbolic cylinders and cones[23]). However, our case is of substantial complexity, as we will be investigating quadric hypersurfaces of graphs generated with inclusion of multiple degrees of freedom and noise. Indeed, as the symmetric matrix of both PCA and DM is positive semidefinite, we anticipate to recognize one of these forms within subspaces of each subsequent embedding (e.g., parabolic cylinders in Fig. S6). With these fundamental similarities in mind, we next display figures (Fig. S17-18) referenced in our main text, which show striking similarity between results obtained by these two techniques. This is further supported by the fact that the eigenvectors of DM generated with very large $\varepsilon$-values converge to the eigenvectors of PCA.

### SM-XIII. THE LAPLACE-BELTRAMI OPERATOR ON A RIEMANNIAN MANIFOLD

The Laplace-Beltrami operator (LBO) acting on a scalar function $f$ on a compact (closed and bounded)[24] Riemannian manifold is given by[25–27]

$$\nabla^2 f = g^{-1/2}\ \partial_i\ (g^{1/2} g^{ij}\ \partial_j\ f)$$

where $g = \det(g^{ij})$ and $g^{ij}$ are the components of the metric tensor. Specifically, we are interested in the eigenfunctions of the LBO, $\nabla^2 f = \lambda f$, noting these form a complete basis in the functional space $L_2(\Omega)$ of measurable and square-integrable functions on the manifold $\Omega$[28]. For a bounded manifold, the eigenfunctions must further satisfy boundary conditions; for example, DM requires the Neumann boundary conditions[16], such that the derivatives on the boundaries vanish. Therefore, the eigenfunctions depend also on the boundary of $\Omega$.

If $g$ is constant on $\Omega$, the expression simplifies to $\nabla^2 f = g^{ij}\ \partial_i\ \partial_j\ f$, where $g^{ij}$ are constant values. For example, in two dimensions

$$\nabla^2 f = (g^{11}(\partial_1)^2 + g^{22}(\partial_2)^2 + 2g^{12}\partial_1\partial_2)f$$

Moreover, if $g^{11} = g^{22} = 1$, we recognize the common Laplacian in Euclidean coordinates, which is the sum of pure second derivatives[25]

$$\nabla^2 = \sum_{i=1}^{d} \partial_i^2$$

It is well understood that the eigenfunctions of the LBO on a manifold $\Omega$ carry useful information about its intrinsic geometry, and are thus important for understanding many systems. For compact manifolds with a boundary, as an example, these eigenfunctions are the modes of vibration of a string (1D) or a membrane (2D). For compact manifolds without a boundary (closed manifolds), the well-known spherical harmonics are eigenfunctions on the surface of the 2-sphere. In the field of structural biology, the eigenfunctions of the LBO on SO(3), which are the Wigner-D functions, have been used for retrieving the unknown orientations of single-particle X-ray and cryo-EM snapshots[29]. In general, the eigenfunctions of the LBO on different manifolds are fundamental to mathematics and science, and describe a wide diversity of seemingly disparate phenomena—reflecting the so-called "underlying unity of nature"—from quantum mechanics to gravitational fields[30].

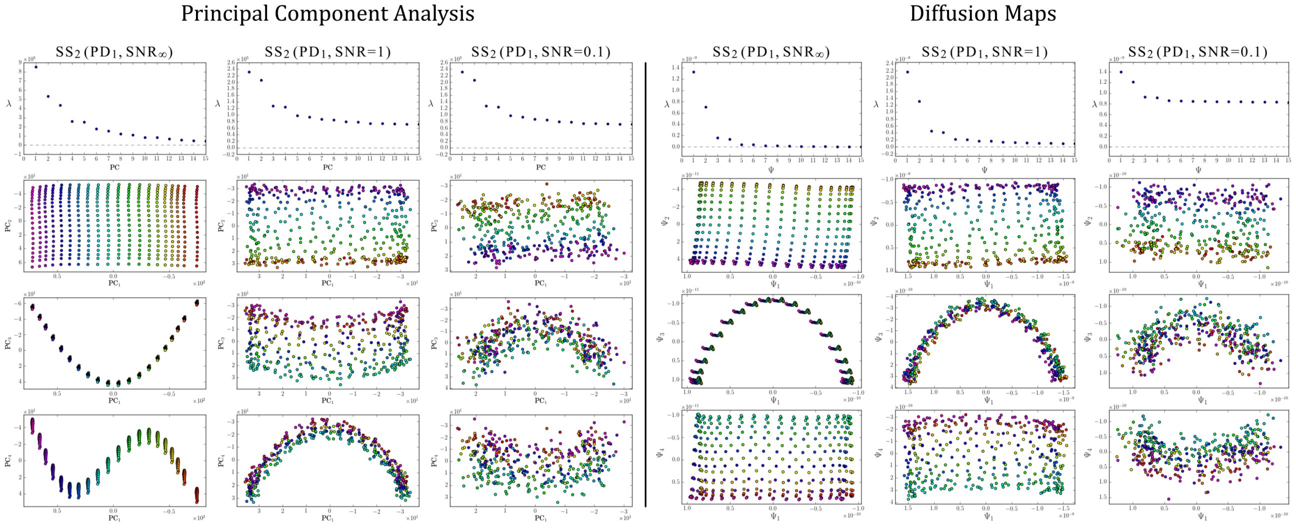


FIG. S17: Comparisons of 2D subspaces and eigenvalue spectra obtained via PCA (left) and DM (right) for $PD_1$ in $SS_2$ across three SNR regimes (one SNR regime per column; with uniform occupancy $\tau = 1$). As can be seen in both linear and nonlinear dimensionality reduction frameworks, the well-defined structure of these subspaces deteriorates rapidly as increasing amounts of additive Gaussian noise is introduced on each image. Overall, the outputs of PCA on these datasets revealed a striking resemblance to those produced by DM. Importantly, the parabolic mode is conserved for both frameworks even within experimental regimes (SNR $= 0.1$).

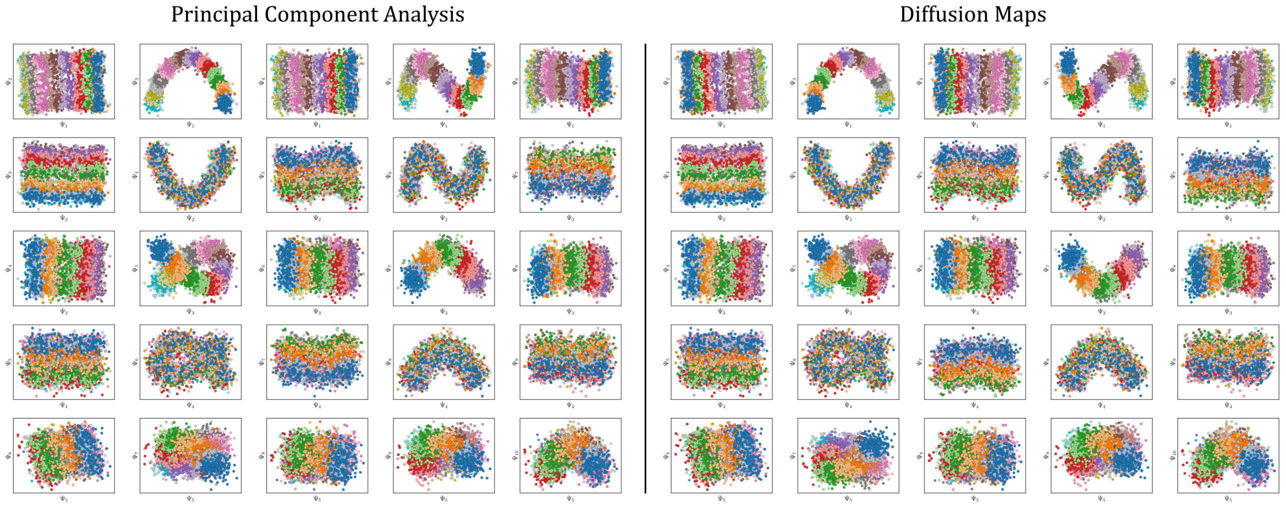


FIG. S18: Comparison of 2D subspaces obtained via PCA (left) and DM (right) for $PD_2$ in $SS_2$, with image sets generated with SNR $= 0.1$ and $\tau = 10$. Colors have been assigned to data points so as to match the ground-truth indices of states covering $CM_2$, which is the most visually pronounced motion as viewed from this PD. Here, the $CM_2$ subspaces can be seen in the first row ($\{\Psi_1, \Psi_i \mid i > 1\}$), the $CM_1$ subspaces in the second row ($\{\Psi_2, \Psi_j \mid j > 2\}$), $CM_2$ first-harmonics in the third row (($\{\Psi_3, \Psi_k \mid k > 3\}$)), $CM_1$ first-harmonics in the fourth row, and $CM_2$ second-harmonics in the fifth row. Overall, the similarity in outputs between these two frameworks is undeniable, with the only visual difference appearing in the arbitrary directionality (*sense*) of each coordinate; which is to be expected.

### SM-XIV. ANALYSIS OF THE LBO EIGENFUNCTIONS

In the following subsections (A-D), we perform an analysis of the eigenfunctions of the LBO on a set of distinct manifolds $\Omega$. First, in subsection A, we use the DM framework to investigate the known eigenfunctions of the LBO on the interval and rectangular domains, which are cosines. We expect this investigation to inform our heuristic discoveries for the PD embeddings ($\Omega_{\mathrm{PD}}$), where similar cosine eigenfunctions were observed for each degree of freedom. As well, we will use these ideal manifolds in A to build intuition for rotations needed to realign essential eigenfunctions, which were observed in the $\Omega_{\mathrm{PD}}$ embeddings. Following this analysis, we ultimately detail how the structure of manifolds obtained from a conformational state space transforms as the data type is translated stepwise from (B) atomic models $\Omega_{\mathrm{ACS}}$, to (C) 3D electron density maps $\Omega_{\mathrm{EDM}}$, and finally, to (D) 2D projections $\Omega_{\mathrm{PD}}$. In the last case, recall that 2D projections are the only form of data readily accessible in a cryo-EM experiment.

#### *A. Eigenfunctions of the latent space*

To get insight into the characteristics of the $\Omega_{\mathrm{PD}}$ eigenfunctions, we abstract the manifold of the PD-dataset as a Euclidean space with rectangular boundaries. This is motivated by the most simple representation of our ground-truth state space of atomic models, where the relationship between equispaced coordinates in the prior matches the relationship between equiangular molecular domain rotations in the latter. By separately embedding the collection of states in each of these two data types and comparing their resulting eigenfunctions, we will show that these two spaces are nearly identical. In effect, the rectangular domain can be viewed as the conformational *latent space* to which our collection of more advanced state spaces is compared. We will additionally show that for the embeddings of 3D electron density maps and 2D projections, the mapping relative to the latent space becomes distorted. This effect can be explained by a change of the metric induced in the process.

In the 1D space, a set of pairwise distances between a collection of equispaced coordinates on a line carries all essential information necessary to model the pairwise distances between a sequence of atomic models with molecular domain rotated by a constant angular increment. To represent our $\mathrm{SS}_1$ PD dataset, we uniformly sample $N = 50$ equispaced points from a 1D interval $X \in [0, \ell = 1] \subset \mathbb{R}$, with each of these 50 points representing a unique state of the molecule. Following the DM framework, we then calculate the distance matrix for this collection of points and embed the data in a low-dimensional space. As a note, for all embeddings that follow, we will show that two characteristic regimes emerge depending on choice of Gaussian bandwidth, which we will denote with $\varepsilon_{\downarrow}$ and $\varepsilon_{\uparrow}$ for the small and large regime, respectively. For the smaller Gaussian bandwidth, a cosine series emerged for all eigenfunctions (Fig. S19-A), in very good agreement with the Laplacian on a 1D Euclidean interval with Neumann boundary conditions. Specifically, we anticipate (and retrieve) canonical[28] eigenfunctions of the form $\psi_v(x) = \{\cos(v\pi x/\ell) \mid k \geq 1\}$. As the Gaussian bandwidth was incrementally increased from $\varepsilon_{\downarrow}$ to $\varepsilon_{\uparrow}$, this cosine series smoothly transformed into a different complete, orthogonal set: the Legendre polynomials (Fig. S19-B). These Legendre polynomials, however, only occur for hyperrectangles, which are $n$-dimensional Cartesian products of orthogonal intervals

Next, to represent our $\mathrm{SS}_2$ PD dataset, we uniformly sample $N = 50 \times 50$ points from a 2D interval $X \times Y \in [0, \ell_x = 1.1] \times [0, \ell_y = 1] \subset \mathbb{R}^2$. For its illustrative properties, we avoid degeneracy by ensuring $(\ell_x/\ell_y)^2$ is not a simple ratio[28]. Again, we follow the DM framework by calculating the pairwise distances between these points and embedding the data in a low-dimensional space. As demonstrated in Fig. S19-C, the set of eigenfunctions obtained in the smaller Gaussian bandwidth regime matched our *a priori* expectations for the Laplacian on a rectangular domain with Neumann boundary conditions. These canonical eigenfunctions are[28]

$$\psi_{vw}(x, y) = \{\cos(v\pi x/\ell_x)\cos(w\pi y/\ell_y) \mid v, w \leq 0\}$$

following the same pattern for higher-dimensional domains $\Omega_R = [0, \ell_1] \times \cdots \times [0, \ell_n] \subset \mathbb{R}^n$ (with $\ell_i > 0$). Again, as we incrementally increased the Gaussian bandwidth from $\varepsilon_{\downarrow}$ to $\varepsilon_{\uparrow}$, this set of complete and orthogonal cosines smoothly transformed into the orthogonal Legendre polynomials set, which are now a function of both $x$ and $y$, as expected (Fig. S19-D). Importantly, the leading Legendre polynomials provide a direct, linear map of the input data points, a consequence of the linear terms $\mathrm{P}_1(x)$ and $\mathrm{P}_1(y)$. This is preferred over the non-linear map achieved by the cosine functions, as it is unencumbered by nuisances such as non-uniform rates of change and parabolic harmonics. In subsections C and D, however, we will show that this preferred behavior cannot be obtained for 3D electron density maps and 2D projections.

Returning to the smaller of the two Gaussian bandwidth regimes, we next compare the previous non-degenerate rectangular results to those from a degenerate square domain, with $N = 50 \times 50$ points equispaced identically along $X$ and $Y$ (Fig. S20-A). Due to the presence of degenerate eigenvalues, which can arise for domains with a rational ratio[28] $(\ell_x/\ell_y)^2$, we encounter pairs of eigenfunctions that appear different from the non-degenerate case of the rectangle[28] (as seen, for example, by pairs $\{\Psi_1, \Psi_2\}$ and $\{\Psi_4, \Psi_5\}$ in Fig. S20-A). In Fig. S20-C, we illustrate that these eigenfunctions are just rotated within their degenerate space, exactly as expected. We note that an eigenfunction associated with a degenerate eigenvalue is a linear combination of the degenerate eigenfunctions[28], where the normalization of the eigenfunctions restricts this linear transformation to a rotation and reflection (i.e., the group of orthogonal

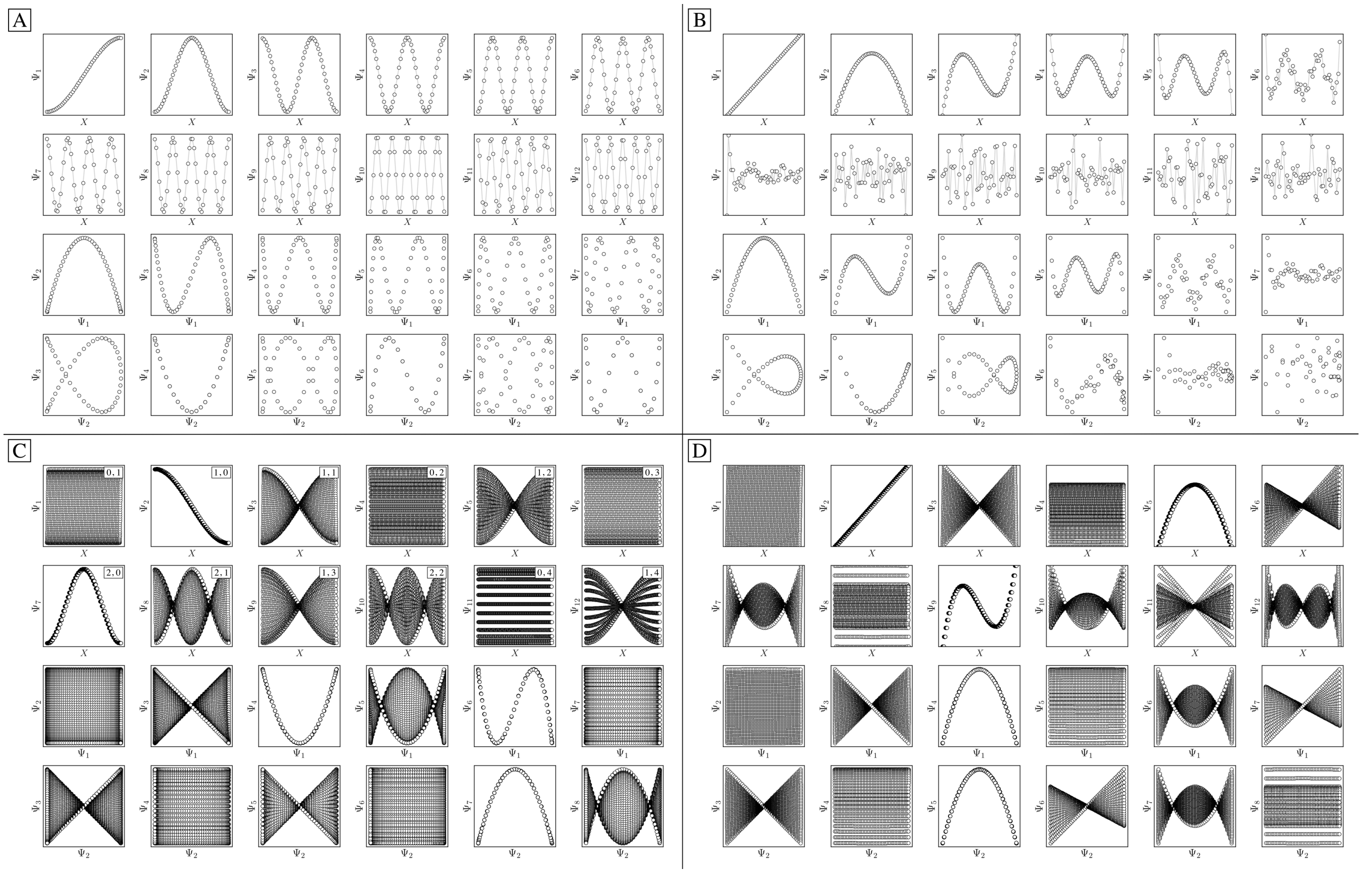

FIG. S19: DM eigenfunctions of the 1D interval for small ($\varepsilon_{\downarrow} = 5 \times 10^{-5}$) and large ($\varepsilon_{\uparrow} = 10$) Gaussian bandwidths are shown in [A] and [B], respectively. Likewise, eigenfunctions of the $N = 50 \times 50$ rectangular (nondegenerate) domain for small and large Gaussian bandwidth are shown in [C] and [D], respectively. As was done in the main text, eigenfunctions have been independently displayed by indexing each by its ground-truth ordering via sequential $x$-coordinates. For [C] and [D], a similar appearance of eigenfunction plots, albeit interchanged, would be seen when indexing instead via sequential $y$-coordinates. In [C], each eigenfunction's corresponding modes $\{v, w\}$ have also been provided in the top right-hand corner. For all four subplots, pairwise combinations of eigenfunctions are additionally shown, which can be visualized after an embedding without any ground-truth knowledge.

transformations). For example, the $\{\Psi_1, \Psi_2\}$ pair is of form $\Psi' = \mathbf{R}^T \Psi$ such that

$$\begin{bmatrix}\Psi'_1 \\ \Psi'_2\end{bmatrix} = \begin{bmatrix}\cos(\theta) & \sin(\theta) \\ -\sin(\theta) & \cos(\theta)\end{bmatrix} = \begin{bmatrix}\cos(\pi x) \\ \cos(\pi y)\end{bmatrix}$$

$$\begin{bmatrix}\Psi'_1(\theta) \\ \Psi'_2(\theta)\end{bmatrix} = \begin{bmatrix}\cos(\theta)\cos(\pi x) + \sin(\theta)\cos(\pi y) \\ -\sin(\theta)\cos(\pi y) + \cos(\theta)\cos(\pi x)\end{bmatrix}$$

As seen for $\Psi_6$ in Fig. S20-A, these summands can also have the form of two products $\Psi_6 = A\cos(\pi x)\cos(2\pi y) + B\cos(2\pi x)\cos(\pi y)$, with any $A$ and $B$ such that $A^2 + B^2 \neq 0$. Hence, it can be seen that these aberrant eigenfunction pairs are defined by an admixture of cosines in a higher-dimensional space, with form

$$\begin{aligned}\Psi_i &= A\cos(v\pi x)\cos(w\pi y) + B\cos(w\pi x)\cos(v\pi y) \\ &= A\psi_{vw} + B\psi_{wv}\end{aligned}$$

By using an appropriate rotation operator $R_{i,j}$, the summands within each eigenfunction pair can be maximally separated among both members $\Psi_i = \psi_{vw}$ and $\Psi_j = \psi_{wv}$, such that the canonical eigenbasis is recovered (Fig. S20-B). As demonstrated using analytical expressions $\psi_{1,0} = \cos(\pi x)$ and $\psi_{0,1} = \cos(\pi y)$ in Fig. S20-C, this separation occurs multiples of $\theta = 90°$ apart. In the Fig. S20-C example, at $R_{1,2}(45°)$, these eigenfunctions have form

$$\begin{bmatrix}\Psi'_1(\theta = 45°) \\ \Psi'_2(\theta = 45°)\end{bmatrix} = \begin{bmatrix}\sqrt{2}/2\cos(\pi x) + \sqrt{2}/2\cos(\pi y) \\ -\sqrt{2}/2\cos(\pi y) + \sqrt{2}/2\cos(\pi x)\end{bmatrix}$$

which decouples back into two distinct modes ($\cos(\pi y)$ and $\cos(\pi x)$ for $\Psi_1$ and $\Psi_2$ respectively) at $R_{1,2}(90°)$. A similar result is obtained by applying this operation on the appropriate eigenfunctions obtained via DM, with each initially assuming a random rotation angle (Fig. S20-A) requiring a specific correction $R_{i,j}(\theta)$ (Fig. S20-B).

While degeneracy suggests a possible cause for the appearance of eigenfunction misalignments in $\Omega_{\mathrm{PD}}$, we note that it is a rather rare event in our data sets and that it

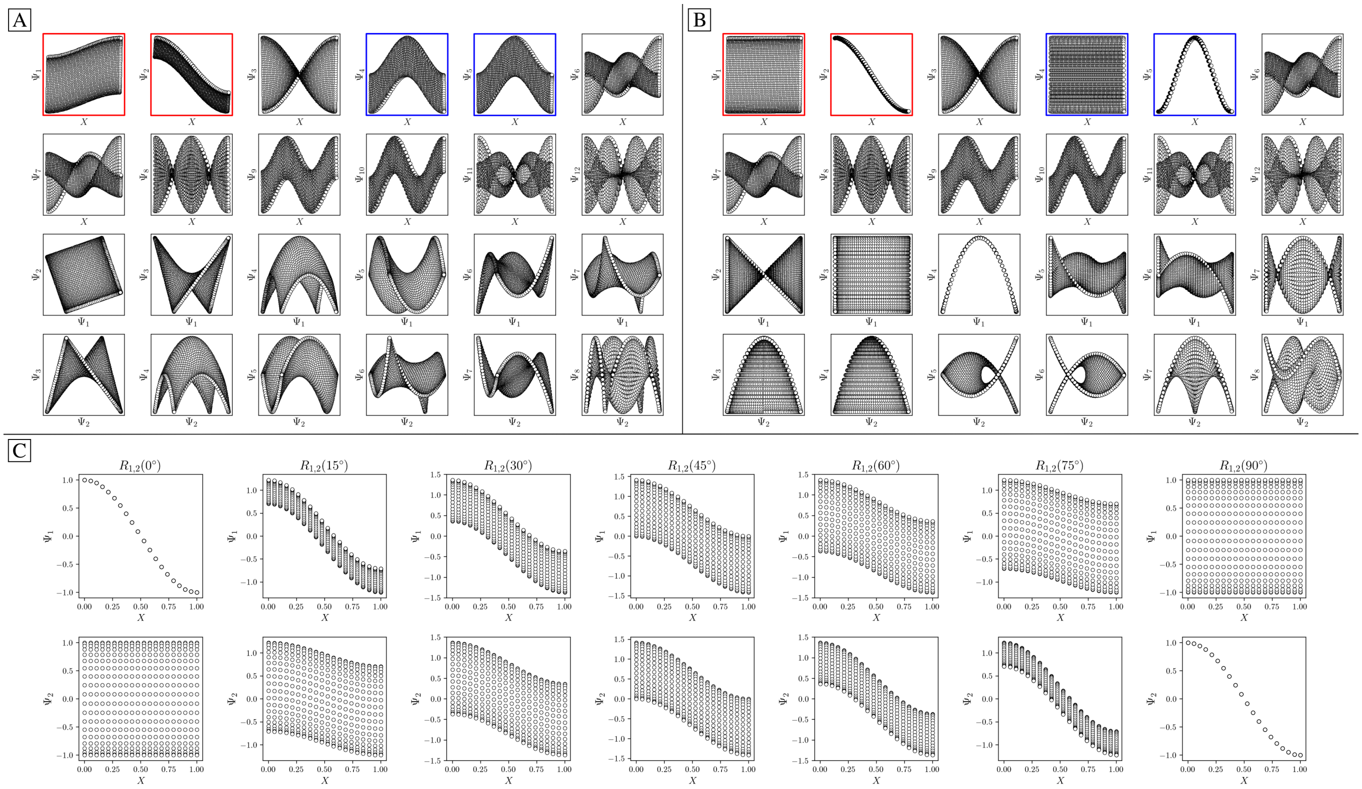


FIG. S20: DM eigenfunctions of the $N = 50 \times 50$ square domain for small Gaussian bandwidth ($\varepsilon_{\downarrow} = 5 \times 10^{-5}$) are shown in [A] and [B] before and after high-dimensional rotations, respectively. It can be seen here that pairs of eigenfunctions exist that contain relationships aberrant to the canonical eigenfunction form (Fig. S19-C). Two such pairs have been highlighted in red and blue, respectively, with the members of each pair always rotated 90° apart. To note, as any rotation can happen in the presence of degeneracy, this initial rotation is an arbitrary one. This property is demonstrated via the schematic in [C], which shows the angular relationship between two analytically-generated functions ($\cos(\pi x)$ and $\cos(\pi y)$, each displayed in the reference frame of $X$) as they are jointly rotated 90°. By applying rotation operators $R_{1,2}(\theta) = -19°$ and $R_{4,5}(\theta) = 45°$ independently to two such aberrant pairs in [A], the canonical eigenfunction form begins to recover in [B], and more so as additional operators are intelligently applied.

can be identified from the eigenvalue spectrum. Pairs of rotated eigenfunctions, at least approximately, can also be mimicked when domains have undergone certain elementary geometric transformations. For example, by performing an affine transformation on a rectangle $\Omega_R$ to form a parallelogram $\Omega_P$ , we observed a rotation of the first two eigenvectors, as similarly seen in Fig. S20-A. Recall that an affine mapping preserves collinearity and ratios of distances, but in general not distances and angles. In subsection C, we will explore the possibility of other classes of geometric transformations on $\Omega_R$.

As a final point in this section, we illustrate our method for retrieving the canonical eigenfunctions buried within an embedding, which has been used in Fig. S19 and Fig. S20, and extensively throughout the main text. Fig. S21 provides a schematic using the known analytical eigenfunctions (Fig. S21-B and Fig. S21-C) chosen so as to match the results from DM on the square, degenerate $\Omega_R$. In Fig. S21-A, we display the eigenfunctions from Fig. S20-A in two different reference frames corresponding to our ground-truth knowledge. Specifically, we plot the points in each eigenvector in a sequence corresponding to their initial ground-truth arrangement along each degree of freedom (for the rectangular Euclidean space, along either $X$ or $Y$), which is shown in the first and second row of Fig. S21-A, respectively. As shown in Fig. S21-B and Fig. S21-C, a given reference frame captures the eigenfunction on a projected plane in the $n$-dimensional space where it resides.

#### B. Eigenfunctions of the atomic models

We next investigate the manifolds obtained from the state spaces formed from a quasi-continuum of atomic-coordinate structures, each represented by a set of 3D atomic-coordinates $3m$ (e.g., Fig. S2). We generate these structures as described in the first step of our synthetic-generation protocol (section SM-I), which are subsequently used there to produce a corresponding set of 3D electron density maps and 2D projections. Importantly, the set of these 3D atomic-coordinate structures in

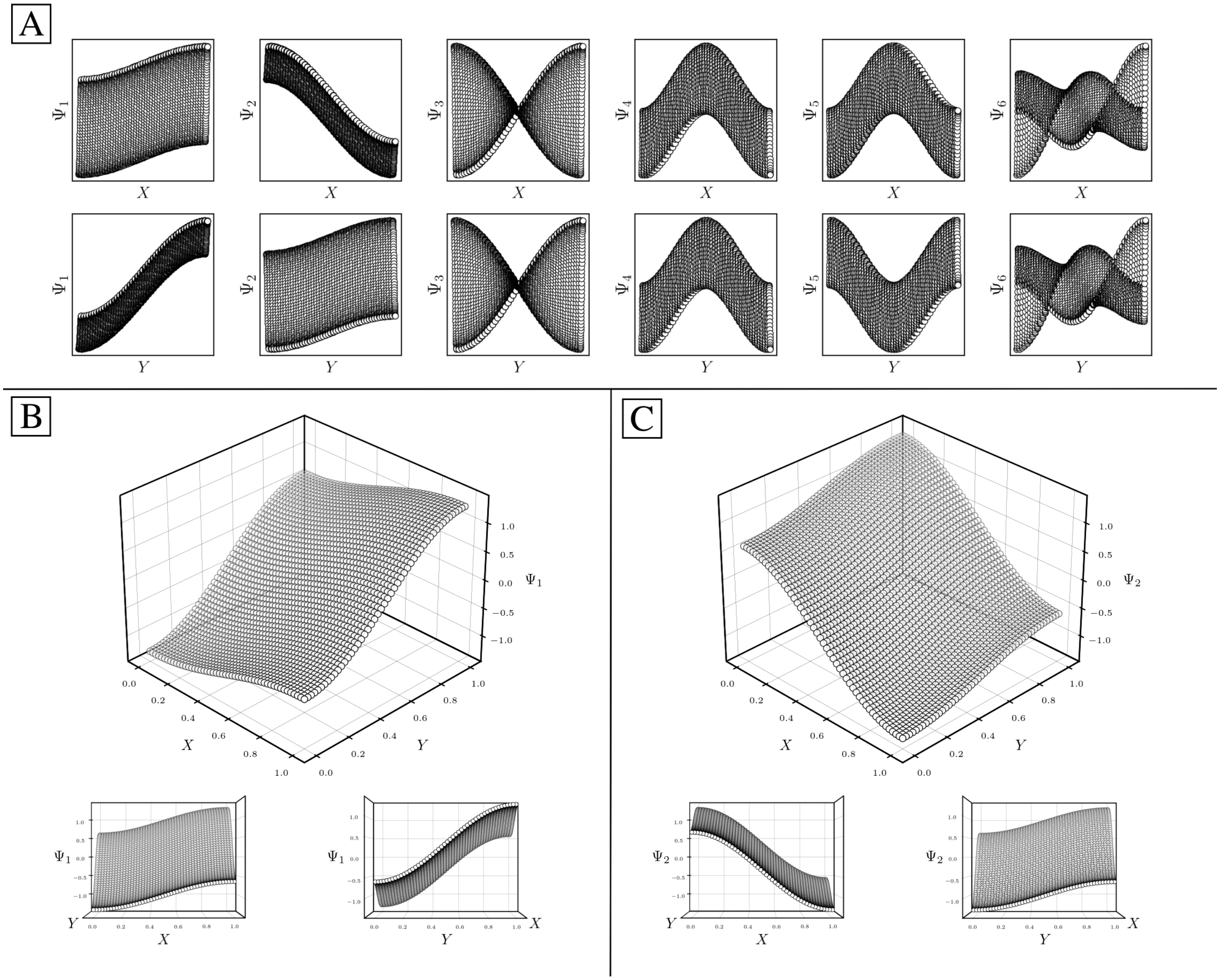


FIG. S21: DM eigenfunctions from Fig. S20-A are shown again in the first row of [A], which were displayed by ordering the points of the embedding in a sequence based on the ground-truth $x$-coordinates. The second row of [A] displays these eigenfunctions instead via the sequential ordering of ground-truth $y$-coordinates. Subplots [B] and [C] are analytically generated so as to match the appearance of $\Psi_1$ and $\Psi_2$ in the first and second row of [A], respectively. For this presentation, the equations $\Psi_1 = \cos(\theta)\cos(\pi x) + \sin(\theta)\cos(\pi y)$ and $\Psi_2 = -\sin(\theta)\cos(\pi x) + \cos(\theta)\cos(\pi y)$ were used, with $\theta = 250°$. As can be seen in [B] and [C], each eigenfunction exists in an $n$-dimensional space defined by the $n$ degrees of freedom of the system. Thus, by displaying a point from the manifold along a sequence in the embedding corresponding to a known degree of freedom, we are effectively viewing each eigenfunction on a projected plane in its $n$-dimensional space.

$\Omega_{\text{ACS}} \subset \mathbb{R}^{3m}$ represent the fundamental biophysical identity of each state, from which the cryo-EM experiment could only obtain two-dimensional information in the form of images. Following the DM framework, we first calculated the distance matrix for $\text{SS}_2$, which we obtained by the root-mean-square deviation (RMSD) for each pair of its 400 3D atomic-coordinates structures (PDB files). The RMSD between two sets of atomic-coordinate structures $X = (x_1, x_2, \ldots, x_m)$ and $Y = (y_1, y_2, \ldots, y_m)$ each composed of $m$ atoms is defined as

$$\text{RMSD}(X, Y) = \sqrt{\frac{1}{m}\sum_{i=1}^{m} \|(x_i - y_i)\|^2}$$

which is up to an irrelevant factor of $m^{-1/2}$ equal to the Euclidean distance $D(X, Y)$.

The resulting DM embeddings for the small and large Gaussian bandwidth regimes are shown in Fig. S22-A and Fig. S22-B, respectively, and share a strong resemblance to those found for the latent rectangular domain (Fig. S19). Again we note the presence of cosine eigen-

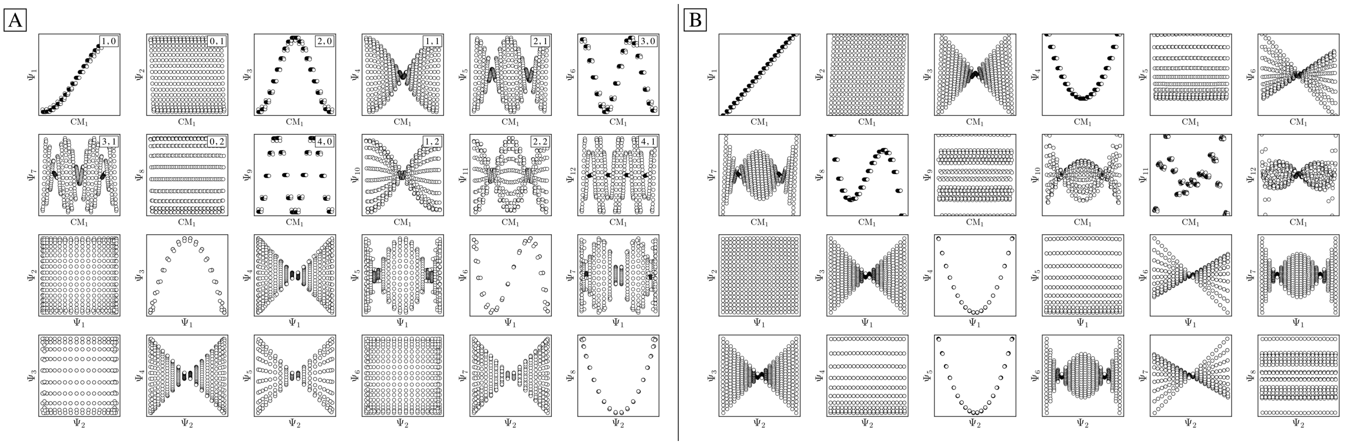


FIG. S22: DM eigenfunctions obtained for $20 \times 20 = 400$ atomic models occupying $SS_2$ for small ($\varepsilon_{\downarrow} = 0.1$) and large ($\varepsilon_{\uparrow} = 1000$) Gaussian bandwidths are shown in [A] and [B], respectively. Leading eigenfunctions are displayed in the first two rows via sequential indexing along the ground-truth $CM_1$ coordinates (i.e., equispaced rotations of chain A). In [A], each eigenfunction's corresponding modes $\{v, w\}$ are provided in the top right-hand corner, showing exceptional agreement with the LBO eigenfunctions on a rectangular domain with Neumann boundary conditions. We additionally note the absence of any significant eigenfunction misalignments.

functions for the small Gaussian bandwidth regime, and a near-perfect linear form (via leading Legendre polynomials) in the large Gaussian bandwidth regime ($\{\Psi_1, \Psi_2\}$ in Fig. S22-B). For the latter, we will show this feature is a luxury not obtained in the other data types explored. In the small Gaussian bandwidth regime, we can identify both our $CM_1$ and $CM_2$ parabolas residing in the subspaces $\{\Psi_1, \Psi_3\}$ and $\{\Psi_2, \Psi_8\}$, respectively. Similar results were found for the manifold embeddings generated using 3D atomic-coordinate structures organized in $SS_1$ and $SS_3$.

The striking similarity between the eigenfunctions of the latent space and the eigenfunctions of the atomic models can be rationalized as follows. If the range of a single body rotation is moderate ($\lesssim 30°$), the distance $D_{ij}$ between any two states $i$ and $j$ within this range is to high accuracy[†] $D_{ij} = \Theta_{ij}(\sum_{k=1}^{m} r_k^2)^{1/2}$, where $r_k$ is the distance of atom $k$ away from the rotation axis, $m$ the number of atoms of the body, and $\Theta_{ij}$ the angular difference between the states. Therefore, $D_{ij}$ is directly proportional to $\Theta_{ij}$. If there are multiple independent body rotations (i.e., CMs) present, the individual distances add in quadrature as in a Euclidean space. Also not investigated in this paper, the linearity holds for body translations as well, where the distance is directly proportional to the magnitude of the translation. Thus, the agreement between the eigenfunctions of the latent space and the ones of the atomic models is a consequence of the linear relationship between distance and the multi-body motions, rotations and translations.

[†] This approximation is justified because the rotation matrix has only linear terms in rotation angle, provided the rotation angle is small.

#### C. Eigenfunctions of the 3D electron density maps

We next demonstrate that the properties of manifolds (as seen in the previous subsection for the 3D atomic-coordinate structures) significantly change when the data representation of their underlying states is altered. Specifically, we investigate how the conformational relationships between states are changed when representation by atomic coordinates is transformed into one by 3D electron density maps (EDMs; e.g., Fig. S3). To this end, we generated the EDMs for each of the 3D atomic-coordinate structures for all previously defined state spaces, as is described in our synthetic-generation protocol (section SM-I). We calculated the pairwise Euclidean distances between these EDMs in $\Omega_{\text{EDM}} \subset \mathbb{R}^V$, with $V$ the number of voxels in an EDM, and performed an embedding via the DM framework. Overall, over a wide range of Gaussian bandwidths, the structure of the resulting eigenfunctions was very similar to the structure of eigenfunctions retrieved for the atomic models in the small Gaussian bandwidth regime. Importantly, these eigenfunctions were still of the form $\psi_{vw}$ (see subsection B), with subspaces having no significant appearance of eigenfunction misalignments.

However, there are a few attributes to consider that distinguish the manifolds obtained for EDMs from those retrieved for the previous data types. First, the difference between small and large Gaussian bandwidth regimes was much less drastic, such that the cosine eigenfunctions appeared in both regimes. For small Gaussian bandwidth regimes (i.e., a few orders of magnitude below the value determined by the bandwidth estimation method $\varepsilon_{\star}$, as defined in section SM-XI), we found that the $CM_2$ parabola-housing subspace was buried deeply in low-ranking eigenvectors (e.g., $\Psi_8$ and higher), with

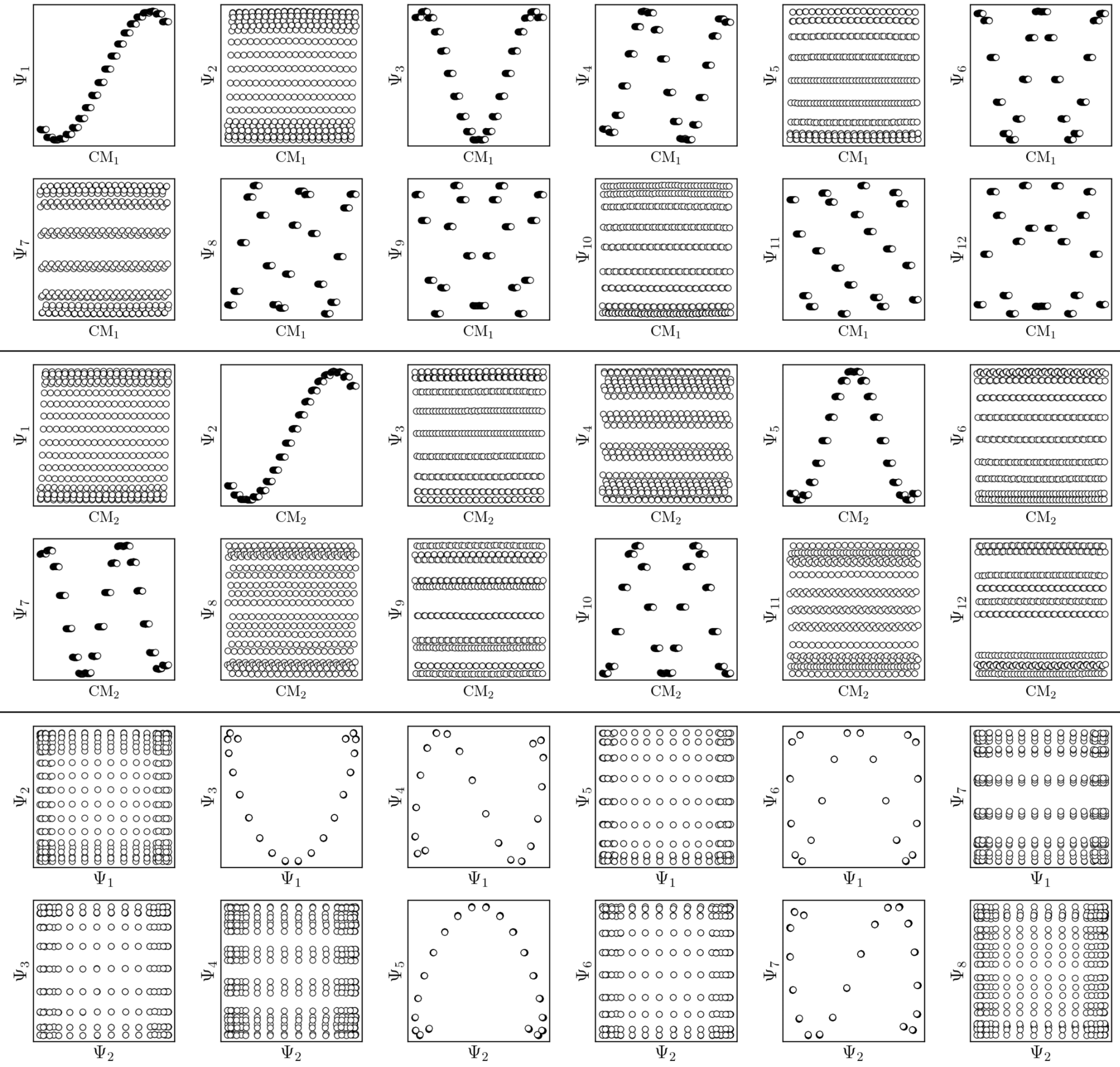


FIG. S23: The results of the DM embedding of EDMs (pure signal) from $\text{SS}_2$ are shown. Leading eigenfunctions as indexed by $\text{CM}_1$ and $\text{CM}_2$ are displayed in the first four rows, followed by their composites. Overall, there is near-perfect alignment of these eigenfunctions with the initially-obtained eigenvector basis, such that no rotations are required. As a final note, the pronounced inward curling at the boundaries of certain subspaces (e.g., $\{\Psi_1, \Psi_3\}$) is due to insufficient sampling, as was identically encountered in our analysis of the embeddings of manifolds obtained from subsequent projections of these EDMs.

numerous $\text{CM}_1$ parabolic harmonics occupying the subspaces in between. In addition, eigenvectors with cross terms $\Psi = \{\psi_{vw} \mid v, w \neq 0\}$ were found scattered mostly in mid-range positions (e.g., $\Psi_{12}$ and higher). Since these properties were not observed in the embeddings of the atomic-coordinate structures, we conclude they are a result of a change in the metric.

In contrast, for larger Gaussian bandwidth regimes (i.e., near and significantly above $\varepsilon_\star$), eigenvectors with cross terms were buried in much deeper subspaces (e.g., $\Psi_{34}$ and higher), with the majority of leading eigenvectors housing content exclusively for either $\text{CM}_1$ ($w = 0$) or $\text{CM}_2$ ($v = 0$). Thus, the corresponding CM parabolas for these $\varepsilon_\uparrow$ regimes typically occupied the first two subspace rows ($\{\Psi_1, \Psi_i\}$ and $\{\Psi_2, \Psi_j\}$), with respective harmonics positioned in trailing subspaces. These CM parabolas also had a near-perfect distribution of points, whereas for the $\varepsilon_\downarrow$ regime, the distribution of points had noticeably less precision to the ideal form. We note that the embeddings obtained above and below these regimes were incoherent in form.

We conclude that the eigenfunctions obtained from the larger Gaussian bandwidth regime would be preferred for several reasons. First, the desired $CM_1$ and $CM_2$ parabolas occupy leading subspaces and are thus easily identifiable. The paucity of leading cross-term eigenfunctions is also convenient, since they provide no useful information for our analysis, while also obfuscating our search for desired subspaces. Additionally, the geometric structure of all subspaces obtained via $\varepsilon_\uparrow$ consistently appears much closer to the ideal form. In Fig. S23, we display the DM eigenfunctions obtained from this regime for the $20 \times 20 = 400$ EDMs occupying $SS_2$. Subspaces indexed in the $CM_1$ reference frame (rows one and two) and the $CM_2$ reference frame (rows three and four) are displayed, as well as a set of leading composites of these eigenfunctions in the rows that remain.

Importantly, as there was no Gaussian bandwidth value that could "recover" the preferred Legendre-like form, it appears that this feature is "lost in translation" upon transformation from atomic models to EDMs, since the metric is changed. (To note, the curved geometry formed by cosines was also in close agreement with the results of applying PCA on this same dataset). As a main agent for this distinction, the distance measure pertaining to EDMs is fundamentally different from the one of the 3D atomic-coordinates. Instead of the 3D coordinate points that stand for the atomic positions of each structure, the data for each EDM is represented by a 3D array of values, one at each voxel. A key difference then, is that in the latter case, the displacement of atoms are no longer accounted for individually. Instead, every voxel in the data of one state is now compared to those same voxel locations in the data structure of another state, with only changes in the value at each voxel entering the distance measure.

Hence, while the eigenfunctions are similar, the relationship between states in these two data types is fundamentally different. To demonstrate this change, Fig. S24 shows a comparison of the pairwise distances between states as calculated for the rectangular latent space, atomic-coordinate structures, and EDMs. As noted in the caption, by assessment of the close similarity between the distances from the latent space and atomic models, we can infer that these two data types are both confined to the rectangular manifold $\Omega_R$ (albeit of different sizes). As a consequence, we observed that their eigenfunctions are nearly identical. In contrast, we see that the distances from the EDMs are starkly different from the rectangular pattern, where neighboring states are spatially arranged via an asymptotic-like trend. From these findings, we must infer the corresponding data live in an altogether different manifold. Although the explicit geometric form of $\Omega_{\text{EDM}}$ is unknown, we have shown that the spectral properties of the Laplacian in $\Omega_{\text{EDM}}$ are essentially preserved via the mapping from the latent space. While detailed knowledge of $\Omega_{\text{EDM}}$ is certainly of interest, it is inconsequential since our analysis only requires an understanding of the eigenfunctions of a manifold, and not necessarily its exact shape.

#### D. Eigenfunctions of the 2D projections

Since a detailed description of the eigenfunctions of 2D projections is provided in the main text, we continue this current narrative only as it pertains to the relationship of the eigenfunctions of the LBO on $\Omega_{\text{PD}} \subset \mathbb{R}^P$ with those from previously-established models (i.e., rectangular Euclidean latent space, atomic-coordinate structures and EDMs). For similarities, as was observed for the EDMs, we found that eigenfunction characteristics could be broadly classified into two classes via either a small or large Gaussian bandwidth regime. In either regime, the eigenfunctions of the PD manifolds were again of the form $\psi_{vw}$, such that only cosines emerged. The lack of the Legendre-like form and a similar asymptote-like appearance of distances between images suggests that the PD states in $\Omega_{\text{PD}}$ reside on a manifold similar to $\Omega_{\text{EDM}}$.

The overall difference between eigenfunctions obtained via $\varepsilon_\downarrow$ and $\varepsilon_\uparrow$ was also much more impactful for PDs than for the EDMs. In the small Gaussian bandwidth regime, $CM_2$ subspaces had a severely suboptimal point distribution, such that, in some PDs, identification of the $CM_2$ parabola was completely obstructed. These $CM_2$ subspaces were also buried in trailing eigenvectors, and interspersed among those with cross terms (i.e., $\Psi = \{\psi_{vw} \mid v, w \neq 0\}$). We also note that the value determined by the bandwidth estimation method ($\varepsilon_\star$) fell within this regime, making it a suboptimal choice for pristine data. In contrast, the large Gaussian bandwidth regime (i.e., one order of magnitude larger than $\varepsilon_\star$ and spanning numerous orders of magnitude above it) was superior in every sense, with $CM_1$ and $CM_2$ eigenfunctions having ideal point distributions and corresponding subspaces occupying leading eigenvectors. As well, the cross-term eigenfunctions were present only in far trailing eigenvectors (e.g., $\Psi_{31}$ and higher), and would not be obstructive during the analysis. Briefly, we note that upon introduction of noise (SNR = 0.1) and duplicate states ($\tau = 5$), $\varepsilon_\star$ was instead an optimal choice (along with numerous orders of magnitude above it), while anything below this range was completely inadequate.

For either Gaussian bandwidth regime, we found that the eigenfunction misalignments can emerge—and with varying magnitude—depending on the projection direction. Since previously we have shown that no such property is apparent in the manifold embeddings generated from the 3D EDMs from which these PDs originate, it is clear that the emergence of these eigenfunction misalignments is tied to PD disparity. We hypothesize that, as different 2D projections are taken of the EDMs via $p : \Omega_{\text{EDM}} \mapsto \Omega_{\text{PD}}$, the geometry of $\Omega_{\text{EDM}}$ can become contorted due to the change of pairwise interatomic distances resulting from foreshortening in projection (Fig. S25), such that the apparent span of one CM

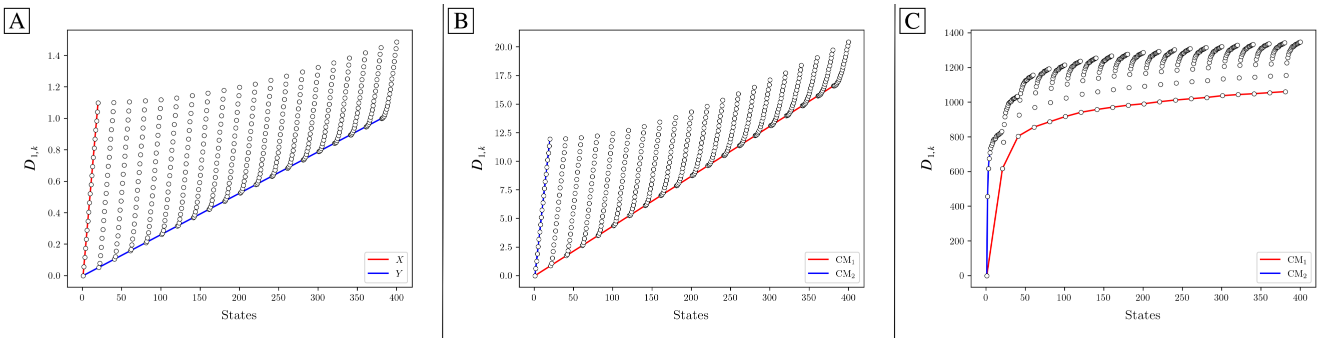


FIG. S24: The first row of the distance matrix $\boldsymbol{D}$ is plotted for the rectangular Euclidean space [A], the 3D atomic-coordinate structures in SS$_2$ [B], and the EDMs in SS$_2$ [C]. Given our ordering of states, the first row $D_{1,k}$ corresponds to the pairwise distance calculated between state 01_01 and all 400 states. For [A], which was calculated for a rectangular domain $\Omega_R \in [0, 1.1] \times [0, 1] \subset \mathbb{R}^2$, one can identify the distance of the first state $(x_1 = 0, y_1 = 0)$ to all other coordinates, such that the red line depicts the base of the rectangle (with maximum distance $D\{(x_1, y_1), (x_{20} = 1.1, y_{20} = 0)\} = 1.1$), and the blue line depicts the rectangle's left-hand side boundary (with maximum distance $D\{(x_1, y_1), (x_{381} = 0, y_{381} = 1)\} = 1$). In [B], a similar rectangular pattern arises for the RMSD values calculated between atomic models. The pattern in [C], however, is starkly different from [A] and [B], such that no rectangular (or rectangle-like) domain could be drawn to reproduce this trend.

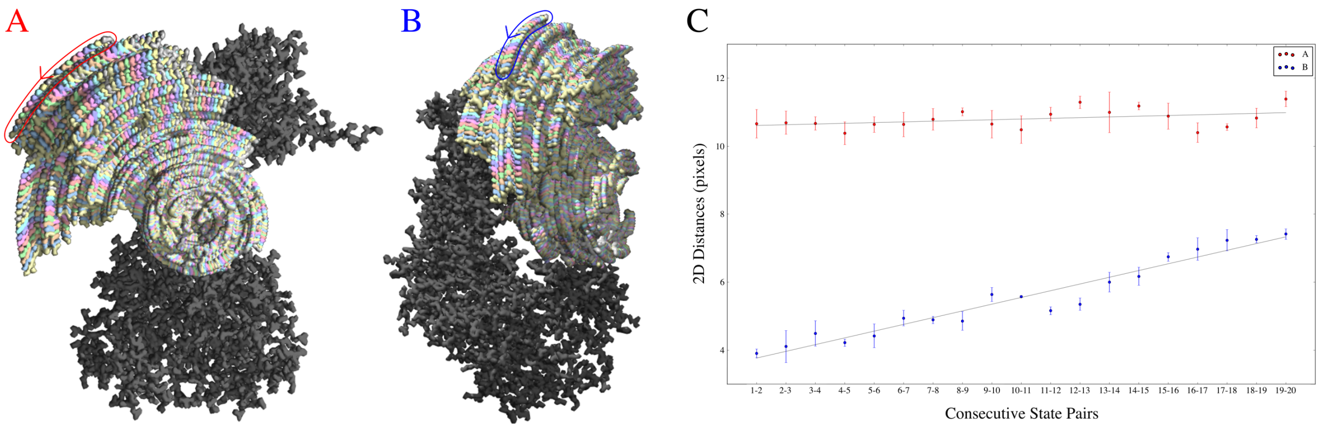


FIG. S25: Here we provide intuition for the emergence of foreshortened distances due to taking 2D projections of 3D EDMs. Two orthographic views of 3D models in the directions of two PDs are shown in [A] and [B], each composed of 20 overlaid 3D volumes from CM$_2$ (i.e., one degree of freedom). The 2D distances (in pixels) were measured between the peripheral ends of each consecutive states' rotated subunit (as seen in the red and blue encircled regions). After conducting three sets of 2D distance measurements on each region in image [A] and [B] independently, the mean distances were plotted with error bars representing standard deviation, followed by linear regression [C]. As can be seen, although the distances between states in the object's 3D form is constant, when projections are taken, these distances can strongly vary based on the current 2D view. While the Euclidean distance matrix calculated in the DM framework is less intuitive and records these changes on a pixel-by-pixel basis for the entire image, we anticipate analogous relationships to emerge there based on PD and CM.

to another depends on PD.

Thus, throughout these sections, we have demonstrated how the embeddings of the manifolds containing the same conformational information change depending on how the data is represented. Nevertheless, we were able to closely approximate the observed $\Omega_{\text{PD}}$ eigenfunctions when allowing for eigenbasis rotations of the form

$$\Psi_i = \cos(\theta)\cos(v\pi x) + \sin(\theta)\cos(w\pi y) = A\psi_v + B\psi_w$$

Using this expression, we generate graphs in Figure S26 which are able to analytically reproduce the heuristically-derived subplots in Figure 3 and Figure 4. Apart from a few minor discrepancies, we observed an outstanding agreement between our analytical functions and the findings from our heuristic analysis. The discrepancies that do emerge can be understood as additional, small-scale perturbations which are currently unaccounted for in our general eigenfunction expression.

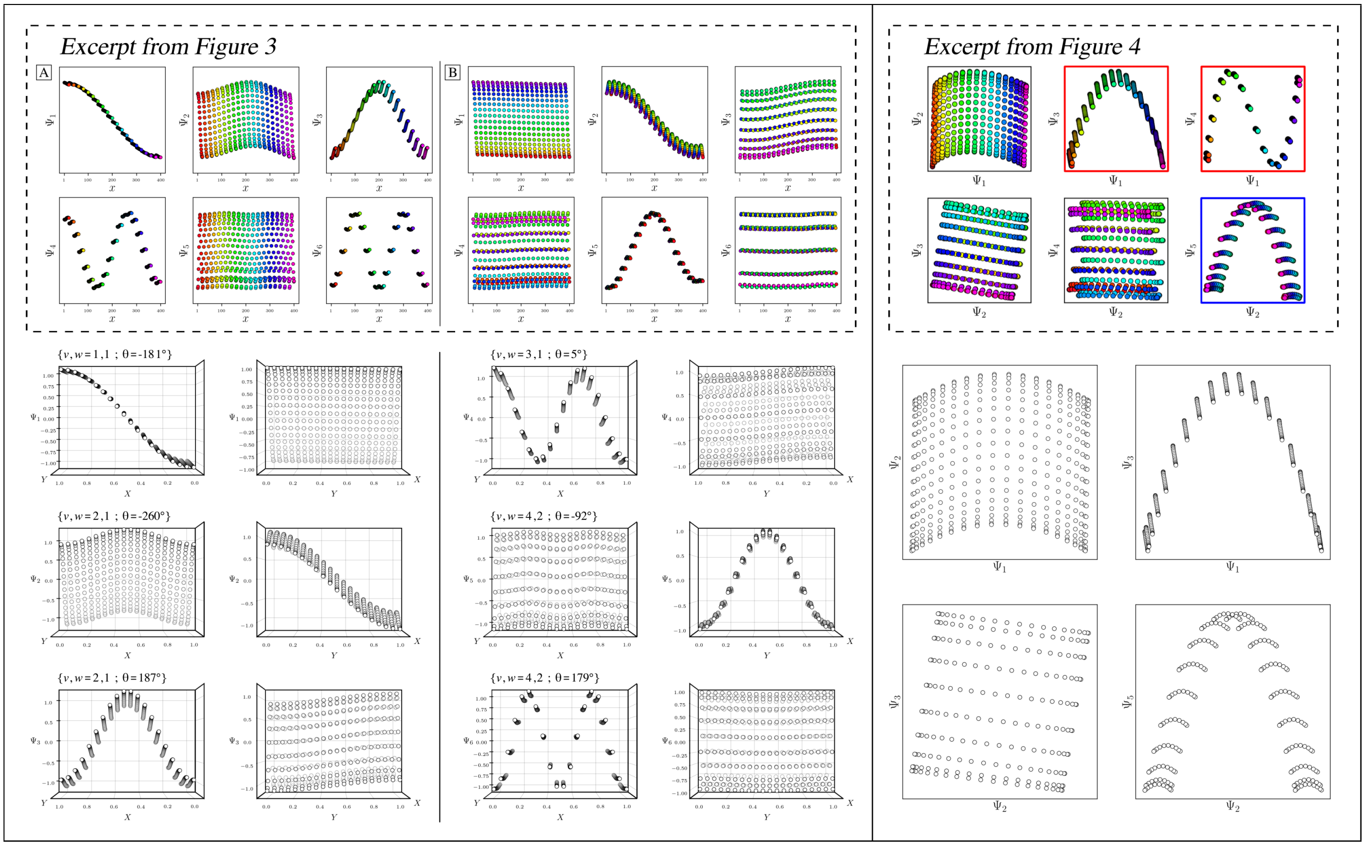


FIG. S26: Comparison of analytically-generated functions with the eigenfunctions previously obtained for $PD_1$ (Fig. 3 and Fig. 4). For each pair of subplots, values for $\theta$ were approximated by eye. Our approximations share a remarkable similarity with earlier heuristic results, and are able to account for geometric minutiae previously unaccounted for, as well as larger-scale rotations seen in the composite of eigenfunctions. Discrepancies can be seen in the slightly tilted appearance of parabolas in $\Psi_3$ and $\Psi_5$ of Fig. 3, as well as the clumping of points as observed in the $CM_2$ reference frame of $\Psi_6$.

### SM-XV. BOUNDARY CONDITIONS OF THE LBO

Here we lay out a strategy for dealing with molecular machines that exhibit each of their domain motions along an independent and mutually unrestricted sequence of quasi-continuous states. The set of all $n$-wise combinations of these bounded intervals (one for each conformational motion) produces an $n$-dimensional shape with a rectangular boundary. In section SM-XIV, we have shown that the corresponding Laplacian eigenfunctions are well defined. However, in general, analytically solving the Laplacian for any arbitrary boundary is impossible. Eigenfunctions can change drastically depending on the boundary, and are analytically only known for certain elementary shapes, such as rectangles, discs, ellipses and special triangles[28,31]. On the other hand, geometric machine learning approaches can obtain solutions numerically, in principle for any boundary. However, such geometric machine learning methods still require the boundary to be known *a priori*. For systems with unknown boundaries, the problem is intractable.

As the set of all possible molecular machines is unfathomably complex, it is unlikely that one single algorithm could ever be so versatile as to anticipate every possible instance. Instead, we are interested in casting a wide enough net to capture the dynamics of a large portion of these systems, which we surmise operate within rectangular boundaries of an $n$-dimensional latent space of multi-body motions. However, one can still imagine all sorts of other situations, such as a system where one domain blocks—via steric hindrance—another domain from its full range of motion in a specific region of the state space (Fig. S27-B). Importantly, our requirement of adequate coverage (as detailed in Discussion) excludes the case of obtaining poorly sampled data from a rectangular domain, which would allow any number of arbitrary shapes to emerge. This exclusion also holds for state spaces with "holes" (i.e., interior boundaries)[28], where the occurrence of certain states is forbidden due to energetic restraints. To better understand the effects of these boundary challenges, we have created a 2D state space with an octagonal domain (noting the Laplacian eigenfunctions of the octagon is an analytically unsolved problem), which was achieved by eliminating states at the four corners of our standard rectangular domain (Fig. S27-A). To circumvent the occurrence of eigenfunction misalignments due to PD disparity, which may complicate the interpretation of the boundary influence, we embed the 3D electron density maps instead of 2D projections.

The corresponding manifold embedding obtained from this octagonal state space is shown in Fig. S27-C, which features a number of deviations from the canonical rectangular eigenbasis (Fig. S23). Manually, we attempt to find a transformation from the octagonally-derived eigenbasis (Fig. S27-C) to the rectangular form (Fig. S23) by intuiting a collection of suitable rotation operators. Indeed, we are able to show that such a transformation is possible, up to some level of uncertainty (Fig. S27-D). Thus, it is not that the eigenfunctions are dramatically changed by the imposed boundaries, but, instead, that the eigenvectors can now contain multiple cosine terms. We note that both the indices and number of rotation operators required for this transformation deviated from our findings on eigenfunction realignment performed on rectangular state spaces (see section SM-XVI), with the collection of decisions required now more complex. Thus we believe this instance only further motivates the need for a future comprehensive method for estimating the preferred eigenbasis rotations. Given our own observations, a maximum-likelihood approach may be better suited for these demands, with such a study deserving of the scale delivered for other ManifoldEM subproblems, such as Belief Propagation[32].

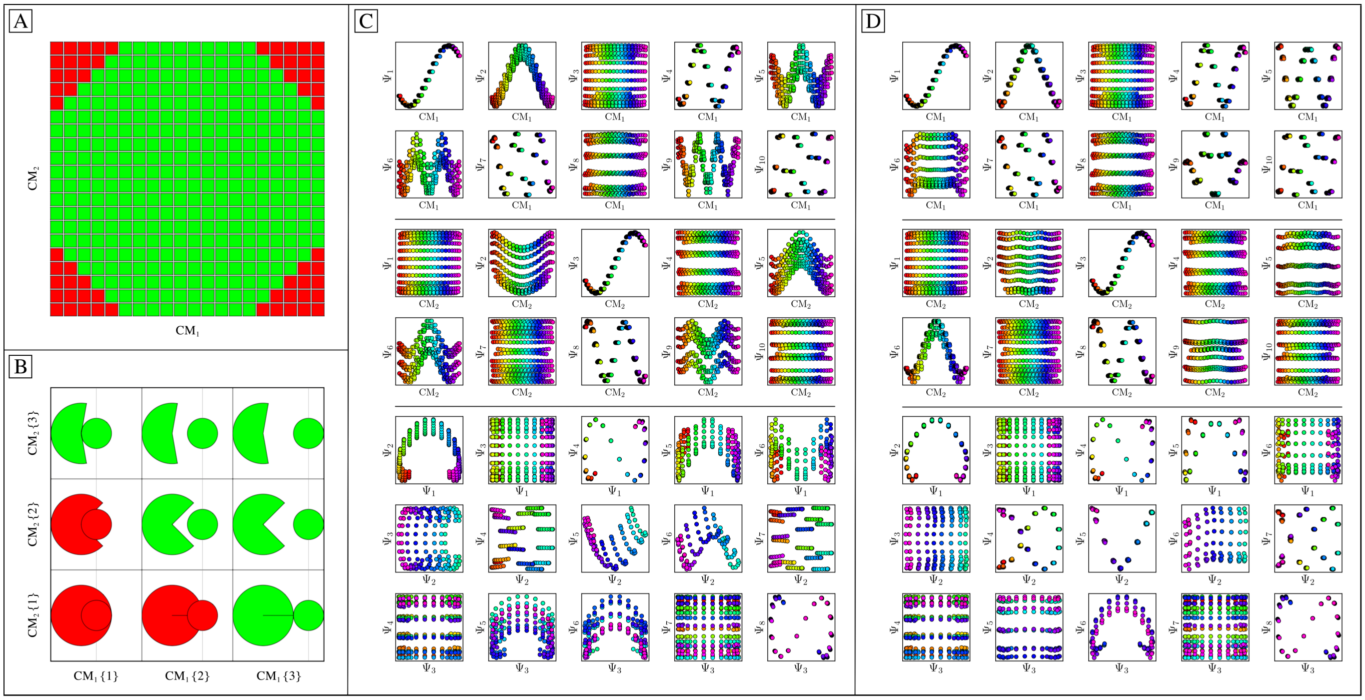

FIG. S27: Analysis of the eigenfunctions [C] associated with the octagonal state space [A] of EDMs. The initial $20 \times 20$ rectangular state space is displayed in [A], where red boxes illustrate states that were removed to form an octagonal grid. The schematic in [B] provides some context for the possibility of a non-rectangular state space, which can be envisioned as a top-down view of (1) a large domain that opens and closes, and (2) a small domain that translates left and right. Naturally, while the larger domain is in a closed or half-closed state, the smaller domain is impeded from accessing a subset of its possible states, and vice versa. The eigenbasis obtained after application of a set of high-dimensional rotations (of dimension $d = 15$) is shown in [D]. The required operators were estimated by hand, and included several large and small transformations: $\{R_{5,6}(40^\circ), R_{2,6}(-15^\circ), R_{2,5}(3^\circ), R_{2,9}(4^\circ), R_{6,9}(20^\circ), R_{6,12}(-25^\circ), R_{2,11}(-6^\circ), R_{9,11}(25^\circ), R_{9,15}(5^\circ), R_{6,11}(3^\circ)\}$. We note these calculations are for completeness, and not a minimal set for a desired subspace. Still, we were unable to perfectly decouple the $\{\Psi_3, \Psi_6\}$ subspace. As $\Psi_3$ appeared well behaved in the $\text{CM}_2$ reference frame, it is possible that either the $\Psi_6$ eigenfunction was fundamentally altered due to the new boundaries, or our choice of an earlier operator trapped us in a suboptimal solution.

### SM-XVI. EIGENFUNCTION REALIGNMENT

In this section, we demonstrate application of our eigenfunction-rotation algorithm and present the operators $R_{ij}$ needed to counter-rotate each CM subspace for our final analysis. In addition, we provide here intuition for the aforementioned elimination procedure, which we apply to remove parabolic harmonics. These strategies are formulated by inductive reasoning based on patterns observed across hundreds of $\Omega_{\text{PD}}$ embeddings.

We start by identifying the eigenvector indices corresponding to the first CM subspace. Since a macromolecule's most prominent conformational signal as seen from the current viewing angle must always be the leading factor in the embedding of a PD manifold, we expect the leading CM subspace to appear in the set $\{\Psi_1, \Psi_i\}$, and use a least-square fitting strategy to discover the second eigenvector $\Psi_A$ (see movie M7). Once this first parabola-housing subspace is identified, we can eliminate all subspaces $\{\Psi_A, \Psi_i\}$ as housing harmonic information corresponding to this leading CM at $\{\Psi_1, \Psi_A\}$. We can additionally eliminate $\{\Psi_B, \Psi_A\}$ in the next non-harmonic eigenvector row as a candidate for the location of the orthogonal CM parabola, which is easily understood via examination of movie M6. Subsequently, once the second CM subspace is identified by similar least-square measures (e.g., $\{\Psi_B, \Psi_C\}$ if $C \neq A$), the previous elimination procedure can be repeated to identify the location of this CM's corresponding harmonic spanning $\{\Psi_C, \Psi_i\}$, and so on, as is afforded by the geometric fidelity to the LBO (and thus ability for parabolic fitting) of these higher-order subspaces. As it is currently designed, this elimination procedure is applied only to ensure removal of the lowest-order harmonics[19], such that the leading set of potential CMs discovered through this approach should always be unique.

Eigenvector indices can be additionally used to narrow down the rotational operators $R_{ij}$ required to adequately rotate each 2D subspace. Specifically, for two degrees of freedom, all pairwise combinations of the eigenvector indices corresponding to the first and second parabola-housing subspace (excluding parabolic harmonics) determine the set of required $R_{ij}$ operators for $n = 2$. As a concrete example, we consider a 6-dimensional case where the $\text{CM}_1$ parabola is found in the $\{\Psi_1, \Psi_3\}$ subspace, the $\text{CM}_2$ parabola in the $\{\Psi_2, \Psi_5\}$ subspace, and the first $\text{CM}_1$ parabolic harmonic in the $\{\Psi_3, \Psi_6\}$ subspace. Then the 15 available rotation operators can be narrowed down to just the use of some combination of $\{R_{1,2}|R_{1,5}|R_{2,3}|R_{3,5}\}$, and applied in order of highest to lowest eigenvector significance while measuring minima via the aforementioned histogram routine. To note, while it is true that finite rotations about different axes do not commute[33], we found only minor deviations in the final orientation of each eigenbasis, given different permutations in the sequence of these four operators. Once the location of these parabola-housing subspaces are established, harmonics can alternatively be discarded if duplicate $R_{ij}$ indices emerge under this scheme. The emergence of duplicate indices can be understood in the current example when taking pairwise combinations of the leading parabola-housing subspace with the $\{\Psi_3, \Psi_6\}$ harmonic, which will incur the trivial $R_{3,3}$ operator.

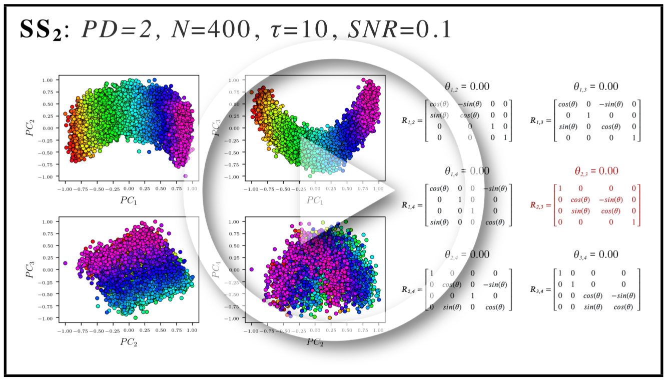


MOV. M5: Effect of applying a 4D orthogonal rotation to the 4D subspace (shown here using four projections of that subspace) obtained from $\text{PD}_2$ (in $\text{SS}_2$ with $\tau = 10$ and SNR $= 0.1$). The six rotation matrices required for the rotation of a 4D subspace are shown on the right, where the number of matrices for a $d$-dimensional subspace scales via $d(d-1)/2$. (Note that the minimum dimensionality of the rotation matrix used must match the dimensionality of the subspace required to encompass all parabolic modes present in the embedding). As can be seen, by only applying rotation operator $R_{2,3}$ with 0.5 radians (28.65°), both $\text{CM}_1$ and $\text{CM}_2$ parabolic modes are corrected (preserving all distances between points) such that they reside completely in the plane of $\{\text{PC}_1, \text{PC}_3\}$ and $\{\text{PC}_2, \text{PC}_4\}$, respectively. `https://www.dropbox.com/s/c9b6vcjeffwziqp/M5_EigRot_4D.mp4?dl=0`

As a final note for this section, there exists a rare occurrence that must be accounted for when the initial eigenbasis of the $\Omega_{\text{PD}}$ embedding is severely misaligned from the preferred coordinate system. In this event, we have observed that CM subspaces cannot be initially located by the presence of favorable $R^2$ values generated from least-squares fits. We encountered this problem for one manifold of the 504 analyzed (0.2%) across both data-type II (using three great circles) and data-type III (using one great circle). This error is easily recognized by the presence of a significantly low $R^2$ value (less than 0.1) for each 2D subspace of a given PD manifold. In such an event, we suggest rotating the eigenbasis by 45° using the $R_{1,2}$ operator and recalculating least-squares fits. As additional cases emerge, it is likely that a more comprehensive strategy may be required.

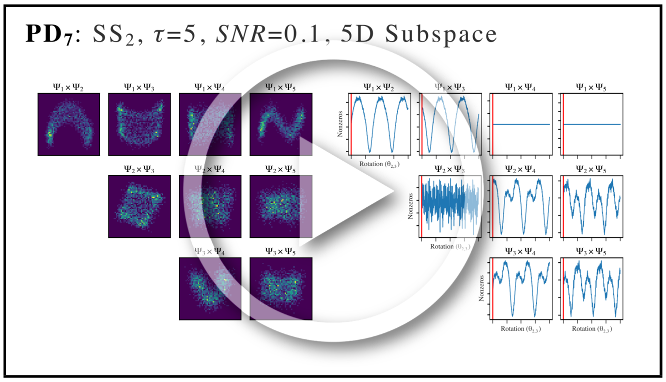


MOV. M6: A movie displaying the 2D histogram approach for finding optimal angles and corresponding parabolic modes. Specifically, the effect of an incremental 4D rotation operator $R_{2,3}$ on each 2D subspace is shown. During these rotations, each 2D subspace exhibits a unique profile which can be characterized by the number of nonzero bins in the corresponding 2D histogram as a function of angle. Note the appearance of specific patterns that emerge between these 2D subspaces as rotations are performed, which we leverage in our algorithm to procedurally eliminate subspaces from potential misuse. `https://www.dropbox.com/s/ekh44n66x5j7sz5/M6_EigRot_Hist.mp4?dl=0`

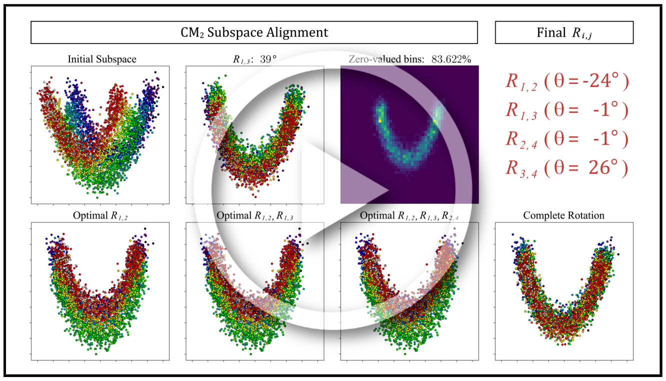


MOV. M7: An example PD from datatype II is chosen to demonstrate the inner workings of our eigenfunction realignment algorithm. Here, a $d = 5$ dimensional subspace is first isolated, with each 2D subspace therein assigned an $R^2$-value based on least-square fits. Given presence of adequate fits, the parabola-housing subspace in each eigenvector row is determined via the best $R^2$ value, with the corresponding eigenvector indices used to procure the four rotational operators (of 10, for $d = 5$) required to align each point cloud with the plane of its subspace. We next demonstrate the generation of 2D histograms as these operators are exercised to determine the optimal angles, as previously detailed in Movie M6. As can be visually assessed, slight inaccuracies may emerge during the histogram optimization (typically no more than 5°), but prove insignificant for downstream procedures. `https://www.dropbox.com/s/u6qqrc6bmm5881t/M7_EigRot_Rij.mp4?dl=0`

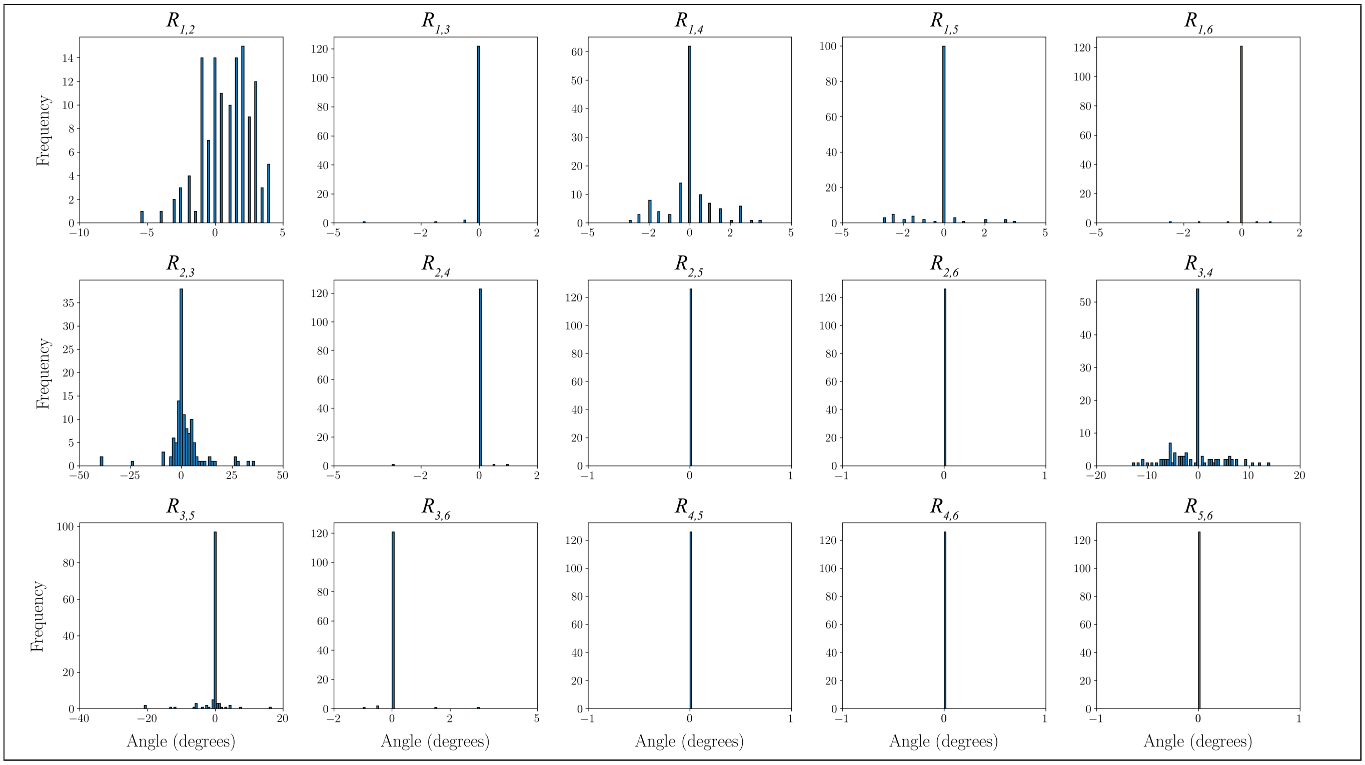


FIG. S28: Frequency of $R_{i,j}$ angles ($d = 6$) required to counter-rotate each $CM_1$ subspace within the 126 PDs along the $S^2$ trajectory spanning half of one great circle. A total of 15 subplots are displayed corresponding to the $d(d-1)/2$ rotation operators necessary for this lower-dimension analysis. As can be seen, the magnitude of rotations required for a significant portion of PDs was substantial. To note, angular trends for each operator are highly subject to the choices of $S^2$ trajectory and CM.

### SM-XVII. RECOMBINATION OF CONFORMATIONAL STATES

The following figure provides intuition for our method of generating multidimensional free-energy landscapes and corresponding 3D movies using the ESPER intersection of image-indices within each PD manifold. For this schematic, we have simplified the problem from that of accounting for a parabolic surface (as observed throughout this study) to a plane; however, the following intuition remains the same in either case.

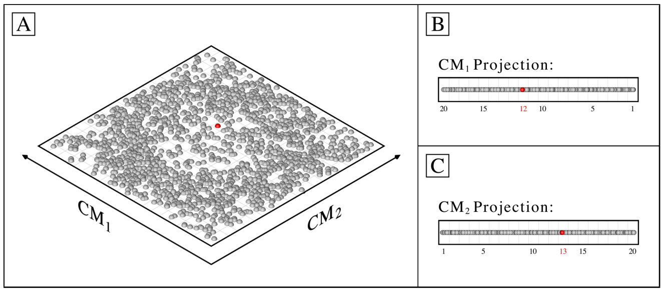


FIG. S29: A schematic to provide intuition for our intersection of image-indices approach, which simplifies the complex parabolic features in our observed PD embeddings into linear ones, on a plane. Let the red point near the center of the plane represent an image with image index $p_i$, such that $p_i$ belongs to both $\mathrm{CM}_1\{12\}$ and $\mathrm{CM}_2\{13\}$, and thus $p_i \in \mathrm{CM}_1\{12\} \cap \mathrm{CM}_2\{13\}$. This point, along with all others in the intersection $\mathrm{CM}_1\{12\} \cap \mathrm{CM}_2\{13\}$, is used to define an occupancy and reconstruct a 3D density map for the respective state 12_13.

Just like the plane in Fig. S29-A, PD images (represented by points) are organized along $n$ orthogonal degrees of freedom (CMs) on a higher-dimensional hypersurface. For our needs, this hypersurface can be approximated as a parabolic surface. Given the aforementioned uncertainty and difficulty in identifying and mapping this surface directly, we can instead refer to its set of $n$ orthogonal projections (e.g., Fig. S29-B and Fig. S29-C), which can be found and mapped with less difficulty. In the case of the plane—as in the case of our simplified illustration—these subspaces are 1D, while for $\Omega_{\mathrm{PD}}$ embedding, a 2D subspace is required to adequately capture each parabolic component. Recall that we identify these subspaces after performing eigenvector rotations to align the parabolic surface, such that only the CM parabola is visible in each of the respective 2D subspaces. Once located, we straighten each CM trajectory in each of these lower-dimensional projections into a 1D trajectory, such that the parabola is transformed into rectilinear form. Next, we partition the points separately into $\beta$ contiguous bins (here, $\beta = 20$), and collect the set of image indices falling into each bin. Note that the size of the bin effectively defines the precision to which we can locate each point on the plane, and determines the range of images falling within each state for our final outputs.

As a result, we are left with $n\beta$ sets of image indices combined across each set of CM coordinates for each PD. For ease of explanation in the following notation, assume $n = 2$. Next, we construct an empty $\beta \times \beta$ (i.e., $\beta^n$) array $\boldsymbol{P}$ and fill each element $P_{x,y}$ with the set of all image indices in the intersection $\mathrm{CM}_1\{x\} \cap \mathrm{CM}_2\{y\}$, where $x$ and $y$ are bins from $\mathrm{CM}_1$ and $\mathrm{CM}_2$, respectively. Since manifolds from each PD were obtained independently, we must also correct for *sense* (the directionality of the CM coordinates) as we accumulate indices in $\boldsymbol{P}$. At the end of this procedure, we sum the total number of entries in each $P_{x,y}$ to form a $\beta \times \beta$ occupancy map (which can then be converted into a free-energy landscape via the Boltzmann relation). We additionally use the indices of images within each $P_{x,y}$ to reconstruct a 3D density map for the set of corresponding images; in this example, producing 400 3D density maps in total. Naturally, this construction can be easily extended to three or more degrees of freedom.

Thus, given only a set of CM subspaces—each a parabolic trajectory defining an orthogonal degree of freedom—and with knowledge of the higher-dimensional relationship between them (i.e., the parabolic surface, as determined throughout our heuristic analysis), we can reconstruct that joint geometrical relationship using only the intersection of image indices obtained in all pairwise combinations of bins from straightened CM coordinates. In effect, this procedure only requires that we collect the indices of images, without the need to integrate them into 1D occupancy maps. This is in contrast to the previous ManifoldEM methodology employing NLSA, which discards the original image indices. This action carries a price, and must be reversed by performing a lengthy tomographic reconstruction using the 1D occupancy maps to obtain the 2D distribution (for more information on the NLSA strategy, see the following section SM-XVIII).

### SM-XVIII. OVERVIEW OF NLSA

Nonlinear Laplacian spectral analysis[34] (NLSA) is primarily used for noise reduction during the extraction of the conformational signal contained in each PD. NLSA is applied independently on each member of a leading set of $\Omega_{\text{PD}}$ eigenvectors in order to assess the "meaning" of each in terms of housing potential CMs of interest. For each of these eigenvectors, NLSA is performed as follows. First, the raw images are concatenated along a chosen eigenvector to produce so-called "supervectors"[35]. These supervectors are then embedded to form a new set of eigenvectors in a different space. This results to high accuracy in a 1-dimensional manifold, with known eigenfunctions $\cos(k\pi\tau)$ parameterized[35] by a conformational parameter $\tau$ (separate from the use of $\tau$ in our analysis). This enables the estimation of a density of points as a function of $\tau$ together with an ordered sequence of noise-reduced (i.e., interpolated, via the supervectors) 2D images. These 2D images can be arranged to form a 2D NLSA movie, designed to represent the conformational signal corresponding to the eigenvector chosen from the initially-embedded PD manifold. Once a set of 2D NLSA movies have been constructed along each of the leading $\Omega_{\text{PD}}$ eigenvectors independently, supervised identification of "meaningful" CM information is next required.

When only one degree of freedom is desired (or available), the 2D movies corresponding to the same CM content in different PD manifolds can be further compiled across $S^2$ to reconstruct 3D density maps and thus a 3D movie representing the CM. The NLSA procedure is more complicated when two (or more) degrees of freedom are desired. After supervised identification of two CMs, their respective eigenvectors for the current $\Omega_{\text{PD}}$ are used to isolate a 2D subspace therein. On this $\{\text{CM}_1, \text{CM}_2\}$ subspace, NLSA is performed independently along the directions of (typically) 180 uniformly-spaced radial lines in the range $(0 \leq \theta \leq \pi)$. This yields a collection of point densities (i.e., 1D occupancy maps) $n(\tau, \theta)$ for each $\theta$. The collection of these 1D maps for all $\theta$ constitutes the 2D Radon transform of a yet unknown 2D density map (i.e., the desired 2D occupancy map). An inverse Radon transform is then applied to reconstruct the 2D density map. In addition, NLSA also retrieves the noise-reduced images at each point in this map. To note, one of the rationales in the way NLSA-based retrieval of images is organized is that it normalizes the initially unknown rates of change in different CM directions[35]. As in the 1D case, this procedure must next be performed for the eigenvector pairs corresponding to $\{\text{CM}_1, \text{CM}_2\}$ in all other embeddings of $\Omega_{\text{PD}}$, from which noise-reduced 3D density maps can be reconstructed to form 3D movies of concerted conformational motions.

### SM-XIX. COMPARISON OF RESULTS FROM ESPER AND NLSA

Here we compare the outputs of ESPER and NLSA for a few example PDs from our final analysis dataset (126 PDs, SNR, CTF and nonuniform occupancy map, as previously detailed), with these PDs selected based on the properties of both their visual appearance and respective embedded geometries. Importantly, the same preliminary steps were performed for both frameworks (i.e., generation of identical manifolds and corresponding embeddings) before a branch in workflows. Also recall that ESPER includes the use of our eigenfunction realignment methodology on these manifold embeddings, whereas ManifoldEM with NLSA currently does not.

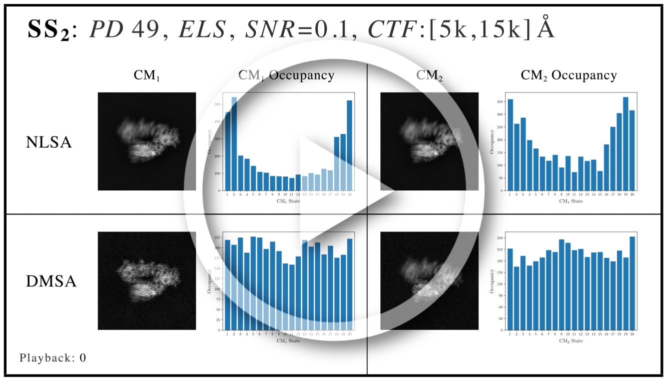


MOV. M8: For our comparison of ESPER and NLSA, three example PDs were chosen from our final analysis dataset for direct comparison of their 2D movies and corresponding occupancy maps. In the first segment of the movie M8, $\text{PD}_2$ represents the class of extremely well-behaved manifolds: near-perfect eigenfunction pre-alignment and irrelevant inward curling of either $\text{CM}_1$ or $\text{CM}_2$ subspaces. Next, $\text{PD}_{33}$ is a representative from the class of manifolds with appreciably unaligned eigenfunctions from the ideal eigenbasis, with the subspace of $\text{CM}_2$ here requiring a larger counter-rotation than $\text{CM}_1$. Finally, $\text{PD}_{49}$ required a minimal $d$-dimensional rotation (much like $\text{PD}_2$), but exhibited significant inward curling at the boundaries of its subspaces. The last segment of movie M8 demonstrates the final 2D movies (one per eigenfunction) output by NLSA for $\text{PD}_{33}$ and $\text{PD}_{49}$. `https://www.dropbox.com/s/qvpwsyqq6zc95qg/M8_Comparison_NLSA.mp4?dl=0`

As can be seen in movie M8, for the most well-behaved PD manifold, $\text{PD}_2$, there is general visual agreement in the 2D movies obtained from NLSA and ESPER for $\text{CM}_1$ and $\text{CM}_2$. However, there are noticeable discrepancies in the outputs between the results from these two techniques. Immediately apparent is the difference in quality of the domains under motion corresponding to the given CM. For ESPER, these domains are highly resolved across all frames produced, while for NLSA these regions are much less resolved and noticeably smeared out. Second, while the visual differences between frames of ESPER appear to evolve at an even pace, differences in frames appear less emphasized near the beginning and end of the NLSA movies, as if the CM movies were decelerating near these regions. In addition, the NLSA occupancies share little resemblance to our ground truth, with accentuated errors near the boundaries. Similar boundary problems exist but are significantly less present for ESPER occupancy maps, as each map shows reasonable agreement with ground truth (i.e., bimodal for $CM_1$ and unimodal for $CM_2$, as expected via Fig. S13).

The NLSA outputs for $CM_1$ in $PD_{33}$ follow similar trends to those described for $PD_2$, with the exception that the overall range of motion for this CM is noticeably reduced compared to outputs from ESPER. For $CM_2$, matters are much worse. While our procedure using ESPER correctly charted a rotated, properly-aligned set of eigenfunctions, ManifoldEM employing NLSA used the existing manifold embedding without applying the essential eigenfunction realignment. As a result, the 2D movie produced by NLSA having closest resemblance to $CM_2$ (i.e., $\Psi_4$) demonstrated a physically-impossible sequence of motions: the splitting of the $CM_2$ domain into two separate domains. At the end of movie M8, the NLSA 2D movies obtained for the leading four eigenfunctions are shown for comparison. Here, both (1) a physically-impossible splitting of the $CM_1$ domain, and (2) a subdued $CM_2$ motion can be seen in the 2D movies obtained for both $\Psi_2$ and $\Psi_3$.

In the third scene of movie M8, the NLSA outputs of $PD_{49}$ can be described most similarly to those obtained in $PD_2$, and aside from flaws previously listed, are in general visual agreement with ESPER outputs. Again, the last segment of movie M8 showcases the alternative NLSA outputs obtained from $\Psi_3$ and $\Psi_4$, for which physically-impossible conformational information is apparent corresponding to $CM_1$ and $CM_2$ domains, respectively. Note that while NLSA and ESPER have been provided the exact same data—even up to generation of identical manifold embeddings—only ESPER is able to fully leverage the geometric structure present to consistently recapitulate ground-truth conformational motions and occupancies from a variety of PD manifolds. Further, while ESPER offers strategies to procedurally avoid introduction of nonsensical contextual output, NLSA can generate 2D movies with a wide range of defects[18], with each having the potential of appearing as a likely CM candidate to the naive eye.

Finally, we note the total computation time for performing these two techniques on the same CM-eigenvector ($\Psi_1$) from the same PD manifold ($PD_2$), with final output a single 2D movie (as seen in movie M8). The ESPER approach required approximately one minute for finding the optimal $d$-dimensional rotations and CM subspaces for this $\Omega_{\text{PD}}$, followed by approximately two minutes to perform subspace partitioning on the leading CM parabola. Meanwhile, the NLSA approach, which does not use our eigenfunction-rotation technique, took 4 hours and 37 minutes to furnish a 2D movie. The total computation time for NLSA was thus over 90 times longer than ESPER, with both methods having been run using a single-processor on the same workstation (3.8 GHz 8-Core Intel Core i7; 8 GB 2667 MHz DDR4). Although not investigated further for this study, we additionally note that in the current ManifoldEM framework[18], it is required that this time-expensive NLSA computation is repeated in its entirety for every $\Omega_{\text{PD}}$ eigenvector chosen during final compilation of the free-energy landscape. Meanwhile, using our intersection of image-indices approach, the ESPER algorithm compiles all PD-eigenfunction content and generates free-energy landscapes within minutes. Recall that these high computational demands were rationalized for NLSA as a way to handle the unknown manifold structures, noise-reduce images and normalize unknown rates of change. Meanwhile, as our heuristic analysis directly informed us of anticipated manifold characteristics and spectral structure, we were able to design ESPER to circumvent these previous unknowns, and perform the necessary operations required to accurately retrieve high-resolution images and corresponding occupancy map of all CM states. The results of our analysis show that ESPER produced appreciably more accurate outputs than the previous technique in a fraction of the time.

### SM-XX. STRUCTURAL VALIDATION OF ESPER OUTPUTS

In the following we provide a validation for the fidelity of the ESPER outputs as they relate to the ground-truth electron density maps obtained from PDB-formatted atomic-coordinate structures. (Recall that, following our synthetic-generation protocol, these EDMs are then projected to form images—and these images then duplicated via multiple occupancy assignments and modified by CTF and noise—before being processed via the ESPER framework). Specifically for our validation, we generated a Fourier Shell Correlation (FSC) curve[7] for the ground-truth simulated map against the corresponding ESPER recovered map. As seen in Fig. S30 for state 05_10, we found favorable global agreement between maps up to a resolution near 3 Å (i.e., the value used to generate each ground-truth EDM).

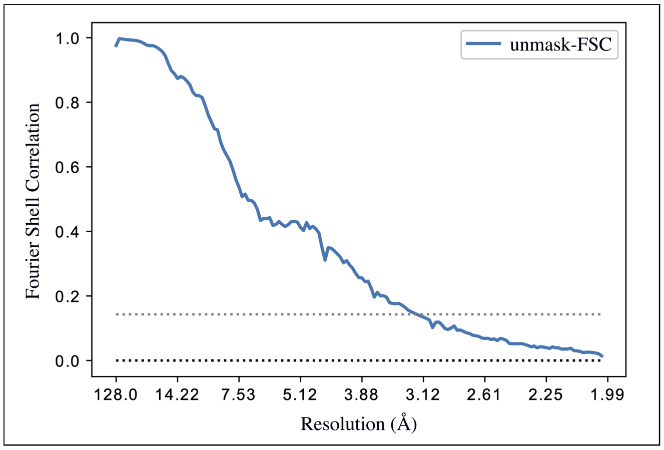


FIG. S30: FSC curve comparing the state 05_10 input (ground-truth) and output (ESPER) maps. Specifically, the FSC measures the normalized cross-correlation coefficient between the two maps as measured over a series of shells in Fourier space. As one proceeds along the $x$-axis from the left (representing the center of the FT) to the right, increasingly larger shells are compared in Fourier space, such that the largest shells (far right) correspond to the highest resolution features. The FSC curve thus provides a global measure of how well one 3D density map matches the other.

For molecules that exhibit domain motions, the global resolution is no longer a good indicator of how well-resolved these regions are in the reconstructed EDM. Thus, a complementary local validation was done by calculating the Q-scores[36] for the same example EDM (state 05_10) output from ESPER; with the use of Q-scores serving as a local indicator of how well our final 3D density maps recover the ground-truth atomic information. The results of this analysis are shown in Fig. S31.

For these statistics, the mean Q-score for the (1) backbone, (2) side chains, and (3) residues was 0.784, 0.768 and 0.778, with standard deviation 0.042, 0.042 and 0.039, respectively. Almost all Q-scores obtained were well above the expected Q-score value (0.5862), which is calculated[36] based on correlations to the reported resolutions of maps in the EMDB. On average, these Q-scores were approximately 1.3 times that of the expected value. We note that exceeding these expectations is anticipated, since our study was initiated with pristine structural information (i.e., atomic models). Outliers were found predominantly in the periphery of the domain corresponding to $CM_2$ and included side chains (minimum Q-score: 0.401) and three residues (minimum Q-score: 0.377). While the FSC provides an indicator for the global agreement between two maps, we believe the Q-score is more appropriate for interpretation of our outputs, especially given the design of our study; i.e., a synthetic-generation protocol from ground-truth atomic coordinates. Similarly favorable structural fidelity was found for the other outputs of ESPER in relation to their corresponding ground-truth atomic representations.

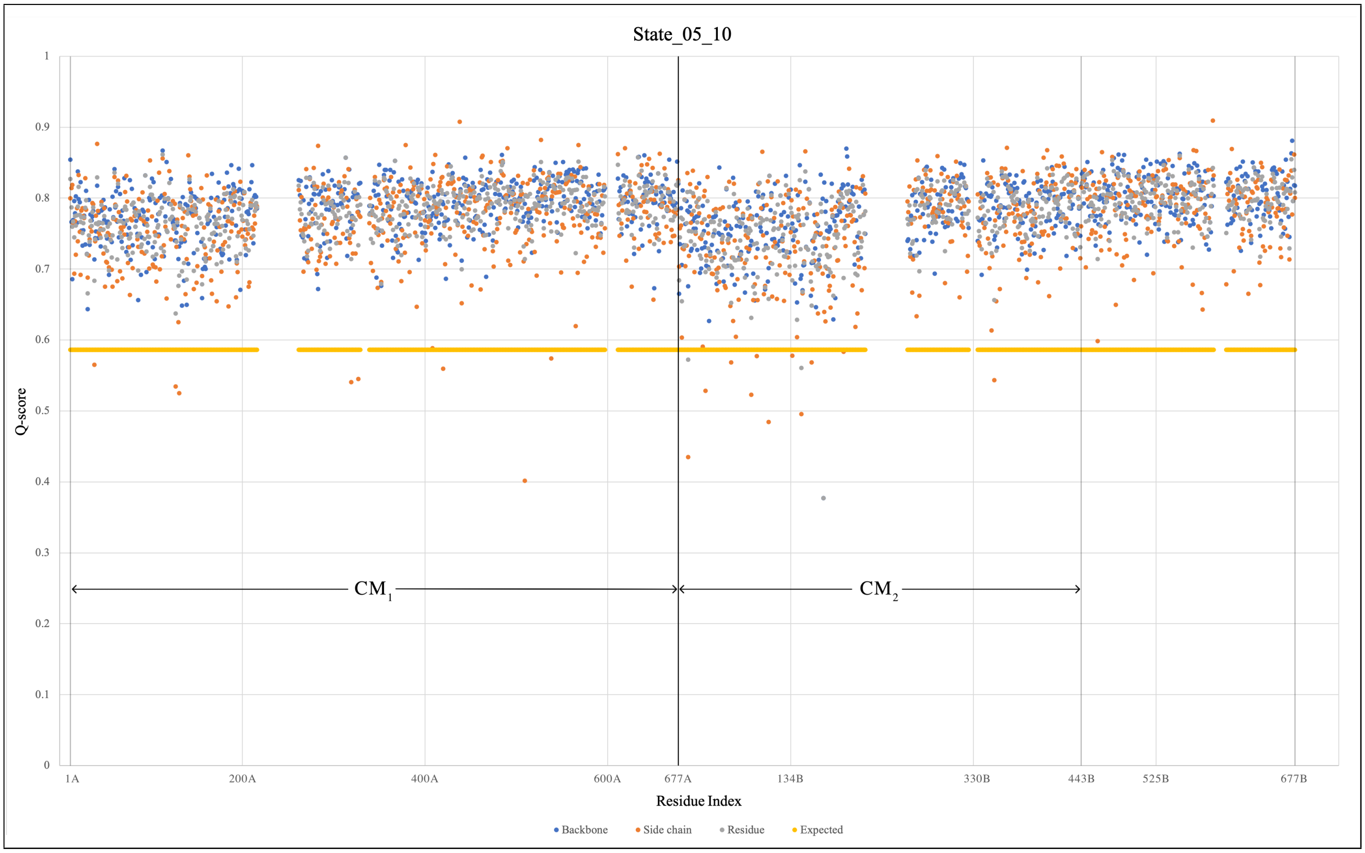


FIG. S31: Q-scores for protein backbone, side chains and residues calculated using ESPER output map 05_10 with corresponding ground-truth atomic-coordinate structure. Q-scores were ascertained using the MapQ[37] plugin for Chimera[6]. As can be seen, the range of residues corresponding to each conformational motion ($CM_1$ and $CM_2$) are demarcated on chain A and B, respectively. Empty Q-scores correspond to those residue indices missing in the initial crystal structure (PDB 2CG9), whether due to insufficient resolution or electron density in the preceding study. The expected Q-score represents the average Q-score at a resolution of 3 Å, and is calculated via MapQ based on the reported resolutions of 3D density maps in the Electron Microscopy Data Bank (EMDB).
